# A Comprehensive Survey of *C. elegans* Argonaute Proteins Reveals Organism-wide Gene Regulatory Networks and Functions

**DOI:** 10.1101/2022.08.08.502013

**Authors:** Uri Seroussi, Andrew Lugowski, Lina Wadi, Robert X. Lao, Alexandra R. Willis, Winnie Zhao, Adam E. Sundby, Amanda G. Charlesworth, Aaron W. Reinke, Julie M. Claycomb

**Affiliations:** University of Toronto

**Keywords:** Argonaute, miRNA, piRNA, 22G-RNA, 26G-RNA, RNAi, *C. elegans*, gene regulation, germline, germ granule, fertility, pathogen response

## Abstract

Argonaute (AGO) proteins associate with small RNAs to direct their effector function on complementary transcripts. The nematode *C. elegans* contains an expanded family of 19 functional AGO proteins, many of which have not been fully characterized. In this work we systematically analyzed every *C. elegans* AGO, using CRISPR-Cas9 genome editing to introduce GFP::3xFLAG tags. We have characterized the expression patterns of each AGO throughout development, identified small RNA binding complements, and determined the effects of *ago* loss on small RNA populations and developmental phenotypes. Our analysis indicates stratification of subsets of AGOs into distinct regulatory modules, and integration of our data led us to uncover novel stress-induced fertility and pathogen response phenotypes due to *ago* loss.

## Introduction

Small RNA-mediated gene regulatory pathways (collectively referred to as RNA interference, RNAi) have been identified in organisms from all domains of life (Swarts *et al*., 2014). These pathways utilize an array of molecular mechanisms in the epigenetic modulation of gene expression, and exert their influence on nearly every step in the lifecycle of a transcript, from transcription to translation (Meister, 2013; Wu *et al*., 2020). At the cellular level, small RNA (sRNA) pathways are key contributors to regulating genome and transcriptome homeostasis, both under normal conditions and stress responses. At the organismal level, sRNA pathways are key regulators of gene expression programs that direct development and differentiation, and mis-regulation of sRNA pathways or components can lead to conditions such as cancer and infertility (Wu *et al*., 2020).

The central effectors of sRNA pathways are the highly conserved Argonaute (AGO) family of proteins. AGOs are the core components of ribonucleoprotein complexes called RISCs (RNA Induced Silencing Complexes), and are guided in a sequence-specific manner by sRNAs (18-30 nucleotides long) to complementary target transcripts (Dueck and Meister, 2014). AGOs have a bi-lobed structure consisting of four major domains: PAZ, MID, PIWI, and a low complexity N-terminal domain. The PAZ and MID domains possess pockets to coordinate 3′ and 5′ end sRNA binding, respectively (Sheu-Gruttadauria and MacRae, 2017). The PIWI domain resembles RNaseH and has the capacity to direct endonucleolytic cleavage of the target RNA if the active site harbors a tetrad of catalytic amino acids (DEDD/H, (Nakanishi *et al*., 2012)). Relatively few AGOs possess this catalytic tetrad, and many AGOs recruit additional proteins to elicit other gene regulatory outcomes, such as mRNA de-capping, de-adenylation, or chromatin modulation.

In *C. elegans*, at least four types of endogenous sRNAs—miRNAs, piRNAs, 22G-RNAs and 26G-RNAs—and as many as 27 AGO-like genes have the potential to contribute to complex networks of gene regulation in different tissues throughout development. miRNAs and piRNAs are genomically encoded and transcribed by RNA Polymerase II, while the 22G-RNAs and 26G-RNAs are generated by the activity of different RNA Dependent RNA Polymerases (RdRPs). miRNAs are known to associate and function with the conserved AGOs ALG-1, ALG-2 and ALG-5 (Brown *et al*., 2017). The Piwi-interacting RNAs (piRNAs, also called 21U-RNAs in *C. elegans*) bind to the PIWI AGO PRG-1 and are thought to maintain germline genome integrity by silencing foreign or deleterious nucleic acids such as transgenes (Lee *et al*., 2012; Shirayama *et al*., 2012). Although piRNA pathways in other animals play a more prominent role in regulating transposable elements than in *C. elegans*, the functions of the piRNA pathway are broadly and consistently required in animal germlines to ensure fertility (Ozata *et al*., 2019).

Two additional types of endogenous sRNAs present in *C. elegans* are the 26G-RNAs and 22G-RNAs (named for their predominant length and 5′ nucleotide). Because these sRNAs are generated by RdRPs, they are thought to exploit perfect complementarity to their targets. The 26G-RNAs are synthesized by the RdRP RRF-3, which generates dsRNA that is processed into 26G-RNAs by the endonuclease DICER and the phosphatase PIR-1(Chaves *et al*., 2021). 26G-RNAs are classified into two groups: those of spermatogenic origin (class I, associated with ALG-3 and ALG-4, (Han *et al*., 2009; Conine *et al*., 2010)) and those of oogenic and embryonic origin (class II, associated with ERGO-1,(Han *et al*., 2009; Vasale *et al*., 2010)).

The 22G-RNAs are generated by the RdRPs RRF-1 and EGO-1, independent of DICER. Currently, 22G-RNAs are divided into two main groups, those that are bound by CSR-1 and target germline expressed protein coding transcripts to protect them from silencing (Claycomb *et al*., 2009; Wedeles, Wu and Claycomb, 2013), and those that are bound by other WAGO class AGOs (such as WAGO-1 and HRDE-1) that silence protein-coding genes, pseudogenes, transposable elements, and cryptic loci (Gu *et al*., 2009). 22G-RNAs are generally thought to act as secondary, amplified sRNAs that are synthesized after a transcript is targeted by a primary sRNA/AGO complex, with the main exception being the majority of CSR-1 associated 22G-RNAs. Primary sRNAs take several forms: piRNAs (PRG-1), 26G-RNAs (ALG-3, ALG-4 and ERGO-1), and exogenous-siRNAs produced by DICER during exogenous RNAi (exoRNAi) (RDE-1).

*C. elegans* AGOs have generally been studied on a case-by-case basis, with *agos* being uncovered via genetic screens, or selected for study based on phenotype (e.g., Tabara *et al*., 1999). Such approaches are limited because not all phenotypes to which *agos* may contribute have been tested, and redundancy among the *agos* could confound their recovery in genetic screens. To date, only one study has taken a systematic approach to understanding AGO functional relationships, examining the requirement for each *ago* in exogenous RNAi (Yigit *et al*., 2006). A lack of antibodies against individual AGOs and difficulties with transgenic approaches have also hampered the development of a cohesive set of reagents to study AGO function. Indeed, several *C. elegans* AGOs have yet to be studied, and others remain only partially characterized.

In this study, we have undertaken a systematic analysis of the *C. elegans* AGOs. We employed CRISPR-Cas9 genome editing to introduce GFP::3xFLAG epitope tags in the endogenous loci of each *ago*, using these strains to examine spatiotemporal expression profiles throughout development, combined with sequencing sRNAs from AGO complexes and *ago* mutants to define the core of *C. elegans* sRNA pathways. We systematically assessed fertility of *ago* mutants and employed phenotypic assays directed by our expression and sRNA sequencing data, enabling us to uncover new roles for specific AGOs in maintaining germline integrity and in regulating immune responses to bacterial and viral pathogens. Collectively, our findings provide a foundation for understanding the full scope of sRNA pathway activity in *C. elegans*. With these AGO tools and knowledge of sRNA binding partners and targets, our findings provide a deeper understanding of sRNA functions throughout development and under varied environmental conditions in *C. elegans*.

## Results

### Systematic analysis of *C. elegans* Argonautes

Previous studies identified 27 *ago* genes in *C. elegans* (Yigit *et al*., 2006), however some have been reclassified as pseudogenes. To define an updated set of *ago* genes to study, we searched the genome (WormBase version WS262) for genes that contain PAZ and PIWI domains, have a predicted protein size of ∼100 kDa, and bear homology to known AGOs. Twenty one genes met these criteria and we ultimately characterized 19 of these AGOs (Table S1, Fig 1A-B). Construction of a phylogenetic tree for these 19 AGOs in relation to *Arabidopsis thaliana* AGO1 and *Drosophila melanogaster* PIWI places seven AGOs in the AGO clade: ALG-1, ALG-2, ALG-3, ALG-4, ALG-5, ERGO-1 and RDE-1; a single AGO in the PIWI clade: PRG-1; and 13 AGOs in the WAGO clade: CSR-1, C04F12.1 (renamed VSRA-1 for Versatile Small RNAs Argonaute-1, see below), WAGO-1, PPW-2, WAGO-4, SAGO-2, PPW-1, SAGO-1, HRDE-1, WAGO-10 and NRDE-3 (Fig 1A).

**Figure 1.**
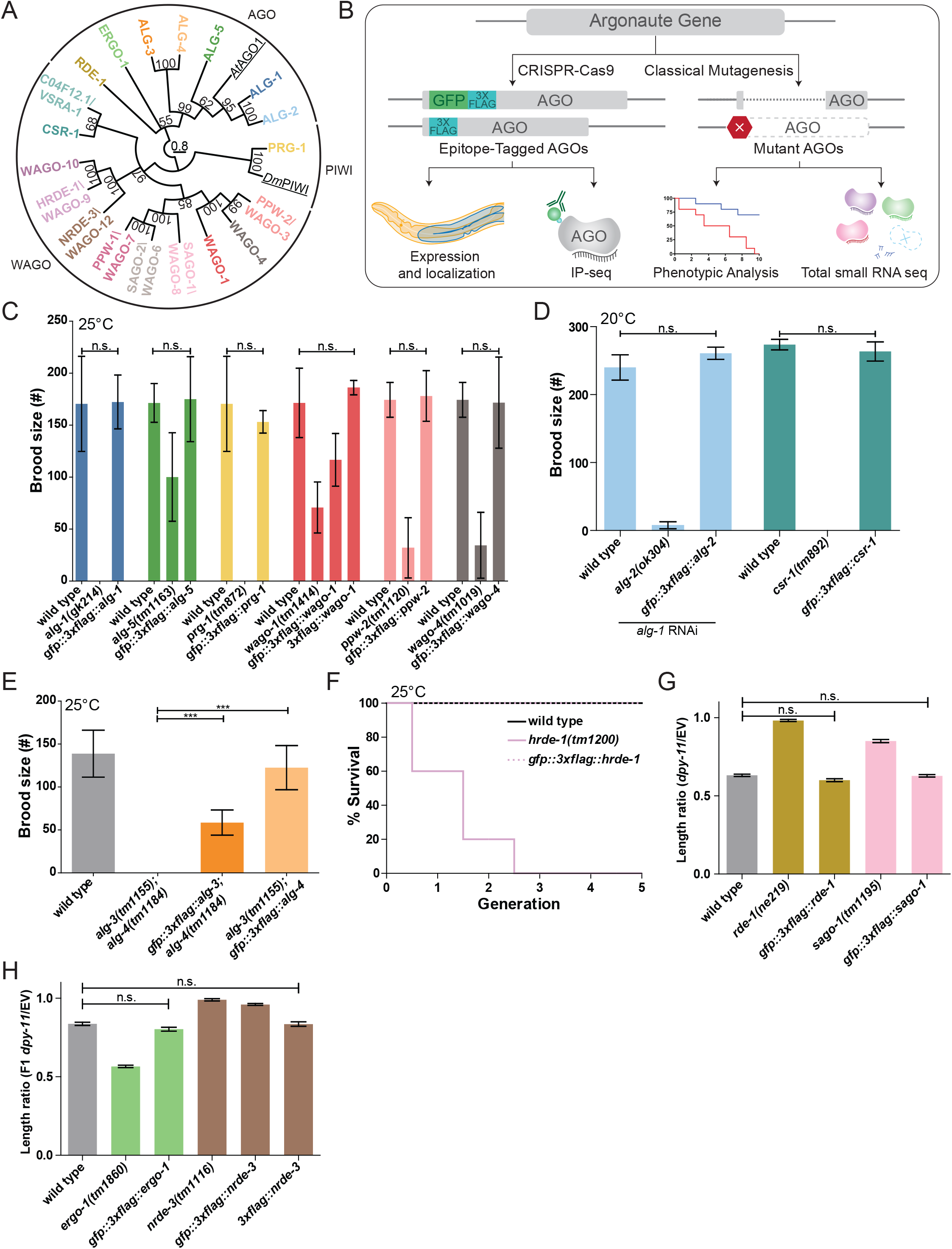
Functional validations of GFP::3xFLAG and 3xFLAG tagged Argonautes. (A) Maximum likelihood evolutionary tree of *A. thaliana* AGO1 (AtAGO1), *D. melanogaster* PIWI (DmPIWI) and *C. elegans* Argonautes. (B) Workflow for characterizing *C. elegans* Argonautes. (C) Functional validations of tagged ALG-1, ALG-5, PRG-1, WAGO-1, PPW-2 and WAGO-4 strains. Brood size was determined at 25°C for each indicated genotype. N ≧ 5 worms per condition. (D) Functional validations of tagged ALG-2 and CSR-1 strains. Brood size was determined at 20°C for each indicated genotype. For the ALG-2 tag validations, the brood size was determined when worms were fed dsRNA of *alg-1*. N = 5 worms per condition. (E) Functional validations of tagged ALG-3 and ALG-4 strains. Brood size was determined at 25°C for each indicated genotype. N = 10 worms per condition. (F) Functional validation of the tagged HRDE-1 strain. A mortal germline assay was conducted at 25°C. N = 5 worms per condition. (G) Functional validations of tagged RDE-1 and SAGO-1 strains. Worms were fed bacteria expressing dsRNA of *dpy-11* or an empty vector (EV) RNAi control. The length ratio of *dpy-11* dsRNA fed P0 worms compared to the average length on EV was determined. N = 30 worms per condition. (H) Functional validations of tagged ERGO-1 and NRDE-3 strains. Worms were fed bacteria expressing dsRNA of *dpy-11* or an empty vector (EV) control. The length ratio of the F1s of the *dpy-11* dsRNA fed worms compared to the average length on EV was determined. N = 30 worms per condition. (C-H) * = *p*-value < 0.05, ** = *p*-value < 0.01, *** = *p*-value, n.s. = not significant. One way ANOVA with Tukey Post Hoc multiple comparison test.

We used CRISPR-Cas9 genome editing to introduce a GFP::3xFLAG tag into the N-terminus or within the first exon of the endogenous gene loci of all 21 *agos* (Dickinson et al., 2015) (Fig S1A). We detected GFP expression for both the transcriptional reporter and GFP::3xFLAG::AGOs for 19 AGOs by confocal microscopy. We verified that the GFP::3xFLAG::AGOs were full-length proteins by western blot analysis (Fig S1B). We were unable to detect WAGO-5 and WAGO-11 fusion protein expression by microscopy or western blot analysis (Fig S2A). RNA-tiling-array data (modENCODE Consortium *et al*., 2009) showed that both *wagos* are expressed at low levels with poor sequencing coverage (Fig S2B), and RT-PCR only detected low amounts of *wago-11* mRNA (Fig S2C). *wago-5* is targeted by WAGO-1-associated 22G-RNAs, suggesting it is silenced by the WAGO pathway (Fig S2D). During this project, the designation of *wago-11* was changed to “pseudogene” in WormBase (version WS275). Our data support that these *wagos* are pseudogenes, therefore we excluded these WAGOs from further analysis.

We tested the function of the tagged AGOs by assessing phenotypes associated with *ago* loss of function (Yigit *et al*., 2006; Batista *et al*., 2008; Guang *et al*., 2008; Han *et al*., 2009; Buckley *et al*., 2012; Brown *et al*., 2017; Xu *et al*., 2018) (Fig 1C-H, Fig 6D-E). While previous studies have defined loss of function defects for several *agos*, some *agos* had no known loss of function defects (Table S1). Of all the AGOs tested, the GFP::3xFLAG tagged AGOs WAGO-1 and NRDE-3 did not behave as wild-type in phenotypic assays (Fig 1C, H). Therefore, we tagged WAGO-1 and NRDE-3 with only 3xFLAG (Fig S2E) for the purpose of small RNA cloning and used the GFP::3xFLAG tagged strain for analysis of expression patterns. Both 3xFLAG tagged AGOs were functional in phenotypic assays (Fig 1C,H). The tagged C04F12.1/VSRA-1 and WAGO-10 were not tested for a specific phenotype as there are no known phenotypes for *C04F12.1/vsra-1* and *wago-10* mutants. As we observed that the GFP tag may interfere with function in some instances (e.g. WAGO-1 and NRDE-3), we tagged C04F12.1/VSRA-1 and WAGO-10 with 3xFLAG at the same position as the GFP::3xFLAG tags for the purpose of sRNA cloning out of an abundance of caution (Fig S2E).

### AGO sequencing reveals sRNA association

The sRNAs associated with each AGO provide important insight into the transcripts the AGOs may regulate. To identify the sRNAs that interact with each AGO, we performed immunoprecipitation (IP) followed by high-throughput sequencing of sRNAs for each of the tagged AGOs in duplicate. In parallel, we sequenced total sRNAs from the same lysates as the IPs (“Input” samples). We conducted IPs on worm populations at the L4 to young adult transition (58h post L1 synchronization), because all but a few AGOs are expressed at this stage. The only exceptions were ALG-3, ALG-4 and WAGO-10, which are only expressed during spermatogenesis (Charlesworth *et al*., 2021). Therefore, we conducted ALG-3, ALG-4, and WAGO-10 IPs during the mid L4 stage (48h post L1 synchronization), during which spermatogenesis occurs. For consistency, we treated all libraries with 5′ polyphosphatase to enable detection of 5′ tri-phosphorylated small RNA species (22G-RNAs), along with 5′ mono-phosphorylated species (miRNAs, piRNAs, 26G-RNAs) (Table S2).

We assessed the length and 5′ nucleotide distribution of sRNAs associated with each AGO and mapped this total set of reads to genomic features (Fig 2A). We also determined how many miRNAs, piRNAs, and other genomic features (protein coding genes, transposable elements, pseudogenes, and long intergenic noncoding RNAs, or lincRNAs that are targeted by antisense endogenous siRNAs including, but not limited to, 22G-RNAs and 26G-RNAs) were *<u>enriched</u>* over two-fold in both IP replicates relative to the input samples (Table S3). For this enrichment analysis, we did not place any constraints on sRNA length or 5′ nucleotide, and considered all genome mapping antisense reads. We defined 22G-RNA reads as 20-24nt with no 5′ nucleotide bias, and 26G-RNAs as 25-27nt with no 5′ nucleotide bias. These stringent criteria led to the assignment of a high confidence set of AGO-enriched sRNAs. We refer to transcripts for which antisense siRNAs are enriched over two-fold as the “targets” of AGO/sRNA complexes.

**Figure 2.**
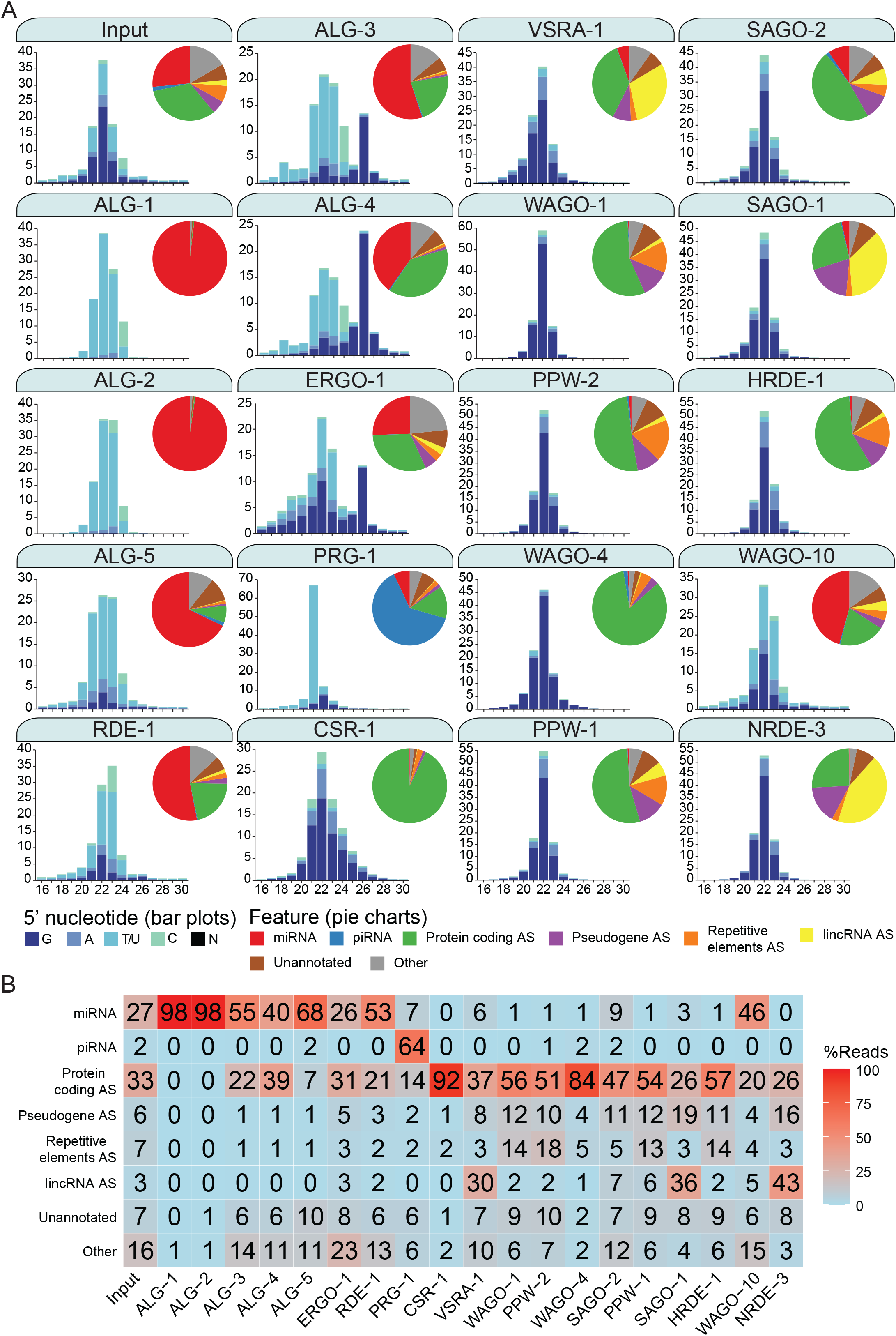
Argonautes associate with different types of sRNAs and target different categories of genetic features. (A) 5′ nucleotide and length of sRNAs present in each Argonaute IP shown in bar graph form. The pie charts depict which type of genetic element (biotype) the sRNAs correspond to, as listed. AS = antisense, S= sense. The “Other” category encompasses: miRNA AS, piRNA AS, protein coding S, pseudogene S, repetitive elements S, lincRNA S, rRNA S/AS, snoRNA S/AS, snRNA S/AS, tRNA S/AS, ncRNA S/AS, and antisense noncoding RNAs (ANRs) (Nam and Bartel, 2012). The average of two biological replicates is shown. The GFP::3xFLAG tagged Argonautes were used for IPs except for C04F12.1/VSRA-1, WAGO-10 and NRDE-3, where a 3xFLAG tag was used. All IPs were performed on Young Adult samples except for ALG-3, ALG-4, and WAGO-10 which were conducted on L4 staged animals. The CSR-1 strain tags both isoforms. (B) A table summarizing the percentage of reads in each set of AGO IPs corresponding to genetic element types in (A).

A subset of the AGO clade primarily associated with miRNAs (Correa *et al*., 2010; Brown *et al*., 2017; Svendsen *et al*., 2019). For ALG-1, miRNAs comprised ∼98% of all associated reads (47 miRNAs enriched); ALG-2, ∼98% (81 enriched), ALG-5, ∼68% (37 enriched); and RDE-1, ∼53% (103 enriched). RDE-1 is involved in exoRNAi (Tabara *et al*., 1999), however we found that it also associates with 22G-RNAs targeting protein coding genes (∼21%, 536 enriched genes). We expect that these 22G-RNAs possess a 5′ mono-phosphate, given that RDE-1 was previously shown to associate with multiple types of sRNAs that are likely DICER products (Correa *et al*., 2010). We also detected abundant miRNAs associated with the rest of the AGO clade AGOs ALG-3, ALG-4 and ERGO-1, with ∼55% (26 enriched), ∼40% (2 enriched) and ∼26% (33 enriched) of reads corresponding to miRNAs, respectively. These three AGOs were previously described as genetically required for 26G-RNA accumulation, and ERGO-1 was shown to physically associate with 26G-RNAs (Conine *et al*., 2010; Vasale *et al*., 2010). Indeed, ALG-3, ALG-4, and ERGO-1 all associate with 26G-RNAs with ∼17%, ∼32% and ∼15% of reads corresponding to 26G-RNAs respectively (Fig 2). The endo-siRNAs in ALG-3 and ALG-4 IPs are primarily antisense to protein-coding genes (∼22%, 2561 enriched and ∼39%, 2848 enriched respectively) and pseudogenes (∼1%, 66 enriched and 1.2%, 97 enriched respectively), while ERGO-1 targets primarily protein-coding genes (∼31%, 411 enriched), pseudogenes (∼5%, 39 enriched) and lincRNAs (∼3%, 30 enriched) (Fig 2).

PRG-1, the only PIWI homolog in *C. elegans*, PRG-1, is known to associate with, and be required for piRNA stability (Batista *et al*., 2008). It is the major piRNA-associated AGO, and ∼64% of the reads in PRG-1 IPs correspond to piRNAs (5932 enriched), (Fig 2). We also detected 22G-RNAs (∼21%) that primarily-targeted protein coding genes enriched in PRG-1 complexes (150 gene targets enriched).

Previous studies predicted that all WAGOs would associate with 22G-RNAs (Guang *et al*., 2008; Gu *et al*., 2009; Buckley *et al*., 2012; Xu *et al*., 2018), and we observed that 22G-RNAs are the most abundant class of small RNAs associated with the WAGOs, many of which are antisense to protein-coding genes, ranging from ∼26% of reads in SAGO-1 IPs to ∼93% of reads in CSR-1 IPs. Groups of WAGOs associate more prominently with sRNAs that target specific genomic features including pseudogenes, lincRNAs, and repetitive and transposable elements. Pseudogenes are primarily targeted by HRDE-1 (∼19% of reads in the IP are antisense to these elements), NRDE-3 (∼17%), WAGO-1 (∼12%), PPW-1 (∼12%), SAGO-2 (∼11%), SAGO-1 (∼11%), and PPW-2 (∼10%) (Fig 2). lincRNAs are primarily targeted by NRDE-3 (∼43% of reads in the IP are antisense to these elements), SAGO-1 (∼35%), C04F12.1/VSRA-1 (∼30%), SAGO-2 (∼7%), and PPW-1 (∼6%) (Fig 2). sRNAs antisense to repetitive and transposable elements are most abundant in PPW-2 (∼19% of reads in the IP are antisense to these elements), WAGO-1 (∼18%), HRDE-1 (∼15%) and PPW-1 (∼14%) complexes (Fig 2).

### miRNAs

#### miRNAs associate with ALG-1, ALG-2, ALG-5, and RDE-1

437 miRNAs are annotated by mirBase 22.1 (encompassing 253 families with individual 5p or 3p strands). We detected reads for 402 miRNAs across our AGO Input and IP samples. Of these, 190 were found to be enriched over two-fold in the AGO IPs. We observed miRNAs in association with the known miRNA binding AGOs, ALG-1, ALG-2, ALG-5, and RDE-1, and in association with the 26G-RNA binding AGOs ALG-3, ALG-4, and ERGO-1 (Fig 3A). Of the miRNAs enriched in the AGO IPs, some were enriched in association with only one AGO while others were enriched in multiple AGOs (Fig 3A). For example, 103 miRNAs were enriched in RDE-1 complexes, with 55 being exclusive to RDE-1. RDE-1 was the only AGO where the majority (63/103) of enriched miRNAs were not conserved with those of the related nematode, *C. briggsae*, suggesting that newly-evolved miRNAs may be routed initially into the RDE-1 pathway before subsequently integrating into the ALG-1/2/5 pathway (Fig 3L).

**Figure 3.**
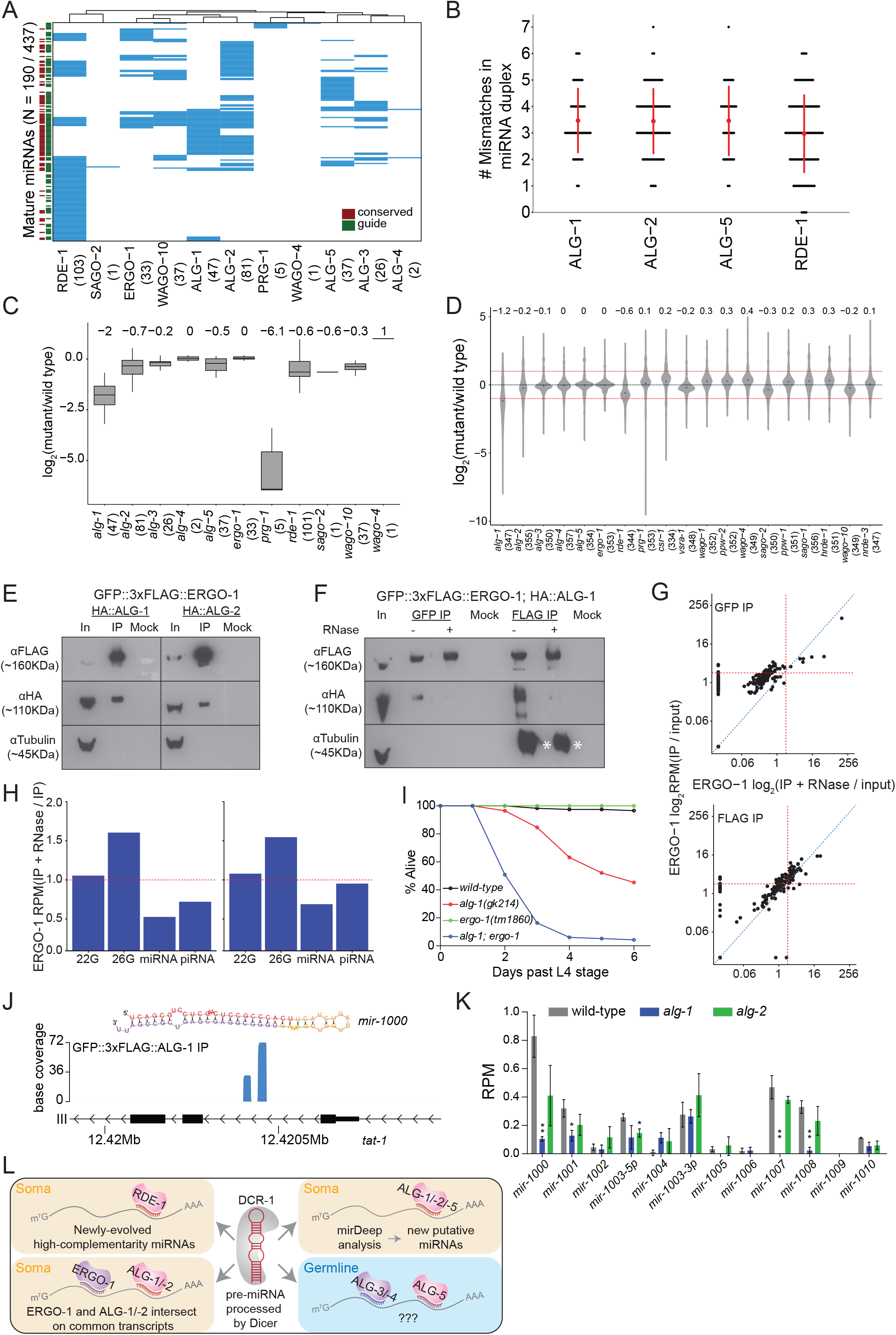
Analysis of miRNAs in AGO IPs and *ago* mutants reveals novel miRNAs. (A) Clustering diagram of miRNAs enriched in AGOs. Each blue line represents an individual mature miRNA sequence. (B) The number of mismatches in the precursor miRNA sequences for which a mature miRNA was enriched in the indicated AGOs. Each black dot represents a single precursor miRNA duplex. The red dot and lines indicate the average and standard deviation. (C) Fold change of miRNAs enriched in AGO IPs in the corresponding *ago* mutant. (D) Fold change of all detected miRNAs in *ago* mutants as compared to wild-type. (E) Western blots of co-IP experiments of ERGO-1 and ALG-1 and ALG-2. GFP::3xFLAG::ERGO-1 was crossed to HA::ALG-1 or HA::ALG-2 strains and IPed using anti-GFP antibodies. (F) As in (E) but GFP::3xFLAG::ERGO-1 IPs were conducted either with anti-GFP or anti-FLAG antibodies with or without RNase treatment. Asterisks indicate IgG. (G) Scatter plots showing enrichment of miRNAs in IP and IP+RNase treated samples of GFP::3xFLAG::ERGO-1. (H) Bar plots showing quantification of sRNA types in IP and IP+RNase treated samples of GFP::3xFLAG::ERGO-1. (I) Survival of worms of the indicated genotype beyond the L4 stage (bottom). N ≧ 100. (J) An example of a novel miRNA within an intron of the gene *tat-1* as determined by mirDeep2 analysis of ALG-1 IPs. (K) Analysis of the levels of predicted novel miRNAs in wild-type, *alg-1* and *alg-2* mutants. Predicted novel miRNAs are provisionally named. (L) A summary of miRNA pathway observations.

Adjusting the number and position of mismatches in the precursor miRNA (pre-miRNA) duplex of a transgenic *let-7* miRNA has been shown to shift the balance between ALG-1 and RDE-1 loading of the transgenic *let-7*. These experiments demonstrated that ALG-1 preferentially associates with miRNAs from mismatched precursors while RDE-1 prefers perfectly matching precursors (Steiner *et al*., 2007). Therefore, we examined the number of mismatches in precursor miRNA duplexes that are enriched in ALG-1, ALG-2, ALG-5 and RDE-1 complexes (Fig 3B). miRNAs loaded into RDE-1 showed a lower average of mismatches in their precursors, and miRNAs derived from precursors with no mismatches were only bound by RDE-1, although some miRNAs derived from mismatched precursors were also loaded into RDE-1. These data suggest that endogenous miRNAs with higher complementarity in their precursor duplex are preferentially loaded into RDE-1.

Given that ALG-1 and ALG-2 are required for the stability of miRNAs (Grishok *et al*., 2001; Brown *et al*., 2017), and sRNAs are generally unstable in the absence of their AGO binding partner, we asked whether these AGOs are required for the stability of their enriched miRNAs or for miRNAs in general, by sequencing sRNAs from *ago* mutants and wild-type worms. We found that the AGO-enriched miRNAs were substantially depleted in *alg-1*, *alg-2*, *alg-5*, and *rde-1* mutants (Fig 3C), and five PRG-1 enriched “miRNAs” were over 60 times lower in abundance in *prg-1* mutants (see below). Loss of ALG-1 had the most substantial effect on global miRNA levels, leading to a greater than two-fold decrease (Fig 3D), indicating that *alg-1* is genetically required for the stability of most miRNAs and potentially explaining why *alg-1* mutants have more severe phenotypes than the other miRNA-associated *ago* mutants (Bukhari *et al*., 2012; Brown *et al*., 2017). Loss of RDE-1 also led to a substantial depletion of miRNAs overall, while loss of ALG-2 or ALG-5 did not result in major changes, which could reflect redundancy or differences in function for these AGOs. Loss of several WAGOs also led to a decrease in global miRNA levels, and we speculate that this is due to indirect effects on protein coding gene regulation, rather than a direct influence on the miRNA pathway.

#### miRNA and ERGO-1 26G-RNA pathways intersect

To understand the association of ALG-3, ALG-4, and ERGO-1 with both 26G-RNAs and miRNAs, we tested whether the GFP::3xFLAG tag interfered with proper sRNA loading, using existing ERGO-1 IP-sRNA sequencing data performed using an ERGO-1-specific antibody (Vasale *et al*., 2010). In this study, the authors detected a subset of ERGO-1 associated miRNAs, but dismissed this as a non-specific interaction. We re-analyzed these data using our custom computational pipeline and identified 26 miRNAs that were enriched two-fold over input and significantly overlapped with our IP dataset,suggesting that miRNA association with ERGO-1 is not a result of the GFP::3xFLAG tag, but that miRNA enrichment may be a property of ERGO-1 IPs.

We examined our AGO expression and localization data, and observed that ERGO-1 expression closely overlaps with ALG-1 and ALG-2, indicating that these AGOs could physically interact *in vivo* (Fig 6A, 6F, 7E, Fig S6-13). To determine whether ERGO-1 physically interacts with one or more of the miRNA-binding AGOs, we crossed GFP::3xFLAG::ERGO-1 worms to HA::ALG-1 and HA::ALG-2 tagged strains (Brown *et al*., 2017) and performed co-IP experiments. We found that ERGO-1 physically interacted with both ALG-1 and ALG-2 (Fig 3E) in an RNA-dependent manner (Fig 3F). These data suggest that the ERGO-1/ALG-1 or -2 interactions are due to AGO associations on shared target transcripts. Further supporting this model, we sequenced sRNAs associated with ERGO-1 IPs after RNAse treatment, and observed that miRNAs were substantially reduced, while 26G-RNAs were enriched, compared to non-RNAse treated ERGO-1 IPs (Fig 3G-H). These observations strongly suggest that the miRNA enrichment observed in the ERGO-1 IPs may be indirect, and due to an interaction between ERGO-1 and ALG-1 or ALG-2 on target transcripts, and implies that co-regulation of target transcripts by 26G-RNAs and miRNAs could occur.

To explore the functional and developmental consequences of physical interaction between ERGO-1 and ALG-1, we created *alg-1; ergo-1* double mutants. Loss of *alg-1* results in heterochronic phenotypes, however, loss of *ergo-1* has no obvious phenotypic impact, aside from enhanced ability to perform exogenous RNAi (Eri phenotype). *alg-1; ergo-1* double mutants appeared sickly, being smaller and more pale than wild-type animals, with many dying prematurely, and only ∼4% of the animals surviving past the L4 stage as compared to ∼45% and 100% for *alg-1* and *ergo-1* single mutants, respectively (Fig 3I). These data indicate that *alg-1* and *ergo-1* genetically interact to ensure survival into adulthood, and are consistent with the idea that coordinated regulation of targets by these AGOs is required for development.

#### Novel miRNAs associated with ALG-1/2

The sequencing depth of our IP experiments allowed us to identify novel and lowly-expressed miRNAs that may have previously eluded detection, been mis-annotated or otherwise not appreciated as bona fide miRNAs because they are not known to associate with a classical (miRNA) AGO. We used the miRNA prediction program mirDeep2 to analyze sequencing data from the miRNA binding AGO IPs (Friedländer *et al*., 2012). We found 10 putative, high-confidence miRNAs (Table S4) present in ALG-1 and ALG-2 complexes. These putative miRNAs are present at very low levels, <1 RPM on average in total sRNA samples (for comparison, the abundant miRNA let-7 is present at ∼6000 RPM levels) (Fig 3J). Consistent with these being bona fide miRNAs, five of the 10 putative miRNAs were significantly depleted in *alg-1* or *alg-2* mutants. Collectively, our results indicate these sRNAs are genuine miRNAs, as they bind to classical miRNA AGOs and rely on these AGOs for their stability (Fig 3K-3L).

### piRNAs

#### “miRNAs” associated with PRG-1 are mis-annotated piRNAs

We found three likely mis-annotated miRNAs (cel-miR-4936, cel-miR-8198-3p, and cel-miR-8202-5p) enriched in IPs of the Piwi AGO, PRG-1, consistent with a previous report of two similarly mis-annotated miRNAs(cel-miR-78 & cel-miR-798) (Batista *et al*., 2008). These miRNAs are 21nt long and have a 5′ uridine (Fig 4A), however four of five were not enriched in association with the miRNA binding AGOs (cel-miR-4936,cel-miR-798, cel-miR-8198-3p, and cel-miR-8202-5p). All five miRNAs were depleted over 60-fold on average in *prg-1* mutants, were only two-fold depleted on average in *rde-1* mutants, and were not depleted in canonical miRNA binding *ago* mutants *alg-1*, *alg-2*, and *alg-5* (Fig 3C, Table S6). These putative piRNA loci possess upstream regulatory sequences that fully or partially resemble Ruby motifs, found at most piRNA loci (Ruby *et al*., 2006) (Fig 4A). Thus, these five sRNAs are piRNAs that were mis-annotated as miRNAs, demonstrating the utility of AGO IP-sequencing data in annotating sRNA features in the genome.

**Figure 4.**
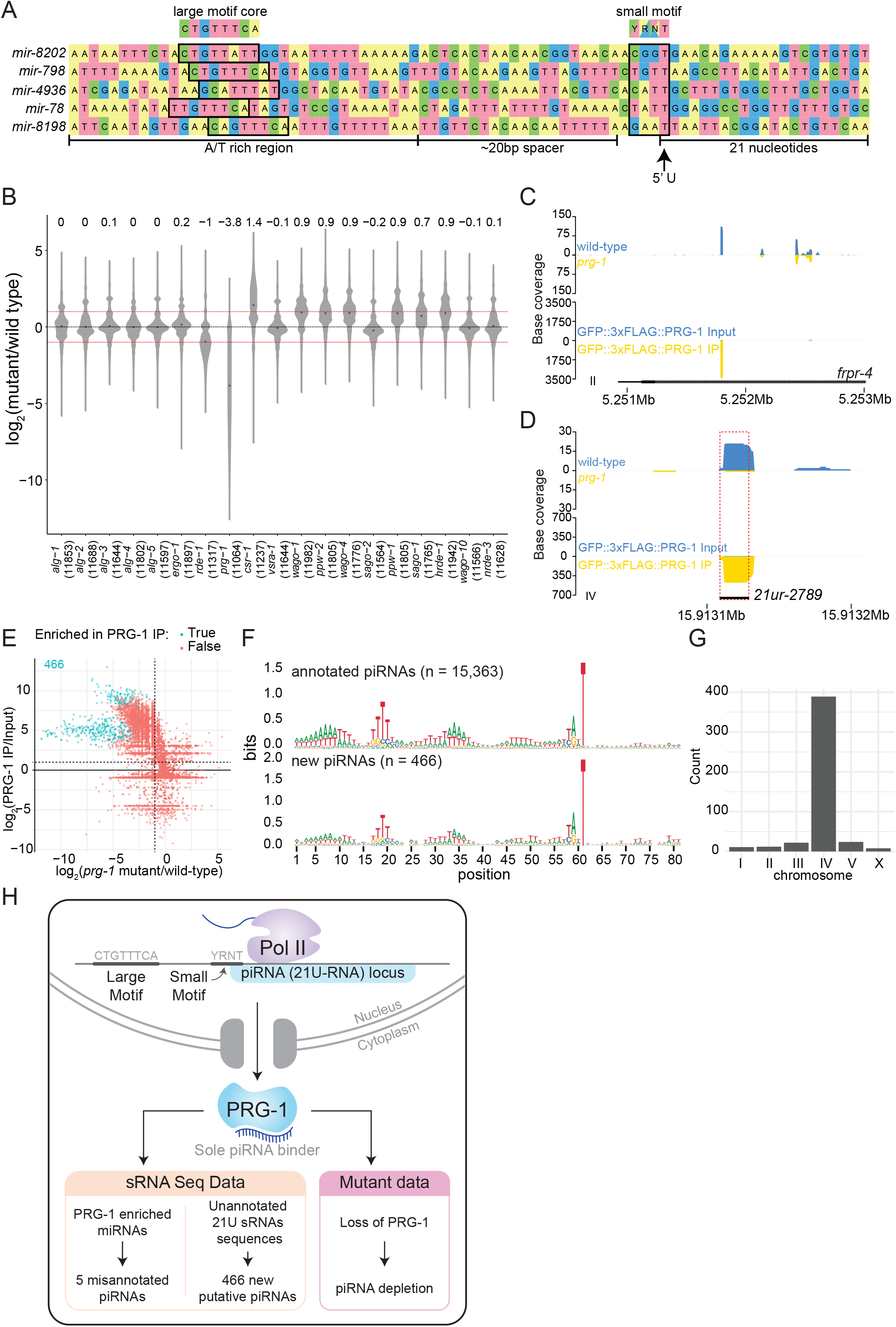
Analysis of piRNAs in AGO IPs and *ago* mutants reveals new piRNAs. (A) Genomic loci of annotated miRNAs that are enriched in PRG-1 IPs and suspected to be piRNAs. Ruby motifs highlighted above. (B) Violin plots depicting the fold change of all detected piRNAs in *ago* mutants as compared to wild-type. (C) An example of a novel piRNA sequence, where a 21U sRNA sequence that was enriched in PRG-1 IPs and depleted in *prg-1* mutants originated from the intron of the gene *frpr-4* in the antisense orientation. Note the difference in scales. (D) An example of a novel piRNA sequence originating from a shift of 3 nucleotides from the annotated *21ur-2789* piRNA (dotted red box, black line). Note the difference in scales. (E) Scatter plot showing individual expression levels of 21U sRNAs that are not annotated as piRNAs. The y-axis shows enrichment in PRG-1 IPs and the x-axis shows depletion in *prg-1* mutants. The cyan dots represent individual 21U sRNAs which are 2-fold enriched in PRG-1 IPs. (F) Sequence logo analysis of annotated piRNA loci (top) and the new 466 piRNA loci (bottom). (G) Chromosome distribution of the 466 putative piRNA sequences. (H) A summary of piRNA pathway observations.

#### piRNAs associate with the Argonaute PRG-1

Across all sRNA datasets, we detected reads for 14,568 out of the 15,363 annotated piRNAs in the genome. Among all of the IP data sets, we found 5,943 piRNAs enriched over two-fold, with Piwi PRG-1 being associated with nearly every enriched piRNA (5,932/5,943). The levels of piRNAs in *ago* mutants revealed that *prg-1* is required to maintain piRNA pools globally, with a ∼13.9 fold reduction in piRNA levels overall (Fig 4B), consistent with previous observations (Batista *et al*., 2008; Wang and Reinke, 2008).

We also observed that sRNAs from annotated genomic features other than piRNAs were enriched in the PRG-1 IPs. Most of these reads were 21 nucleotides long, possessed a 5′ uridine, and were depleted in *prg-1* mutants (Fig 4C). In some instances, 21U sRNA sequences partially overlapped annotated piRNAs; for example, a 21U sRNA sequence was shifted by three nucleotides from the 5′ end of the piRNA *21ur-2789* (Fig 4D). In total we found 466 sequences that were 21U, enriched over two-fold in both PRG-1 IP replicates, and were not perfectly matching to annotated piRNA sequences. All of these 21U sRNAs were depleted, and 222 were significantly depleted, in *prg-1* mutants compared to wild-type (Fig 4E, Table S5). Sequence logo analysis of the sequence upstream of the loci generating 21U sRNAs demonstrates similarity to the Ruby motifs of piRNAs (Fig 4F), and most of these sequences (375/466) originated from a previously described piRNA cluster which spanned coordinates 4.5-7.0M (Ruby *et al*., 2006) (Fig 4G). We conclude that these 466 21U sRNAs are previously uncharacterized piRNAs (Fig 4H).

### Endo-siRNAs: 22G-RNAs and 26-RNAs

#### Endo-siRNA associated AGOs cluster into four groups

We examined the AGOs that interact with 22G- and 26G-RNAs. Because these sRNAs are generated by RdRPs, we can predict their targets based on sequence complementarity. We focused on endo-siRNAs antisense to protein coding genes, pseudogenes, lincRNAs, and repetitive and transposable elements, as these are the most abundant targets of the endo-siRNA binding AGOs (Table S2). For this analysis we defined 22G-RNAs as reads that are 20-24nt long and 26G-RNAs as reads that are 24-26nt long with no 5′ nucleotide restriction for either sRNA species (Table S6).

Of the 19,999 annotated protein-coding genes, we detected endo-siRNA reads against 19,579 genes across all of our data sets, suggesting that sRNAs are generated against the entire protein-coding transcriptome at some level. 10,127 genes had sRNAs that were enriched at least two fold in association with at least one AGO, indicating that at least half of the protein coding genome has the potential to be regulated by AGO/sRNA complexes at this developmental stage. Hierarchical clustering of the enriched target genes of each AGO enabled us to identify four clear clusters of AGOs that target similar sets of protein coding genes, and could therefore function together in regulating those genes (Fig 5A,B). We compared the gene targets of each AGO to previously described sRNA pathways: (1) ALG-3 and -4 26G-RNA targets, defined as targets that are depleted of sRNAs in *alg-3; alg-4* mutants (1428 targets, Conine *et al*., 2010); (2) ERGO-1 26G-RNA targets, defined as enriched in ERGO-1 IPs (60 targets, Vasale *et al*., 2010); (3) CSR-1 targets, defined as enriched in CSR-1 IPs (4230 targets, Claycomb *et al*., 2009); (4) WAGO targets, defined as the overlap of sRNAs depleted in *rde-3*, *mut-7* and *mago12* (a strain containing null mutations in 12 *wagos*) (1136 targets, Gu *et al*., 2009); and (5) Mutator targets defined as targets that are depleted of sRNAs in *mut-16* mutants (3625 targets, Phillips *et al*., 2012). We compared the gene targets of each AGO to the genes depleted of sRNAs in the RdRP mutants *rrf-1* (131 targets), *rrf-3* (319 targets)*, ego-1* (5403 targets) and *ego-1;rrf-1* (6595 targets, Sapetschnig *et al*., 2015) to determine which RdRP generates each type of AGO-associated sRNA. We also compared the AGO-enriched targets to sRNA targets that are enriched (82 transcripts) or depleted (4357 transcripts) of sRNAs in germline-less *glp-4(bn2)* mutants (Gu *et al*., 2009), representing sRNAs that are enriched in the soma or germline, respectively.

**Figure 5.**
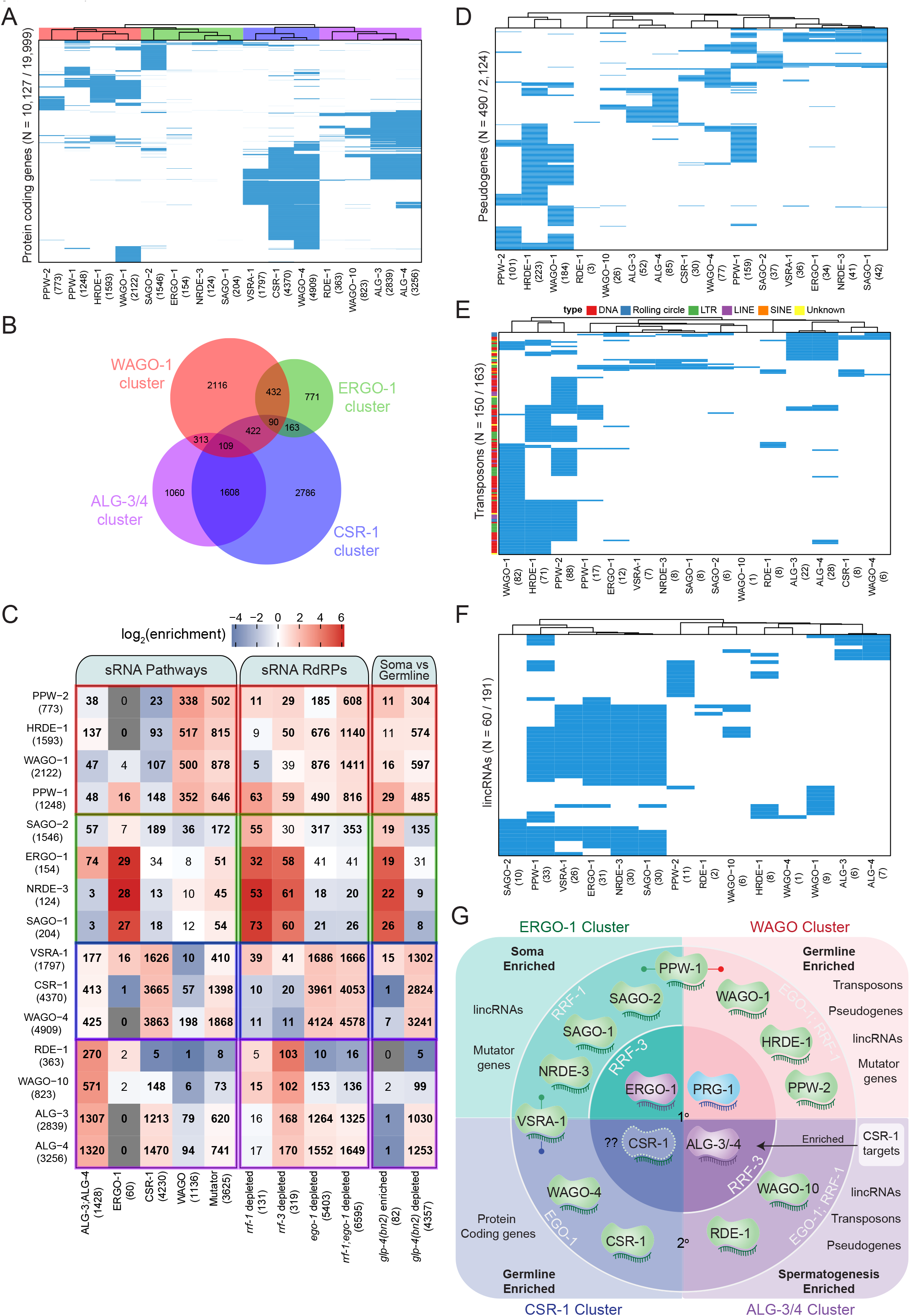
Analysis of endo-siRNA binding AGOs reveals functional categorization of AGOs to regulate distinct genetic elements. (A) Clustering diagram of AGO protein coding gene targets. Each blue line represents a transcript for which endo-sRNAs were enriched 2-fold or more relative to input in both IPs. 22G-RNAs (defined as 21-24nt with no 5′ nucleotide bias) were considered for all AGOs except ALG-3, ALG-4 and ERGO-1, for which only 26G-RNAs (defined as 25-27nt with no 5′ nucleotide bias) were considered. (B) A Venn diagram showing the overlaps of protein coding gene targets of the AGO clusters as highlighted by the color scheme in (A). (C) Enrichment analysis of the AGO protein coding gene targets in previously described datasets. (D) Clustering diagram of AGO pseudogene targets as in (A). (E) Clustering diagram of AGO transposon targets as in (A). (F) Clustering diagram of AGO lincRNA targets as in (A). (G) A schematic summary highlighting the major AGO/sRNAs networks uncovered by endo-siRNA analysis.

The first AGO cluster consists of the WAGOs WAGO-1 (1,814 targets), PPW-2 (636 targets), HRDE-1 (1,295 targets), and PPW-1 (870 targets) (Fig 5A), which we term the WAGO cluster. These AGO complexes are enriched for sRNAs targeting WAGO and Mutator class genes, and are largely depleted of sRNAs targeting ALG-3 and -4, ERGO-1, and CSR-1 class genes (Fig 5C). The targets of these WAGOs also strongly overlap with targets of sRNAs depleted in *ego-1; rrf-1* double mutants (Fig 5C). Moreover, the targets of these WAGOs primarily overlap with transcripts depleted of sRNAs in *glp-4* mutants (Fig 5C). These observations are consistent with the germline expression of WAGO cluster AGOs (Fig 6A). Gene Ontology (GO) analysis of the WAGO cluster targets shows enrichment for kinase activity and protein binding, along with signaling, motility, and morphogenesis (Table S7).

The second AGO cluster consists of the AGOs SAGO-2 (1,537 targets), SAGO-1 (181 targets), ERGO-1 (239 targets), and NRDE-3 (116 targets) (Fig 5A), which we term the ERGO-1 cluster. These AGOs are enriched for previously described ERGO-1 and Mutator target genes and depleted for ALG-3 and -4, CSR-1, and WAGO targets (Fig 5C). By comparing the ERGO-1 cluster targets to genes depleted of sRNAs in *RdRP* mutants, we observed overlap with *rrf-3* and *rrf-1* but not *ego-1* (Fig 5C). The ERGO-1 cluster also overlaps with *glp-4* sRNA enriched transcripts (somatic genes) and is largely depleted of *glp-4* sRNA depleted transcripts (germline genes) (Fig 5C). Consistent with the expression of ERGO-1, these data indicate that the targets of the ERGO-1 cluster are largely somatic. GO analysis of the targets of the ERGO-1 cluster revealed enrichment in various developmental processes, and the most significant terms were those associated with immune, defense, and stress responses (Table S7).

The third cluster consists of the WAGOs CSR-1 (4,182 targets), WAGO-4 (4,815 targets) and C04F12.1/VSRA-1 (1,797 targets), which we term the CSR-1 cluster. The targets of this cluster significantly overlap with the set of previously described CSR-1 targets and are depleted for WAGO targets (Fig 5B,C). This cluster also significantly overlaps with the 26G-RNA targets of ALG-3 and -4 and Mutator targets (Fig 5B,C), consistent with recent observations detailing the sRNA association of each CSR-1 isoform (Charlesworth *et al*., 2021; Nguyen and Phillips, 2021). The CSR-1 strain we used tags both isoforms of CSR-1, and these IPs were performed at a developmental time when CSR-1b is highly expressed in the oogenic germline and CSR-1a is expressed in the intestine. The CSR-1 cluster significantly overlapped with genes depleted of sRNAs in *ego-1* and in *glp-4* mutants, as previously described (Claycomb *et al*., 2009), and consistent with germline expression. The CSR-1 cluster targets are associated with many biological process GO terms (up to 604, Table S7), including terms related to meiosis and chromosome segregation for which CSR-1 is known to be essential (Claycomb *et al*., 2009).

The fourth cluster consists of the AGOs RDE-1 (388 targets), WAGO-10 (877 targets), ALG-3 (2,966 targets), and ALG-4 (3,378 targets) (Fig 5A), which we term the ALG-3/4 cluster. This cluster is enriched for previously described targets of ALG-3 and ALG-4 26G-RNAs and depleted of WAGO targets (Fig 5B,C). The cluster can be further subdivided into ALG-3 and -4 versus WAGO-10 and RDE-1, where ALG-3 and -4 are also enriched for CSR-1 and Mutator targets and WAGO-10 and RDE-1 are depleted for such targets (Fig 5C). These AGOs are also depleted of previously published ERGO-1 26G-RNA targets (Fig 5C). The ALG-3/4 cluster AGOs significantly overlapped with genes depleted of sRNAs in *rrf-3* mutants and in *ego-1* mutants (Fig 5C). The ALG-3/4 cluster showed significant overlap with genes depleted of sRNAs in *glp-4* mutants, consistent with targeting germline enriched genes (Fig 5C). GO term analysis reveals many biological processes are regulated by the ALG-3/4 cluster, including terms associated with gamete generation, and specifically spermatogenesis (Table S7), consistent with previous roles attributed for ALG-3 and ALG-4 (Conine *et al*., 2010), and with the spermatogenic-restricted expression patterns of ALG-3, ALG-4 and WAGO-10 (Fig 6B) (Charlesworth *et al*., 2021).

**Figure 6.**
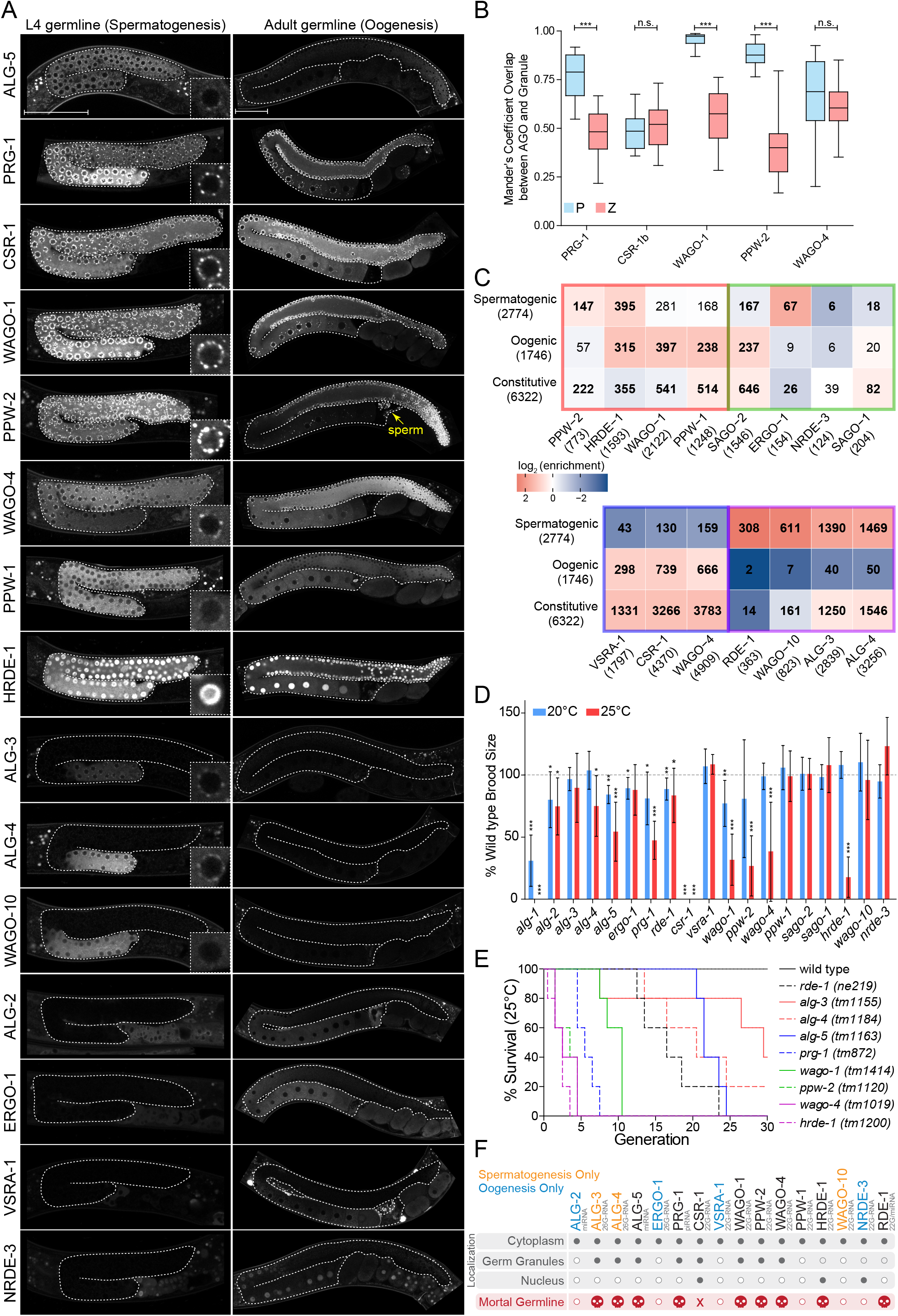
AGOs are differentially expressed in the germline and differentially regulate germline gene expression to promote fertility. (A) Expression patterns of germline AGOs in L4 (left) and adult (right) germlines. Inset image zoomed in at an individual germ cell nucleus. Yellow arrow points to sperm within the spermatheca. Scale bar represents 50μm. (B) Quantification of the number of GFP::AGO pixels that overlap with PGL-1::mRFP (blue) or HA::TagRFP::ZNFX-1 (coral) pixels using Mander’s Correlation. For each data set, five Z stacks of proximal germline regions from six different animals per strain were counted (N = 30 slices, approximately 80–100 nuclei per worm). *** = *p*-value < 0.001, n.s. = not significant. One way ANOVA with Bonferroni’s Post Hoc multiple comparison test. (C) Analysis of the enriched sRNA targets in each of the AGOs in comparison to germline constitutive, oogenic and spermatogenic expressed genes (Ortiz *et al*., 2014). Bolded numbers indicate significant enrichment or depletion (p < 0.05), Fisher’s Exact Test. The colored borders represent the AGO clusters as defined in Figure 5A. (D) Brood size analysis of all ago mutants at 20°C and 25°C. Data was aggregated from different experiments and normalized to the mean of wild-type control samples. * = p < 0.05, ** = p < 0.01, *** = p < 0.001, two sided t-test. N ≧ 10 P0 worms. (E) Mortal germline assay of *ago* mutants showing a Mrt phenotype in (D). N = 5 P0 worms. (F) A summary of the spatial and temporal localization of AGOs in the germline and Mrt phenotypes. CSR-1 has an “X” to indicate it is essential.

#### sRNA profiles suggest similar sRNA biogenesis and targeting mechanisms

The distribution of sRNAs along sets of target transcripts may provide insights into the mechanisms of sRNA biogenesis and target regulation. To visualize sRNA distribution across sets of protein-coding targets for each AGO, we used metagene plots (Fig S3). Patterns of sRNA distribution were generally more similar within AGO clusters than between clusters, however there were some differences. The WAGO cluster generally displays high levels of sRNA enrichment, and these sRNAs are present over most of the transcript (Fig S3). The ERGO-1 cluster has the highest levels of sRNAs overall, which tend to be either distributed over most of the target transcript (SAGO-1 and NRDE-3), or biased toward the 3′ end (SAGO-2 and ERGO-1) (Fig S3). The CSR-1 cluster shows strong similarity between the WAGO-4 and CSR-1 sRNA distribution profiles, with enrichment at the 3′ and to a lesser extent the 5′ end (Charlesworth *et al*., 2021), while the profile of C04F12.1/VSRA-1 sRNAs appears more similar to that of SAGO-1 and NRDE-3 (Fig S3). Consistent with previous observations, the 26G-RNAs associated with the ALG-3/-4 pathway are enriched at the 5′ and 3′ ends of target transcripts (Conine *et al*., 2010) (Fig S3). We observed a similar 5′ enrichment for the 22G-RNAs associated with RDE-1 and WAGO-10, but not a 3′ sRNA enrichment (Fig S3). These observations may be indicative of differences in sRNA biogenesis or targeting mechanisms, and will necessitate further study.

#### Rapidly evolving and “non-self” elements in the genome are targeted by distinct AGO/sRNA pathways

Although protein coding genes comprise the majority of sRNA targets, we also examined pseudogenes, repetitive and transposable elements, and lincRNAs. We detected sRNA reads against 1,741 out of 2,124 annotated pseudogenes and found that 490 were enriched with antisense sRNAs across our IP data sets. Hierarchical clustering of the enriched targets of each AGO largely resembled the four clusters observed for the protein coding targets of the AGOs (Fig 5D) with two exceptions: PPW-1 and C04F12.1/VSRA-1 clustered with the ERGO-1 pathway instead of the WAGO-1 pathway (PPW-1) and CSR-1 pathway (C04F12.1/VSRA-1). The majority of the pseudogenes were targeted by the WAGO cluster (Fig 5C, Fig S4A). Many pseudogenes were specifically enriched in one pathway over the others. For example, out of the 95 pseudogenes targeted by the ALG-3/4 cluster, 59 were unique to this cluster (Fig S4A). These differences may reflect tissue-specific regulation by different pathways or partitioning of transcripts into separate pathways.

We detected reads against all 163 annotated transposable elements. Of these, 150 were enriched for antisense sRNAs across our IP data sets (Fig 5E). This indicates that all transposable elements have the potential to be regulated by sRNAs and that most of them are likely to be highly regulated by AGO/sRNA complexes. As with pseudogenes, the PPW-1 and C04F12.1/VSRA-1 targets overlapped with the ERGO-1 cluster targets, but this cluster appears to regulate a relatively small number of transposable elements overall, with only one TE uniquely regulated by this cluster (Fig 5E, Fig S4B). ALG-3/-4 regulate a group of 36 transposable elements that are likely to be expressed during spermatogenesis. Most transposable elements (134) are targeted by the WAGO cluster AGOs, with 104 of these being unique to this cluster (Fig 5E, Fig S4B). This suggests that the WAGO cluster is the major regulator of transposable elements, particularly in the germline.

Transposable elements can be subdivided based on their class and method of transposition: DNA transposons (cut-and-paste DNA transposons, Rolling circle replication DNA transposons), and retrotransposons (LTR, LINE, SINE). We observed that different AGOs show differential sRNA enrichment for specific types of transposons. For example, the ERGO-1 cluster AGOs are relatively depleted of cut-and-paste DNA transposon, LINE and LTR element-targeting sRNAs but enriched for Rolling circle-targeting sRNAs (Fig S4C). In contrast, the WAGO cluster was generally enriched for sRNAs targeting all other elements except Rolling circle transposons (Fig S4C).

We detected reads against 168 of the 191 annotated lincRNAs (defined in (Nam and Bartel, 2012) and WormBase version WS276), across all IP data sets, and, among these, 60 of these were enriched for antisense sRNAs (Fig 5F). Again, PPW-1 and C04F12.1/VSRA-1 lincRNA targets overlapped with the ERGO-1 cluster targets (Fig 5E). The majority of lincRNAs (48) are targeted by the WAGO cluster, followed by the ERGO-1 cluster (35). Many of the targets overlapped between the WAGO and ERGO-1 clusters (26), with the majority of this overlap (25/26) originating from PPW-1 targets (Fig S4D). 17 lincRNA targets were specific to the WAGO cluster, three were specific to the ALG-3/4 cluster, and three were specific to the ERGO-1 cluster (Fig S4D). Our results point to lincRNAs as another important, yet poorly studied category of sRNA targets with the potential for tissue-specific regulation.

#### Hierarchy and interconnected regulation of multiple sRNA pathways

To begin to understand why *C. elegans* encodes so many AGOs, we must first understand how the different sRNA pathways interact with each other. Loss of one component may affect the sRNA landscape in the organism, potentially allowing us to infer the hierarchical relationship between sRNA/AGO pathways. Our analysis indicated that some AGOs may have distinct roles and participate in different pathways depending on the tissue in which they are expressed. To address how loss of one component may affect the system, we sequenced sRNAs from all *ago* mutants in triplicate. We examined how the sRNAs antisense to protein-coding, pseudogene, repetitive and transposable element, and lincRNA targets of each AGO are affected in each *ago* mutant (Fig S5).

It is possible that AGOs may regulate other AGOs, especially as among the targets of AGO/sRNAs were other *agos* (Fig S4E), indicating there may be regulatory networks in place in which sRNA pathways can regulate others, and/or participate in regulatory feedback loops. The WAGOs CSR-1 and WAGO-4 form a hub that targets most of the constitutively-expressed germline AGOs (see below), while the spermatogenesis-specific AGOs, ALG-3, ALG-4, and WAGO-10 form a self-regulatory hub (Fig S4E). Below we outline several observations from this analysis and put these results in the context of our current knowledge of hierarchy in sRNA targeting.

Loss of the miRNA-binding AGOs ALG-1, ALG-2 and ALG-5 did not have large effects on the endo-siRNA pathways (Fig S5), therefore, we begin our analysis by focusing on established primary AGOs: PRG-1, ERGO-1, and ALG-3/4. Given that CSR-1 seems able to play both primary and secondary AGO roles, we will include it in this group. Loss of the piRNA-binding primary AGO PRG-1, which results in the loss of most piRNAs (Fig 4B), led to downregulation of secondary sRNAs associated with the WAGO cluster AGOs sRNAs (Montgomery *et al*., 2021). This result is consistent with the model that piRNA targeting induces the production of 22G-RNAs which are then loaded into WAGO-1 and HRDE-1 (Lee *et al*., 2012; Cornes *et al*., 2022). Our complete analysis reveals that PPW-1 and PPW-2 also work within the context of PRG-1 targeting. With similar logic, loss of the 26G-RNA-binding primary AGO ERGO-1 resulted in the loss of ERGO-1 cluster sRNAs (Fig S5). Consistent with our observation that NRDE-3 sRNAs are depleted, it was previously shown that the translocation of NRDE-3 to the nucleus requires sRNA binding, and loss of ERGO-1 results in NRDE-3 remaining cytoplasmic (Guang *et al*., 2008). Our complete analysis reveals that SAGO-1- and SAGO-2-associated 22G-RNAs are depleted upon loss of ERGO-1, indicating that SAGO-1 and SAGO-2 also work within the context of ERGO-1 targeting.

We observed that the sRNAs of some 22G-RNA-associated AGOs are differentially affected by loss of primary AGOs. For example, PPW-1-bound sRNAs that target protein-coding genes, pseudogenes, and transposons are depleted in *prg-1* mutants but not *ergo-1* mutants (Fig S5). Conversely, PPW-1-bound sRNAs that target lincRNAs are depleted in *ergo-1* mutants but not *prg-1* mutants (Fig S9). When taken together with the clustering analysis of targets (Fig 5A,D-F), it appears that PPW-1 acts downstream of both the PRG-1 piRNA pathway and the ERGO-1 26G-RNA pathway. This PPW-1 duality is dependent on the tissue (germline and soma) in which it is expressed. Thus, it is likely that in the germline PPW-1 acts downstream of PRG-1 piRNAs, and in the soma it acts downstream of ERGO-1 26G-RNAs. A similar observation is made for C04F12.1/VSRA-1-bound sRNAs. C04F12.1/VSRA-1 acts downstream of ERGO-1 26G-RNAs to target pseudogenes, transposons and lincRNAs, and acts in conjunction with CSR-1 22G-RNAs to target protein coding genes in the germline (Fig 5A,D-F, Fig S5). Because of this dual association with the CSR-1 cluster and the ERGO-1 cluster, we named C04F12.1 VSRA-1, for Versatile Small RNAs AGO.

Loss of the 26G-RNA-binding AGOs ALG-3 and ALG-4 individually did not affect sRNA populations, likely due to their partial redundancy with each other (Fig 1E, (Conine *et al*., 2010)). Interpreting the loss of CSR-1 is more complicated, given that it targets nearly one quarter of the protein-coding genome, is encoded as two isoforms with distinct expression profiles and functions, and is required for both gene licensing as well as silencing (Charlesworth *et al*., 2021; Nguyen and Phillips, 2021). Loss of CSR-1 resulted in depletion of CSR-1 cluster sRNAs targeting protein-coding genes (Fig S5), however, the sRNA levels of other AGOs including WAGO-1, HRDE-1, PPW-1, SAGO-2, and SAGO-1 were also decreased. This may be due to secondary effects arising from loss of CSR-1, which appears to be capable of regulating many other AGOs (Fig S4E).

Loss of single 22G-RNA-associated AGOs had different effects depending on the AGO. Like loss of CSR-1, loss of WAGO-4 also resulted in depletion of CSR-1 cluster sRNAs. Loss of the WAGO cluster WAGOs had varying effects. Loss of WAGO-1 or HRDE-1 resulted in the depletion of PPW-2-associated sRNAs, while loss of HRDE-1 alone also resulted in depletion of WAGO-1- and HRDE-1-associated sRNAs. This indicates that HRDE-1 is required for the stability of most of the WAGO cluster associated sRNAs, likely reflecting the requirement of HRDE-1 in producing tertiary sRNAs (Sapetschnig *et al*., 2015). Loss of SAGO-1 resulted in downregulation of ERGO-1 cluster-associated sRNAs primarily targeting pseudogenes and lincRNAs. This suggests SAGO-1 is the main AGO targeting these elements downstream of ERGO-1/26G-RNAs. Loss of PPW-2 did not result in downregulation of WAGO cluster small RNAs, but rather CSR-1 cluster sRNAs, highlighting the potential for interplay between pathways.

In sum, we have defined which classes of genetic elements are targeted by each AGO and the genetic requirements for each *ago* in accumulating sRNAs targeting these elements. These results define the relationships between different AGO pathways and highlight the complexity of target regulation. The clear targeting of different genetic elements by different clusters of AGOs implies that features intrinsic to the target transcript likely encode determinants for AGO/sRNA pathway specificity. Furthermore, the spatiotemporal expression profile of targets, sRNA biogenesis components, and AGOs are also likely to be major contributors to the patterns we have observed.

### AGOs have distinct spatiotemporal localization patterns during development

To link our molecular observations with the AGO expression profiles and gain insight into where each AGO exerts its function, we visualized GFP fluorescence using confocal microscopy at each stage of worm development, revealing that AGOs are expressed in a variety of tissues throughout the life cycle. We identified 16/19 AGOs expressed in the germline, consistent with known roles for AGOs in the germline. Eight of these show germline restricted expression and the other eight are also expressed in somatic tissues. 11/19 AGOs are expressed in the soma, with 3 AGOs being somatically restricted (SAGO-1, SAGO-2, ALG-1). We observed AGOs expressed in various somatic tissues, including, but not limited to, the: vulva, hypodermis, muscle, seam cells, intestine, neurons, somatic gonad, and spermatheca (Fig 7E, Figs S6-13). AGOs expression levels vary, and low expression levels or expression under specific (e.g. stress induced) conditions may have precluded detection of some AGOs in some tissues. Specifically, RDE-1 was hardly detectable as a translational fusion. However, the first step in our CRISPR protocol generates a transcriptional reporter (Dickinson *et al*., 2015), by which we observed strong GFP expression under the control of the *rde-1* promoter. Thus, for RDE-1, we use the transcriptional reporter to deduce expression patterns (Fig S13).

**Figure 7.**
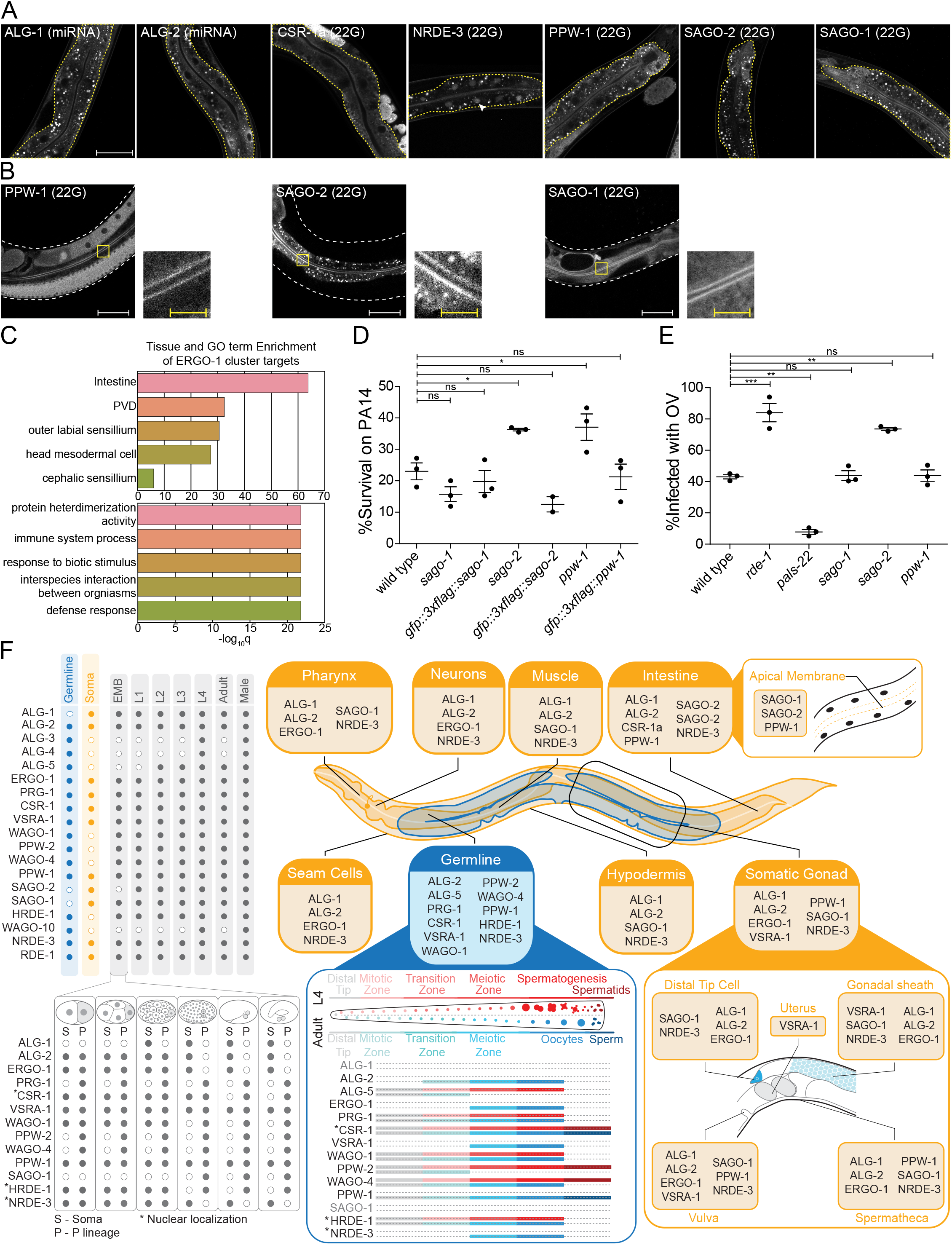
AGOs are expressed in multiple somatic tissues and several are required for normal immunity towards pathogens. (A) AGOs expressed in the intestine. Adult worms shown. Brackets indicate the type of sRNA AGOs associate with. Intestines are outlined in yellow. Arrowhead indicates intestine cell nuclei. Scale bar represents 50μm. Bright foci throughout intestinal tissue are autofluorescent gut granules. (B) Apical intestinal membrane localization of PPW-1, SAGO-2, and SAGO-1. Worm body is outlined in white. Zoomed in panels are indicated with a yellow box. White scale bar represents 50μm. Yellow scale bar represents 10μm. Note that PPW-1 is also expressed in the germline. (C) Tissue enrichment analysis (top) and Gene Ontology analysis (bottom) of the ERGO-1 cluster sRNA targets. (D) Percent of worms alive after 72 hours of exposure to a *P. aeruginosa* PA14 lawn is shown for each strain. This is a representative experimental run out of 3 conducted. Each dot represents 50 worms. * = p-value < 0.05, n.s. = not significant. One way ANOVA with Dunnett’s Post Hoc multiple comparison test. (E) Percent of worms infected with Orsay Virus (OV) 16 hours post infection is shown for each strain. Each dot represents >=100 worms. N = 3. ** = *p*-value < 0.01, *** = *p*-value < 0.001, n.s. = not significant. One way ANOVA with Tukey Post Hoc multiple comparison test. (F) A summary of the expression patterns of all *C. elegans* AGOs throughout development.

### Germ granule-localized AGOs are required for transgenerational fertility

The majority of AGOs are expressed in the germline (16/19), where they display striking temporal and spatial localization patterns. Nine AGOs are constitutively expressed throughout the germline from the emergence of the P1 cell through oogenesis in adults: RDE-1, ALG-5, PRG-1, CSR-1b, WAGO-1, PPW-2, WAGO-4, PPW-1 and HRDE-1 (Figs S6-13). Of these, two AGOs are more highly expressed at the mitotic zone of the adult oogenic germline: ALG-5 and PPW-2 (Fig 6A). The other seven germline-expressed AGOs exhibit gamete-specific expression patterns. ALG-3, ALG-4, WAGO-10, and the CSR-1a isoform are expressed only during spermatogenesis, at the L4 stage in hermaphrodites and constitutively in the male germline (Charlesworth *et al*., 2021; Nguyen and Phillips, 2021)(Fig 6A and Fig S12). Only PPW-2 and CSR-1b are detectable in spermatids (Fig 6A) (Charlesworth *et al*., 2021; Schreier *et al*., 2022). ALG-2, ERGO-1, and NRDE-3 are specifically expressed in oocytes starting at the young adult hermaphrodite stage (Fig 6A). C04F12.1/VSRA-1 is also restricted to the oogenic germline in adult worms, and shows expression in the primordial germ cells of L1 larvae (Fig 6A and Fig S7). Published mRNA expression data from dissected hermaphrodite and male germlines shows largely the same mRNA expression patterns as the tagged AGO proteins (Fig S14B) (Tzur *et al*., 2018). All germline-expressed AGOs, except those that are expressed only during oogenesis, are also expressed in the male germline, and display the same patterns of expression (Fig S12).

The germline AGOs encompass different subcellular localization profiles. Half (8/16) of the germline AGOs are enriched in perinuclear phase-separated germ granules: ALG-3, ALG-4, ALG-5, PRG-1, CSR-1a, b, WAGO-1, PPW-2 and WAGO-4 (Fig 6A, 6F). PPW-1 is also weakly detected in germ granules, mostly restricted to the pachytene region of the adult germline (Fig 6A). There are four types of germ granules in *C. elegans* that are thought to play different roles in sRNA pathways: P granules, Z granules, Mutator Foci, and SIMR Foci (Sundby, Molnar and Claycomb, 2021). PRG-1, CSR-1, and WAGO-1 were previously reported to co-localize with P granules (Batista *et al*., 2008; Claycomb *et al*., 2009; Gu *et al*., 2009), and WAGO-4 was identified as a Z granule component (Wan *et al*., 2018). To determine which types of germ granules the germline-constitutive WAGOs, CSR-1b, and PRG-1 are associated with, we determined the overlap in pixels between the AGOs and components of P granules (PGL-1) or Z granules (ZNFX-1) using confocal microscopy in adult worms. The CSR-1 cluster AGOs CSR-1 and WAGO-4 overlapped roughly equally with both P and Z granule pixels, whereas the WAGO cluster AGOs PPW-2 and WAGO-1 were strongly biased toward overlap with P granules (Fig 6B). PRG-1 was intermediate to both of these phenotypes. Thus, AGOs with similar target preferences display similar germ granule localization patterns (Fig 6B).

Two AGOs are predominantly nuclear: HRDE-1 and NRDE-3 (Fig 6A, 6F). While HRDE-1 is known to be a nuclear germline AGO required for RNAi inheritance (Buckley *et al*., 2012), NRDE-3 was previously shown to be a somatic nuclear AGO, required for somatic RNAi inheritance (Guang *et al*., 2008). That we observed NRDE-3 localizing to oocyte nuclei may explain how it is capable of propagating certain RNAi responses into the next generation.

Due to the split in AGOs between germline-constitutive and sex-specific expression, we asked whether targets were also sex-biased. A previous report defined the spermatogenic, oogenic, and germline-constitutive transcriptomes of *C. elegans* (Ortiz *et al*., 2014). Therefore, we compared the protein-coding target transcripts for 22G- and 26G-RNA binding AGOs to this dataset (Fig 6C). The 22G-RNA targets of the WAGO cluster AGOs PPW-2, HRDE-1, and WAGO-1 were depleted of germline constitutively -expressed transcripts. Instead, PPW-2 and HRDE-1 targets were enriched for spermatogenic transcripts, and WAGO-1, HRDE-1 and PPW-1, targets were enriched for oogenic transcripts (Fig 6C). This suggests that the WAGO cluster can be further subdivided and is responsible for sex-specific gene regulation during the young adult stage when oogenesis occurs: PPW-2 regulates spermatogenic genes, and WAGO-1 and PPW-1 regulate oogenic genes. Both branches of this WAGO cluster appear to act upstream of or in parallel with the nuclear HRDE-1. The 22G-RNA targets of the CSR-1 cluster AGOs CSR-1, C04F12.1/VSRA-1, and WAGO-4 were depleted of spermatogenic transcripts and were enriched for both oogenic and germline-constitutive transcripts in the young adult stage (Fig 6C). We have previously reported that CSR-1b and WAGO-4 associate with oogenic and germline constitutive transcripts during the L4 stage, and in contrast CSR-1a enriches for spermatogenic transcripts that overlap with the ALG-3/4 cluster (Charlesworth *et al*., 2021). During the L4 stage, targets of the ALG-3/4 cluster AGOs WAGO-10, ALG-3 and -4 were all enriched for spermatogenic transcripts and ALG-3 and -4 also showed enrichment for germline constitutive transcripts (Fig 6C). Similarly, during the young adult stage, the ALG-3/4 cluster AGO RDE-1 targets were also enriched for spermatogenic transcripts. Thus, germline AGOs have differential roles in regulating sex-specific germline transcripts in a spatiotemporal manner.

Given their germline expression, we next asked whether each germline AGO is required for fertility. Previous work examined a role for each *C. elegans ago* in fertility and found that only *csr-1* is essential at the normal laboratory-culturing temperature of 20°C (Yigit *et al*., 2006). However, in recent years it has become apparent that stressful conditions, such as elevated temperature, can have a substantial impact on the fertility of mutants. This temperature-dependent fertility defect can manifest in the first generation subsequent to a temperature shift or take several generations to reach its full impact. Several germ granule component mutants and *ago* mutants, including *hrde-1*, *prg-1*, *wago-4*, and *ppw-2* have been shown to exhibit a Mortal germline (Mrt) phenotype, in which fertility decreases over several generations at elevated temperature (Buckley *et al*., 2012; Simon *et al*., 2014; Wan *et al*., 2018; Schreier *et al*., 2022). We therefore performed brood size assays at 20°C and 25°C for recently outcrossed hermaphrodites of each single *ago* mutant (Fig 6D). At 20°C, seven *ago* mutants showed compromised fertility: *alg-1*, *alg-2*, *alg-5*, *ergo-1*, *prg-1*, *rde-1* and *wago-1* (Fig 6D). When shifted to 25°C, the same mutants displayed a low brood size, which was even more pronounced in some cases (for example, *alg-1* was sterile). We also observed fertility defects not present at 20°C for four *ago* mutants: *alg-4*, *ppw-2*, *wago-4*, and *hrde-1* (Fig 6D). *csr-1* mutants were sterile at both temperatures (Fig 6D). These results indicate that more than half (10/19) of the *C. elegans* AGOs contribute individually to optimal fertility.

To further assess the role of temperature stress on the fertility of *ago* mutants, we conducted a Mrt assay, following worms for 30 generations at 25°C. We observed complete loss of fertility over varied numbers of generations for *alg-5* (25 generations), *rde-1* (24), *prg-1* (8), *wago-1* (11), *ppw-2* (5), *wago-4* (5) and *hrde-1* (4) (Fig 6E). We also observed substantially reduced fertility that took longer to manifest in *alg-3* and *alg-4* single mutants (Fig 6E). These *agos* have been shown to act partially redundantly (Conine *et al*., 2010) and loss of both results in sterility at 25°C (Fig 1E). Collectively, these findings implicate 11/19 AGOs and every type of sRNA pathway in maintaining fertility over generations.

Understanding of the Mrt phenotype is currently incomplete. Factors involved in telomere and chromatin maintenance as well as germ granule components result in a Mrt phenotype when mutated, and it has been suggested that this phenotype occurs when epigenetic stress accumulates over generations (Cecere, 2021; Sundby, Molnar and Claycomb, 2021). The *hrde-1* Mrt phenotype at 25°C was shown to be reversible upon transferring the animals back to 20°C, suggesting it may not be genotoxic in origin (e.g. transposon mobilization) (Spracklin *et al*., 2017). Similarly, the Mrt phenotype of *prg-1* mutants is reversible when the animals are subjected to starvation (Simon *et al*., 2014). We therefore explored whether *ago* mutant Mrt phenotypes display the same characteristics as each other and other Mrt mutants. We focused on the WAGOs that have the most severe Mrt phenotype: *wago-1*, *ppw-2*, *wago-4* and *hrde-1*.

To test whether genotoxic stress is responsible for the Mrt phenotype, we maintained animals at 25°C for three generations, then transferred them back to 20°C and measured brood size at each generation. For all four mutants, we observed the same pattern; a reversal of the Mrt phenotype (Fig S15A), suggesting that genotoxic stress is not responsible for the Mrt phenotype for any of these w*agos*.

Given that the Mrt phenotype results in a gradual decrease in brood size, we hypothesized that the germline might suffer from general proliferation defects. To test this, we examined the germlines of the Mrt *wagos,* focusing on the mitotic zone. Counting germ cells at the mitotic zone at 20°C and 25°C revealed Mrt *wagos* produced fewer germ cells than wild-type controls, suggesting a proliferation defect (Fig S15B). The Mrt *wagos* also showed a reduced number of oocytes in adult animals (Fig S15C). These results are consistent with a previous observation that *hrde-1* mutants have variable defects in both oogenesis and spermatogenesis (Buckley *et al*., 2012) and suggest that the defects observed in the Mrt *wagos* arise early in germ cell proliferation. In our analysis of the sRNA complements and targets of these WAGOs, we showed that WAGO-1, PPW-2 and HRDE-1 cluster together to target repetitive elements and other silenced germline genes (Fig 5D,E), while WAGO-4 mainly targets germline expressed genes that are co-targeted by CSR-1 (Fig 5A,C). Whether a common mechanism or different mechanisms underlie the Mrt phenotype in these *ago* mutants from different clusters remains to be resolved.

### AGO somatic expression and tissue-specific gene regulation

We found broad expression patterns for many AGOs in the soma, suggesting they may have gene regulatory roles in tissues where sRNA pathways have not been deeply explored (Fig 7E). The intestine is a key interface between the worm and its environment. Foreign RNA from bacterial food or pathogens, such as viruses, can enter the worm via the intestinal epithelium (Franz *et al*., 2014; Braukmann, Jordan and Miska, 2017). The intestine is the somatic tissue that expresses the most AGOs: ALG-1, ALG-2, ERGO-1, RDE-1, CSR-1a, PPW-1, SAGO-2, SAGO-1, and NRDE-3 (Fig 7A, Fig S13) (Charlesworth *et al*., 2021). Most AGOs were broadly cytoplasmic; NRDE-3 was nuclear; and three AGOs, SAGO-1, SAGO-2, and PPW-1, showed an accumulation along the apical membrane of the intestinal cells. SAGO-2 is not detectable in other tissues (Fig S6-13).

Given the role of the intestine, and the abundance of AGOs representing three of the four sRNA pathways (miRNA, 22G- and 26G-RNAs), we wondered how these pathways might function here. The entire ERGO-1 cluster is represented in the intestine, along with an isoform of CSR-1, CSR-1a (Fig 7A). Our previous work demonstrated that CSR-1a silences genes involved in immune and pathogen defense responses in the intestine. Loss of *csr-1a* led to an upregulation of these genes and enhanced worm survival on the bacterial intestinal pathogen *Pseudomonas aeruginosa* (PA14) (Charlesworth *et al*., 2021). Further, the GO terms associated with the ERGO-1 cluster are enriched for immune and defense responses, particularly in the intestine (Fig 7B, Table S7). GO terms associated with the targets of individual AGOs were primarily derived from the targets of SAGO-2 (Table S7), which is closely related to PPW-1 and SAGO-1 (Fig 1A). Additionally, PPW-1 targets were enriched for the innate immune signaling pathway, MAPK (Table S7) (Kim, 2002). Given their localization and GO analysis, we hypothesized that SAGO-1, SAGO-2, and PPW-1 may play a role in immune responses to pathogenic bacteria. In agreement, survival assays of *sago-2*, *ppw-1* and *sago-1* mutants on PA14 revealed that both *sago-2* and *ppw-1* are partially resistant to killing by PA14 infection (Fig 7C).

Previous research has shown that PPW-1, SAGO-1, and SAGO-2 act as secondary AGOs, downstream of RDE-1 in exo-RNAi (Yigit *et al*., 2006), with SAGO-1 and SAGO-2 being required for efficient exo-RNAi in the soma, and PPW-1 for efficient exo-RNAi in the germline. Over-expression of any of these three AGOs in the muscle cells of a compound *ago* mutant rescues the RNAi deficiency of the mutant (Yigit *et al*., 2006). RDE-1 and other components of the exo-RNAi machinery are also involved in antiviral responses against the Orsay virus. RDE-1 targets the viral RNA and recruits the RdRP RRF-1 to generate secondary 22G-RNAs that combat the virus. Consistent with this, loss of *rde-1* or the RdRP machinery leads to increased viral proliferation (Felix *et al*., 2011).

Given our understanding of the roles of the secondary AGOs in exo-RNAi, we hypothesized that viral secondary 22G-RNAs could be loaded into these AGOs, and that their mutation could render the worms more sensitive to Orsay virus. To test this, we analyzed the response of *sago-2*, *ppw-1* and *sago-1* mutants to infection by Orsay virus, using *rde-1* and *pals-22* as controls for sensitivity and resistance, respectively (Reddy *et al*., 2019) (Fig 7D). Only *sago-2* showed a higher infection level, which phenocopied that of *rde-1* mutants (Fig 7D). Thus, SAGO-2 may have dual roles in mediating immunity in the intestine. First, SAGO-2 targets immune response genes, and loss of *sago-2* enhances the ability of the worms to survive on PA14. Conversely, loss of *sago-2* decreases viral RNA targeting, likely by RNAi mechanisms in which SAGO-2 is loaded with secondary 22G-RNAs that were generated after targeting of viral RNA by the primary AGO RDE-1.

## Discussion

Here we have analyzed the 19 AGO proteins in *C. elegans* using CRISPR-Cas9 genome editing and next generation sequencing. Analysis of the expression patterns and sRNA complements of each AGO identifies sRNA regulatory networks employed in tissues throughout the animal, and reveals specific and shared features of each AGO, advancing understanding of the functions and mechanisms of these pathways in the context of a whole animal.

### *C. elegans* small RNA pathways

Our analysis provides a framework for categorizing the AGOs and their sRNAs. Consistent with current models, the AGOs can be divided into four groups based on the type of sRNA they interact with: (1) the miRNA binding classical AGOs, ALG-1, ALG-2, ALG-5, and RDE-1; (2) the piRNA binding PIWI, PRG-1; (3) the 26G-RNA binding classical AGOs, ERGO-1, ALG-3, and ALG-4; and (4) the 22G-RNA binding WAGOs, CSR-1, VSRA-1, WAGO-1, PPW-2, WAGO-4, PPW-1, SAGO-2, SAGO-1, HRDE-1, and NRDE-3. Our analysis of the 22G- and 26G-RNA binding AGOs revealed they can be further classified into 4 major clusters based on their targets: (1) the CSR-1 cluster: CSR-1, WAGO-4, and C04F12.1/VSRA-1, which target germline expressed protein coding genes; (2) the WAGO cluster: WAGO-1, PPW-2, and HRDE-1 which target silenced germline genes, pseudogenes and repetitive/transposable elements; (3) the ERGO-1 cluster: ERGO-1, PPW-1, SAGO-1, SAGO-2, and NRDE-3 which target many somatic genes, pseudogenes and lincRNAs; and (4) the ALG-3/4 cluster: ALG-3, ALG-4, and WAGO-10, which are restricted to the spermatogenic germline and predominantly target spermatogenesis genes.

Among these groups, several AGOs bind to one type of sRNA (e.g. ALG-1, -2, -5, and PRG-1), while others have broader specificity or act as scavengers (e.g. RDE-1). We observed a physical association between the 26G-RNA binding AGOs and miRNAs, which may reflect coordinated regulation of transcripts by both miRNAs and 26G-RNAs. The 22G-RNA binding AGOs represent a more varied group, in which some AGOs cluster differently depending on the portion of the genome they target (e.g. VSRA-1 clusters with CSR-1 to target protein coding genes, and with ERGO-1 to target lincRNAs). However, how sRNAs and targets are “selected” by different AGOs remains poorly understood.

### AGO/small RNA specificity

Individual AGO target specificity involves many factors, including: the intrinsic structural and biochemical properties of the AGO; the sRNA biogenesis mechanisms and biochemical features; the features and co-factors of target transcripts; and the expression and localization patterns of AGOs, sRNA machinery, and targets. For instance, it has been shown that the preference for a specific 5′ nucleotide is determined in large part by interactions between the sRNA, the 5′ binding pocket within the MID domain and another region of the AGO termed the specificity loop (Ma *et al*., 2005; Frank, Sonenberg and Nagar, 2010). In *C. elegans*, the biogenesis mechanisms of sRNAs contribute to specificity through differences in 5′ nucleotide chemistry, in which miRNAs, piRNAs, and 26G-RNAs possess a mono-phosphate (DICER-dependent), whereas the 22G-RNAs possess a tri-phosphate (DICER-independent). This difference is reflected in the AGO phylogeny; the AGOs more closely related to AGOs in other species retained the ability to bind 5′ mono-phosphorylated nucleotides and possess highly conserved residues within the 5′ binding pocket (e.g. Y529, K533, Q545, K570, C546 in hAGO2) while the WAGOs preferentially bind tri-phosphorylated nucleotides and possess a more divergent set of residues in these positions. These results, coupled with structure-function based analyses *in vivo,* will enable us to understand the mechanisms of AGO loading and sRNA preference more comprehensively, despite a lack of *in vitro* sRNA loading assays.

We observed that most of the transcriptome, including nearly all protein coding genes, has 22G- and 26G-RNAs directed against it. This suggests that sRNA production (1) happens broadly across tissues, and (2) that most of the transcriptome has the potential to become a substrate for RdRP activity. However, we did not observe sRNA enrichment in AGOs for all of the transcripts with detectable sRNAs. This could be for a number of reasons. First, our experimental design and analysis pipeline is biased for more abundant and more enriched sRNAs, to produce a high-confidence set of targets for each AGO. If we reduced the thresholds of 5RPM, two-fold enrichment, and requirement for enrichment in both replicates, we might detect additional sRNAs (and thus targets) enriched in association with AGOs that occur at very low abundance or are expressed in a small number of cells. Second, features of the transcript are likely to influence the extent to which it can serve as a template for sRNA synthesis, including: sequence motifs, intron/exon, content, 3′ UTR and poly-A length, secondary structure, expression level, association with other RBPs, and subcellular routing and/or localization. Third, targeting by an AGO is generally thought to initiate an sRNA amplification loop.

Thus, targets for which sRNAs are successfully loaded into AGOs and that reach a critical threshold of AGO regulation may be the “winners” that we have detected in our sRNA sequencing. In-depth computational analyses of the features of high confidence target transcripts will reveal specific features of “successful” targets. Measuring transcript levels and localization in specific cell types, using single cell-seq and smFISH, will inform our understanding of what target levels and which subcellular locations (for example, germ granules) are associated with high levels of sRNAs. Finally, examination of RdRP mechanisms in specific cell types and *in vitro* are necessary to fully understand AGO/sRNA specificity.

### Temporal and spatial specificity of small RNA regulation

Our analysis maps the expression patterns of every AGO throughout development. While some AGOs are broadly expressed in a variety of tissues, others are restricted to specific tissues, cell types, and developmental stages. For example, ERGO-1, ALG-2 and NRDE-3 are expressed in similar patterns within a variety of tissues that include neurons, the somatic gonad, intestine, and oocytes. PRG-1, which has mainly been studied for its role in germline gene regulation, is also present within muscle during larval stages. It remains to be determined if AGOs have the same functions and targets in each cell type in which they are expressed, and whether the targets change during development.

Our analysis mainly focused on the L4 to YA transition stage in *C. elegans,* because all AGOs, aside from the spermatogenesis specific AGOs, are expressed at this time. Also, sRNA populations change in abundance during development (Ambros *et al*., 2003; Ruby *et al*., 2006), which could reflect changes in expression or association with AGOs. Using the tools developed here, it is possible to probe different life stages of *C. elegans* to observe the temporal dynamics of AGO/sRNA complexes and gain a better understanding of the regulation of targets throughout development, either in a whole animal or a tissue/cell-type-specific manner.

Several studies have analyzed cell-type-specific functions of AGO/sRNA pathways. However, most genomic studies on *C. elegans* AGOs and sRNAs used whole worms to obtain sufficient material for IP/sRNA sequencing, and mainly considered two tissue types (soma and germline) using mutants and subtractive approaches. While using whole worms enables a broad overview of AGO/sRNA targets, it may miss low abundance sRNAs that could participate in cell-type specific functions. We are now able to identify AGO complexes and pools of sRNAs in specific cells or tissues with low amounts of starting material, and can use tissue-specific promoters and 3′ UTRs to drive AGO expression in specific cells and tissues. Furthermore, functional assays, such as reporter assays, are growing increasingly more precise, and coupling these with auxin-degron mediated AGO depletion (Zhang *et al*., 2015) will allow for enhanced control over AGO activity in specific tissues. Our analysis of expression patterns provides an atlas of AGO expression. This and the phenotypes we have uncovered point to specific tissues of interest, and will help prioritize specific cells and tissues for subsequent analyses.

### Stress reveals phenotypes

*C. elegans* laboratory culture conditions are chosen to minimize stress and promote growth. The natural environment for *C. elegans* presents a much more challenging set of conditions to which the worm must continually adapt. Previous studies did not observe phenotypes for most single *ago* mutants under normal laboratory culture conditions (Yigit *et al*., 2006). However, one major function of sRNA pathways is to regulate gene expression to ensure robustness against stressful conditions, with a growing body of literature demonstrating that sRNA pools are altered in response to changes in the environment (Rechavi, Minevich and Hobert, 2011; Rechavi *et al*., 2014; Schott, Yanai and Hunter, 2014; Ni *et al*., 2016; Moore, Kaletsky and Murphy, 2019; Ewe *et al*., 2020; Kaletsky *et al*., 2020; Houri-Zeevi *et al*., 2021). For example, in sRNA pathway and germ granule mutants, elevated temperature strongly affects *C. elegans* fertility, leading to a Mrt phenotype (Ahmed and Hodgkin, 2000; Sundby, Molnar and Claycomb, 2021). While several *agos* had been associated with the Mrt phenotype previously, our systematic analysis revealed additional AGOs whose loss also contributed to reduced fertility under temperature stress, and implicated all types of worm sRNA pathways in this process. Therefore, the molecular mechanisms underlying this phenotype may be different for each *ago* mutant. Further in-depth characterization of germline development and gene expression in the *ago* mutants will be necessary to better understand this phenomenon. However, the potential for indirect effects due to mis-regulation of large groups of genes remains a challenge to disentangle.

Pathogens are another set of stressors that worms are frequently exposed to in the wild, but rarely encounter in the lab. Our sequencing analysis of SAGO-2 and PPW-1 bound sRNAs revealed that these AGOs regulate immunity and pathogen response genes, which led us to test their roles in response to various pathogens. Our studies revealed a dichotomy for two closely related paralogs. These two AGOs share greater than 98% sequence identity at the nucleotide level and the same expression pattern in the intestine, yet PPW-1 is also expressed constitutively in the germline. Loss of either *sago-2* or *ppw-1* led to enhanced survival when confronted with the bacterial pathogen PA14, yet loss of *sago-2* alone resulted in increased Orsay virus infection. We suspect this is because SAGO-2 and PPW-1 regulate immune responsive genes, and their loss is likely to cause mis-regulation of these genes. This may indirectly provide the worms with an enhanced ability to ward off infection by PA14. On the other hand, SAGO-2 is likely to be directly involved in the antiviral response, downstream of RDE-1, while PPW-1 is either redundantly required or dispensable for this response. These two AGOs display different phenotypes, yet vary at only twelve amino acids, and challenging us to understand the molecular mechanisms by which these AGOs act.

While we do not know all the conditions under which AGO/sRNA pathways are required, the worm’s natural environment may provide important clues and experimental contexts for further analyzing the roles of the AGOs. This, in combination with the AGO expression profiles and sRNA sequencing data described here will help define environmental stressors and the functions of sRNA pathways in adapting to them.

### Noncoding RNA targets of sRNA pathways

Transposable elements are tightly regulated in the germline. In many animals, transposable elements are silenced by the piRNA pathway. However, in worms the piRNA pathway directly regulates only a handful of DNA (cut and paste) transposable elements. Instead, the WAGO cluster, including HRDE-1, WAGO-1, PPW-1, and PPW-2 is responsible for nearly all transposable element regulation in the worm. These AGOs likely function constitutively in the germline, while a handful of transposable elements are regulated during spermatogenesis by ALG-3/4.

While sRNA regulation of protein coding genes and transposable elements is well-studied, we show that lincRNAs and pseudogenes are also prominent targets. Both pseudogenes and lincRNAs appear to be regulated by various AGO clusters, implying tissue-specific regulation. For instance, the WAGO cluster targets germline lincRNAs, the ALG-3/4 cluster targets spermatogenesis lincRNAs, and the ERGO-1 cluster targets somatic lincRNAs, when integrating sRNA enrichment and AGO expression data. Both lincRNAs and pseudogenes have the capacity to regulate gene expression themselves (Pink *et al*., 2011; Statello *et al*., 2021), and warrant further study as sRNA targets.

### AGO isoforms and differential functions

Recent studies on two CSR-1 isoforms demonstrated that different isoforms encoded from a single *ago* gene can have different expression patterns, sRNA binding partners, and functions (Charlesworth *et al*., 2021; Nguyen and Phillips, 2021). SAGO-2, PPW-1, ALG-1, ALG-2, and ERGO-1 also exhibit the potential to encode more than one protein, with one or more exons that vary between isoforms. For all but SAGO-2, the differential exons are encoded at the 5′ end of the gene, leading to N-terminal differences in encoded proteins. The reagents we generated tag both isoforms of ALG-1, ALG-2, and SAGO-2. However, we have only tagged the longest isoforms of ERGO-1 and PPW-1. Future studies will be needed to identify and characterize any additional functions of distinct AGO isoforms.

## Conclusion

Our work provides a framework for understanding the complete portrait of sRNA biology in the worm, which is a longstanding model for sRNA research. Our studies point to tissue-specific roles for AGOs in regulating particular facets of the genome, highlight networks of AGO function, and reveal novel, stress linked phenotypes when these pathways are lost. This knowledge provides a basis for elaborating detailed mechanistic insights and opens new avenues of research into AGO and sRNA function.

## Acknowledgements

We are grateful to the Toronto *C. elegans* community for various reagents and helpful discussions, and the lab of Dr. Thomas Hurd for the use of their confocal microscope. We thank members of the Claycomb and Reinke labs and Dr. Guy Riddihough from Life Sciences Editors for feedback on the manuscript.

Funding sources: This work was funded by Canadian Institutes of Health Research Project Grants PJT-156083, PJG-175378, PJT-178076, and PJT-400784 (AWR) and Natural Sciences and Engineering Research Council of Canada Discovery Grant RGPIN-2020-06235 to JMC. JMC is a Canada Research Chair Tier II in Small RNA Biology. AWR is supported by an Alfred P. Sloan Research Fellowship FG2019-12040. Some strains were provided by the CGC, which is funded by NIH Office of Research Infrastructure Programs [P40 OD010440].

## Author Contributions

Conceptualization: JMC, US; Data curation: US, AL, LW; Formal Analysis: US, AL, LW; Funding acquisition: JMC, AWR; Investigation: US, RXL, ARW, WZ, AES; Methodology: US, JMC; Project administration: JMC, US; Resources: JMC, AWR; Software: US, AL, LW; Supervision: JMC; Validation: US, RXL, ARW, WZ; Visualization: US, AGC; Writing: US, JMC.

## Materials and Methods

### Contact for Reagent and Resource Sharing

Further information and requests for resources and reagents should be directed to and will be fulfilled by the Lead Contact, Julie Claycomb.

### Experimental Models and Subject Details

#### *C. elegans* strains

A complete list of strains used in this study is provided in Table S8. The Bristol strain N2 was used as the reference strain.

#### Nematode growth

All strains were maintained at 20°C unless otherwise indicated on 3.5cm plates containing Nematode Growth Medium (NGM) seeded with *E. coli* OP50 bacteria as a food source (Brenner, 1974).

### Method Details

#### Phylogenetic tree construction

Protein sequences of Argonautes were aligned using MUSCLE with default settings (Madeira *et al*., 2019). Evolutionary analyses were conducted in MEGA X (Kumar *et al*., 2018). The evolutionary history was inferred by using the Maximum Likelihood method. The initial tree for the heuristic search was obtained by applying Neighbor-Join and BioNJ algorithms to a matrix of pairwise distances estimated using a JTT model, and then selecting the topology with superior log likelihood value. A discrete Gamma distribution was used to model evolutionary rate differences among sites (2 categories (+G, parameter = 1.9757)). The rate variation model allowed for some sites to be evolutionarily invariable ([+I], 0.62% sites). The bootstrap consensus tree was inferred from 1000 replicates. Branches corresponding to partitions reproduced in less than 50% bootstrap replicates are collapsed.

#### CRISPR/Cas9 genome editing

Tagging genes with GFP::3xFLAG was conducted as previously described (Dickinson *et al*., 2015). Single guide RNAs (sgRNAs) were designed using CRISPOR (Concordet and Haeussler, 2018) and cloned into pDD162 via site directed mutagenesis PCR (see Table S7 for primers used). For repair templates, homology arms of 500-700bp on either side of the insertion site were amplified using Q5 High Fidelity Polymerase from N2 genomic DNA and cloned into pDD282 cut with ClaI and SpeI restriction sites using NEBuilder HiFi Assembly mix. The homology arms for *ppw-1* and *sago-2* were amplified from *sago-2(tm894)* and *ppw-1(tm914)* genomic DNA respectively since these harbor deletions allowing for the design of primers that will specifically amplify the intended genomic regions (*ppw-1* and *sago-2* are highly similar in sequence). If the sgRNA site was not destroyed by the insertion of the repair template, synonymous mutations were introduced into the PAM sequence or 3-4 synonymous mutations were introduced into the sgRNA sequence (see Table S7 for primers used). Inserts of all cloned plasmids were verified by Sanger sequencing. An injection mix was used as follows: 10ng/µl repair template, 50ng/µl sgRNA, 10ng/µl pGH8, 5ng/µl pCFJ104, 2.5ng/µl pCFJ90.

Tagging genes with 3xFLAG alone was conducted as previously described (Dokshin *et al*., 2018). Single guide RNAs (sgRNAs) were designed using CRISPOR (Concordet and Haeussler, 2018). *S. pyogenes* Cas9 protein and guide RNAs (tracrRNA and crRNA) were ordered from IDT. The 3xFLAG repair template was ordered from IDT as an ultramer with ∼35bp of homology arms flanking the insertion site (see Table S7): 5′ 35bp-flanking-region-GATTATAAAGACGATGACGATAAGCGTGACTACAAGGACGACGACGACAAGCGTGATTACAAGGATGACGATGACAAGAGA-35bp-flanking-region 3′. The pRF4 *rol-6(su1006)* injection marker was used to screen for successfully injected worms. These were screened via PCR flanking the intended insertion site to search for integrations. An injection mix was used as follows: Cas9 250ng/µl, tracrRNA 100ng/µl, crRNA 56ng/µl, 220ng/µl repair template, 20ng/µl pRF4. The 3xFLAG::WAGO-1 strain was generated as described for the GFP::3xFLAG procedure but the GFP sequence was cloned out of the pDD282 plasmid, leaving only the 3xFLAG (see Table S7 for primers used).

To generate indels, sgRNAs spanning a genomic region were designed and injected. Mutation of the *dpy-10(cn64)* gene at the same time was used as a co-CRISPR marker to identify and enrich candidate editing events (Arribere *et al*., 2014). An injection mix was used as follows: 20ng/µl *dpy-10* conversion template, 50ng/µl sgRNA, 10ng/µl pGH8, 5ng/µl pCFJ104, 2.5ng/µl pCFJ90. Candidate mutants were screened via PCR spanning the genomic region to be excised. This method was used to generate the *wago-10(tor133)* allele which deletes the region between the 695nt and the 2394nt (1699bp deletion).

To generate single nucleotide polymorphisms a similar approach to generating 3xFLAG insertions was used where the repair template oligo was ∼100bp of the genomic sequence with mutations to insert with the sgRNA cut site in the middle. With this approach the *sago-2(tor135)* allele was generated where the start methionine and 8^th^ amino acid were changed to stop codons (see Table S7).

#### Brood size assays

Five L4 hermaphrodites were transferred to a 15mm NGM plate seeded with OP50 and allowed to propagate at the desired temperature (20°C or 25°C). The progeny of these animals were used in the brood size assay. An Individual L4 hermaphrodite was transferred to a 15mm plate and transferred to a fresh plate every day until egg laying ceased (typically 3 days at 25°C and 4 days at 20°C). The hatched progeny were counted. At least 10 hermaphrodites were assayed per strain.

#### Mortal germline assays

The assays were performed similarly to (Ahmed and Hodgkin, 2000). Five L4 worms were picked to five individual plates to establish five lines and incubated at 25°C. Each generation (every 3 days) five L4 worms were picked from each plate to a new plate. A line was considered mortal if there were no more progeny to pick five L4s from.

#### RNAi

RNAi by feeding was conducted as described (Ahringer, 2006). Three L4 worms were placed on NGM plates supplemented with 25µg/ml carbenicillin and 1mM IPTG and seeded with the specific RNAi bacterial strain. The bacteria were grown overnight in LB supplemented with 100µg/ml carbenicillin. The progeny of these worms were tested for the expected RNAi phenotype.

#### PA14 survival assays

*Pseudomonas aeruginosa* (PA14) was streaked on standard LB plates supplemented with carbenicillin at 100μg/ml and grown overnight. Single colonies were picked and grown in 3ml of LB overnight culture. 20μl of PA14 was seeded on 3.5cm slow killing (SK) NGM plates as previously described (Tan, Mahajan-Miklos and Ausubel, 1999). These SK plates were subsequently incubated overnight at 37°C and then equilibrated for two days at 25°C. All strains used for the PA14 survival assay were grown to gravid adults on 3.5cm NGM plates at 20°C and bleached. The progeny that survived bleaching were then grown to the L4-stage on NGM plates at 20°C. 50 L4s were then plated on SK plates in technical triplicates and subsequently moved to 25°C. Worms were transferred to new SK plates every 24 hours. Counts of the number of dead worm carcasses was performed after 48 hours prior to transferring and performing a final count of both dead worm carcasses and live worms after 72 hours.

#### Orsay Virus infection assay

Orsay virus filtrate was prepared as previously described (Bakowski et al., 2014). Plates of Orsay virus–infected worms were maintained until starvation. Virus from infected worms was collected by washing plates with M9, passing through 0.22μm filters (Millipore Sigma), and stored at −80°C. Next, ∼1000 L1 worms were mixed with 100μl of 10×OP50-1 and 500μl of the viral filtrate and then plated on 6cm NGM plates. At 16 hpi (hours post infection), animals were fixed and fluorescent in situ hybridization (FISH)-stained to assess infection status. Worms were fixed in 1ml of 4% paraformaldehyde (PFA) in PBS containing 0.1% Tween20 (PBST), for 20 min at RT or overnight at −20°C. Worms were then washed once in 1 ml hybridization buffer [900 mM NaCl, 20 mM Tris (pH 8.0), and 0.01% SDS] and incubated overnight at 46°C in 100μl hybridization buffer containing FISH probe (5 to 10 ng/μl) conjugated to a Cal Fluor 610 dye (LGC Biosearch Technologies). Orsay Probe 1 (gacatatgtgatgccgagac) and Orsay Probe 2 (gtagtgtcattgtaggcagc) were mixed at a 50:50 (10 ng/μl) ratio and used to detect Orsay virus. Stained animals were washed once in 1ml wash buffer (hybridization buffer containing 5mM EDTA) and incubated in 500μl fresh wash buffer for a further 30 min at 46°C. Worms were resuspended in 20μl EverBrite Mounting Medium (Biotium) and mounted on slides for imaging. Worms with any number of cells stained with the FISH probes were considered infected.

#### Protein lysate preparation

Synchronous populations of ∼100,000 L1 worms were plated on 15 cm NGM plates with ∼2ml of 5x concentrated OP50 *E. coli* as a food source. Five of these plates were used as starting material for protein isolation. Worms were grown for 48h for L4 staged worms or 58h for young adults (worms that had transitioned to producing oocytes but not yet with embryos). Worms were washed three times with M9 buffer (22mM KH_2_PO_4_, 42mM Na_2_HPO_4_, 86mM NaCl) and one time with EDTA buffer (10% glycerol, 10mM EDTA, 30mM HEPES, 100mM Potassium Acetate). The pellet was flash frozen in a dry ice/ethanol bath. The frozen pellets were stored at – 80°C.

The frozen pellet was resuspended 1.5:1 (v/v) in ice-cold IP buffer (10% glycerol, 10mM EDTA, 30mM HEPES, 100mM Potassium Acetate, 2mM DTT, 0.1% NP-40) supplemented with protease and phosphatase inhibitors (1 tablet per 5ml buffer of cOmplete™, mini, EDTA-free protease inhibitor cocktail [Roche], 1:100 phosphatase inhibitor cocktail 2 [Sigma], 1:100 phosphatase inhibitor cocktail 3 [Sigma]). If the pellet was to be used for RNA purification, 1% (v/v) SuperaseIN RNAse inhibitor (ThermoFisher) was added. The pellet was homogenized using a stainless steel dounce homogenizer (Wheaton Incorporated) until intact worms were no longer visible. Extracts were centrifuged at 13,000g for 10 min at 4°C, and the supernatant transferred to a fresh tube. A Lowry assay (Lowry et al., 1951) was performed to determine total protein concentration using a Nanodrop 1800C spectrophotometer (ThermoFisher).

#### Immunoprecipitation of Argonaute complexes

All IPs were conducted on 5mg of total protein per reaction. Input control samples were made by taking 10% of the lysate before the addition of antibodies. For the IP of GFP tagged Argonautes, GFP-Trap_MA beads (ChromoTek) were equilibrated by washing them three times in 1ml of IP buffer. 20μl of beads were added to 5mg total protein in a reaction volume of 500μl and rotated at 4°C for an hour. The beads were then washed four times with 1ml of IP buffer, separated from the supernatant on a magnetic stand, and rotated for 10 min at 4°C between each wash. For the IP of 3xFLAG tagged Argonautes, Monoclonal Anti-FLAG M2 (Sigma-Aldrich) were bound to Dynabeads™ Protein G (ThermoFisher) or GB-Magic Protein A/G Immunoprecipitation Magnetic Beads according to manufacturer’s instructions. 5μg of Anti-FLAG M2 in 200μl of PBS-T were added to 50μl of Dynabeads and rotated at room temperature for 10 min. Then IPs were conducted as described above except the whole 50μl of ANTI-FLAG M2 bound beads were used in the IP.

For small RNA sequencing, three to six IP reactions were combined in 200μl of IP buffer. 800μl of TRI Reagent (Molecular Research Centre) was added and the samples were frozen at -80°C until RNA extraction and sequencing were done as described below.

#### Western blot analysis

Proteins were resolved by SDS-PAGE on precast gradient gels (4-12% Bis-Tris Bolt gels, ThermoFisher) and transferred to Hybond-C membrane (Amersham Biosciences) using a Bio-Rad semi-dry transfer apparatus at 25V for 45min. The membrane was washed three times with PBST (137mM NaCl, 2.7mM KCl, 10mM Na_2_HPO_4_, 1.8mM KH_2_PO_4_ pH 7.4, 0.1% Tween-20) and blocked with 5% milk-PBST (PBST, 5% skim milk) for 1h at room temperature. The membrane was then incubated overnight at 4°C with 1:2000 M2 Anti-FLAG antibody (Sigma-Aldrich) in 5% milk-PBST. The membrane was washed three times with PBST and then blocked with 5% milk-PBST for 30 minutes at room temperature. The membrane was incubated with 1:1000 anti-mouse IgG HRP-linked antibody (Cell Signaling Technology) in 5% milk-PBST for 1 hour at room temperature. The membrane was washed three times in PBST and then developed by using Luminata Forte Western HRP substrate (Millipore).

#### RNA isolation

RNA was isolated using TRI Reagent (Molecular Research Centre). Samples were mixed with TRI Reagent in a 1:4 ratio and frozen at -80°C. Samples were vortexed at room temperature for 15min and frozen again at -80°C for 15min. This was repeated for a total of three times. 100μl of chloroform was then added to the samples and centrifuged at 12,000g for 15min at 4°C. The top aqueous phase was transferred to a fresh tube. Phenol:chloroform:isoamyl alcohol (Sigma-Aldrich) was added in a 1:1 ratio and centrifuged at 12,000g for 15min at 4°C. The top aqueous phase was transferred to a fresh tube. 20μg of glycogen (Ambion) and a 1:1 ratio of isopropanol was added to the samples and incubated at -20°C for 30min. Samples were then centrifuged at 16,000g for 30min at 4°C and the supernatant was removed. The pellet was washed with 900μl of 70% ice-cold ethanol for 10min and centrifuged at 16,000g for 10min at 4°C. This was repeated twice. The pellet was then left to air dry for 10min at room temperature and then resuspended in 6-25μl of RNase free water preheated to 70°C.

#### Small RNA library preparation and sequencing

Small RNA libraries were prepared with the NEBNext Small RNA Library Prep Set for Illumina (New England Biolabs) following the protocol provided by the manufacturer. 1μg of total RNA or immunoprecipitated RNA was used as starting material. The resulting DNA library was visualized using 8% PAGE and bands corresponding to small RNAs of size range 16-30bp (∼135-150bp on gel) were excised. The DNA was eluted from the excised bands by rotating overnight in 500μl of DNA Gel Elution buffer at room temperature. The DNA was precipitated with 20μg glycogen (Ambion), 50μl of 3M Sodium Acetate pH 5.2 and 1 volume of isopropanol (Sigma-Aldrich) as described above and ultimately resuspended in 12μl of Ultra-Pure water. The library DNA was quantified using a Qubit HS DNA kit (ThermoFisher). Between 12-19 Libraries were pooled in equal amounts and sequenced on a HiSeq 2500 Sequencing System (Illumina).

#### Small RNA sequencing analysis

The small RNA sequences obtained from the sequencer were first assessed for quality using FastQC (version 0.11.5, (Andrews, 2010)). Adapter sequences were then removed using cutadapt (version 1.15, (Martin, 2011)) using the following command: -a ADAPTER -f fastq -m 16 -M 30 --discard-untrimmed.

The sequences were then run through FastQC again to assess quality. The trimmed reads were then aligned to the C. elegans PRJNA13758 ce11 genome assembly (WormBase version WS276) with STAR (version 2.6.0c, (Dobin et al., 2013)) using the following commands: --runThreadN 12 --outSAMtype BAM SortedByCoordinate --outFilterMultimapNmax 50 --outFilterMultimapScoreRange 0 -- outFilterMismatchNoverLmax 0.05 --outFilterMatchNmin 16 -- outFilterScoreMinOverLread 0 --outFilterMatchNminOverLread 0 --alignIntronMax 1.

The reads were then counted using a custom R (version 3.6.3) script against publicly available genome annotations. The WormBase version WS276 PRJNA13758 ce11 canonical geneset annotations were used (excluding miRNAs, repeats and transposons). *C. elegans* miRNA annotations were obtained from miRBase (release 22.1). For repeats and transposons RepeatMasker + Dfam (ce10 - Oct 2010 - RepeatMasker open-4.0.6 - Dfam 2.0) annotations were used. The UCSC Lift Genome Annotations tool was used to convert ce10 coordinates to ce11 coordinates. Briefly, the counting script used the findOverlaps function from the GenomicAlignments package (version 1.22.1) to assign reads to features. Multiple aligning reads or reads that align to more than one feature were dealt with by counting reads in a sequential manner to the different gene biotypes in the following order (AS stands for antisense): miRNA, piRNA, rRNA, snoRNA, snRNA, tRNA, ncRNA, lincRNA, repeats AS, protein coding AS, pseudogene AS, lincRNA AS, antisense RNA, rRNA AS, snoRNA AS, snRNA AS, tRNA AS, ncRNA AS, miRNA AS, piRNA AS, protein coding, pseudogene, antisense RNA AS, repeats. For reads that align to more than one feature in the same biotype group, the read count was split between the features based on the fraction of uniquely aligned reads to each of those features (unique weighing). Subsequent analysis was performed using custom R scripts (https://github.com/ClaycombLab/Seroussi_2022).

To determine small RNAs enriched in Argonaute IPs, reads were first normalized to library size (reads per million - RPM). We used two approaches for this. In the first approach we normalized the reads to the entire library size. In the second approach, we normalized the reads to library size minus sense reads of: rRNA, snoRNA, snRNA, tRNA, ncRNA, lincRNA, protein coding, and pseudogene. These likely represent RNA degradation products so removing them may eliminate noise. The first approach was used in the initial analysis of small RNAs associated with Argonaute IPs for complete transparency and unbiased assignment of small RNA types and targets. Indeed, the vast majority of reads in all libraries were antisense rather than sense. Thus, subsequent enrichment analysis and comparisons between Argonaute IPs and identification of likely targets used the second approach. To determine whether small RNAs against a particular target were enriched in an Argonaute IP, the following calculation was made: Enrichment = IP RPM + 0.01 / Input RPM + 0.01. The 0.01 represents a pseudocount to eliminate the possibility of dividing by zero. A target was considered enriched if in every replicate it was at least 2-fold enriched and had at least 5 RPM in the IP replicates. To further refine the analysis, where indicated in the text, only 22G or 26G-RNAs were used to calculate enrichment. These were defined as reads of 20-24nt and 25-27nt respectively with no 5’ nucleotide constraints.

To determine differential expression of small RNAs in mutant *argonaute* strains, we used the R package DESeq2 (version 3.14 (Love, Huber and Anders, 2014)).

Published datasets were used as follows: WAGO-1 small RNA targets and glp-4 enriched/depleted small RNA targets (Gu et al., 2009), ERGO-1 enriched small RNA targets (Vasale et al., 2010), *alg-3; alg-4* depleted small RNA targets (Conine et al., 2010), CSR-1 enriched small RNA targets (Claycomb et al., 2009), *mut-16* depleted small RNA targets (Phillips et al., 2012) defined as two-fold depleted in mutant and having at least 10RPM, gamete specific expressed genes (Ortiz et al., 2014), RdRP mutants-depleted small RNA targets were reanalyzed as described above and defined as two-fold depleted in mutant and having at least 5RPM (Sapetschnig et al., 2015). All published gene lists used were converted to WS276 gene names using WormBase Converter (Engelmann et al., 2011) before being compared to gene lists generated in this study.

Gene Ontology (GO) analysis was performed using gProfiler and Wormbase Enrichment Suite (Angeles-Albores *et al*., 2016, 2018) (Table S7).

#### Microscopy

All images were taken on a Leica DMi8 TCS SP8 confocal microscope, except for those in Fig. 7B, which were taken on a Nikon TiE microscope with a C2 confocal module. All images presented are a single 0.4μm slice, taken using a 488nm laser, and in most instances Normarski images are also displayed. Images were processed for figures using FIJI, Adobe Photoshop, and Adobe Illustrator.

#### Quantification of Argonaute and germ granule factor overlap

Staged GFP::3xFLAG::AGO expressing worms were washed in M9 and immobilized on positively charged glass slides with 10μL of 10mM Levamisole. Germlines were dissected with a 17-gauge needle, by cutting at the vulva or head/tail. A coverslip was added, and the slides were placed on a flat aluminum block in dry ice for at least 10 minutes. Slides were either kept at -80°C or fixed immediately. The coverslip was popped off with a razor blade and samples were fixed at -20°C for five min in each of, 100% methanol, 50/50 methanol/acetone, and 100% acetone. Samples were air dried, and a hydrophobic marker was used to outline the sample. All washes and incubations were performed in a humidity chamber (i.e. a lidded plastic tray covered in aluminum foil with wet paper towels and a plastic rack to hold the slides). Samples were washed 2 x 5 minutes with PBST, then blocked for 30 minutes at room temperature with PBST+BSA (1x PBS, 0.1% Tween-20 and 3% BSA). Samples were then incubated with primary antibodies (Anti-HA [Sigma] or Anti-PGL) overnight at 4°C. Slides were washed 3 x 10 minutes with PBST, then blocked with PBST+BSA for 30min at room temperature. Samples were incubated with secondary antibodies (Anti-rat::TRITC or anti-mouse IgM::TRITC) for 1h at room temperature then washed 3 x 10min with PBST and 3 x 5 minutes with PBS. Samples were stained with DAPI (1μg/ml) for 10 minutes then washed 3 x 5min with PBS and mounted in 2μl of Vectashield (Vector labs). Samples were kept at -20°C until imaged.

Colocalization of proteins was calculated with the ImageJ plugin JaCoP (Bolte and Cordelieres, 2006). One germline from each of six different animals was imaged per strain and developmental time point. Regions of interest (ROI) were generated using the 3D objects counter plugin in ImageJ (Schneider, Rasband and Eliceiri, 2012) by adjusting the threshold until only germ granule pixels are detected. Five Z-stacks (0.9 μM apart) were quantified per germline to capture the overlap over a 3D space. Mander’s co-localization coefficients are calculated using JaCoP (Bolte and Cordelières, 2006).

### Data And Software Availability

All high-throughput sequencing data are available through GEO, accession number GSE208702. Custom R scripts are available via GitHub: https://github.com/ClaycombLab/Seroussi_2022.

### Key Resources Table

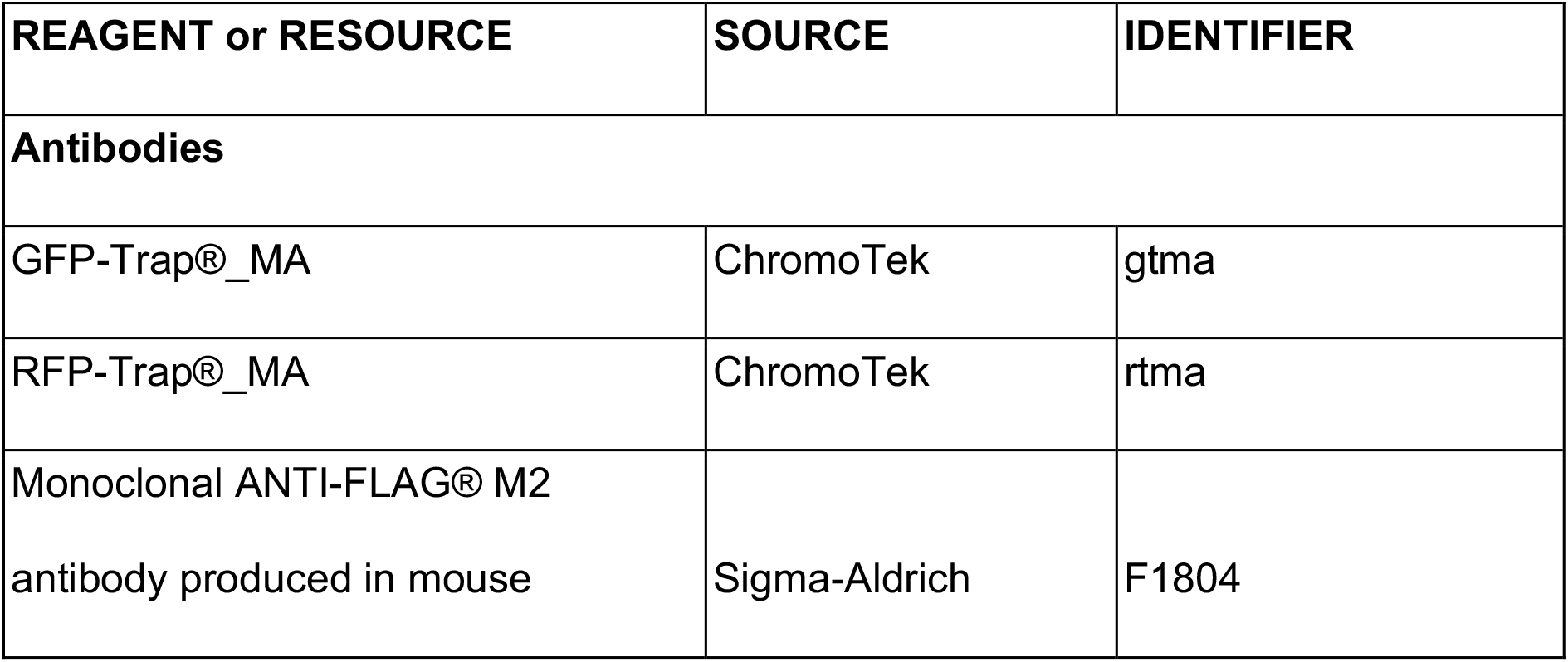
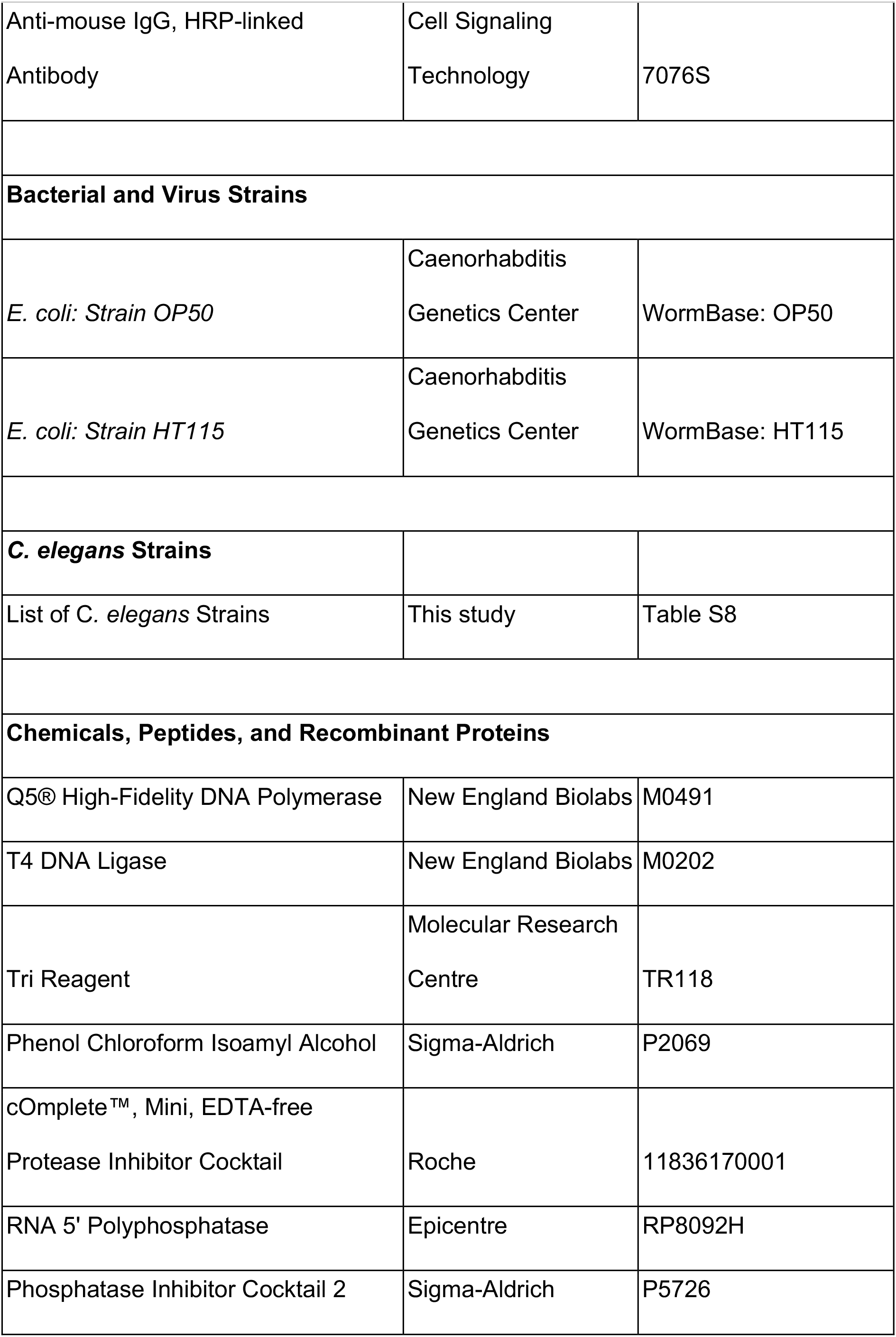
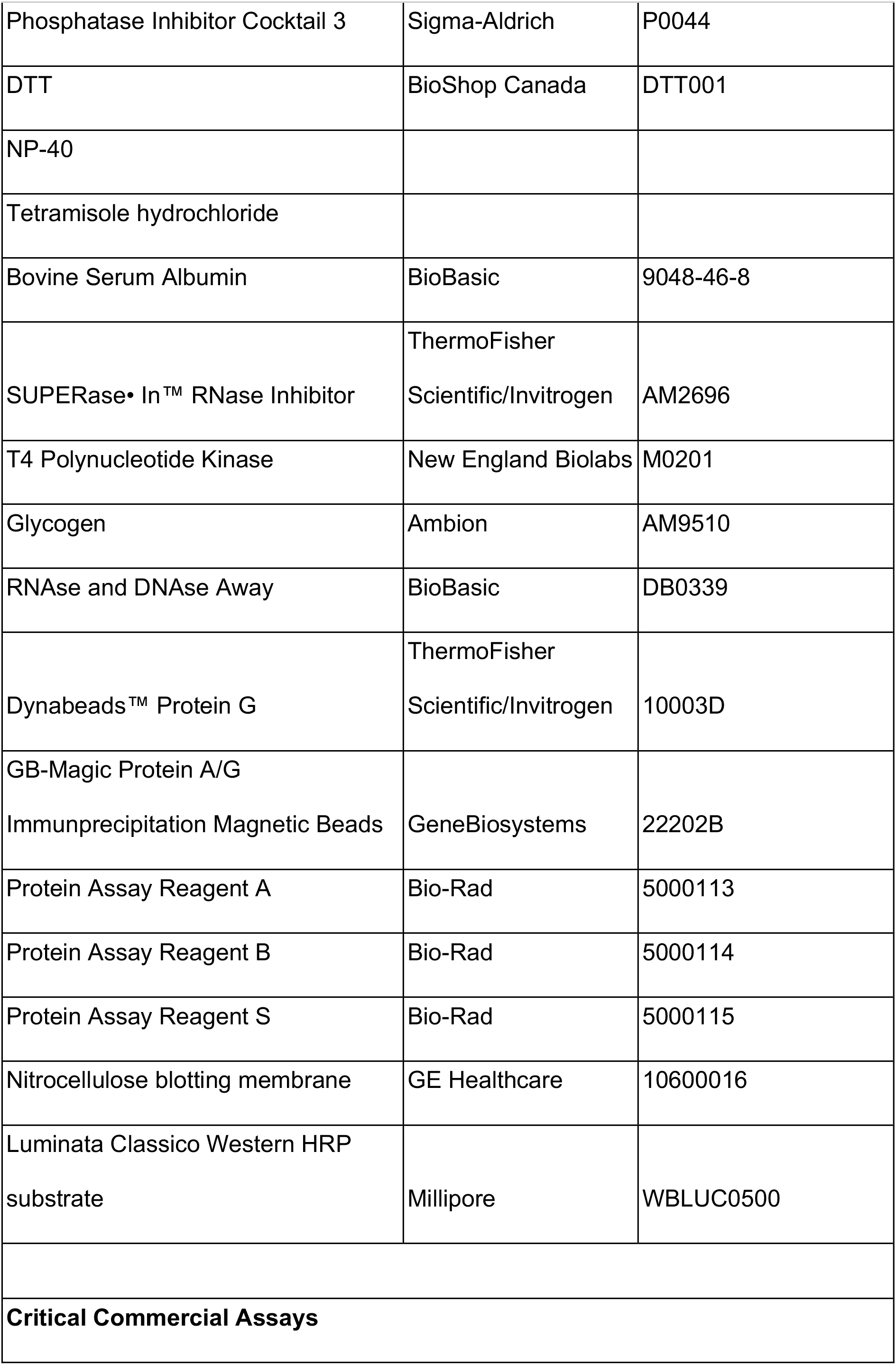
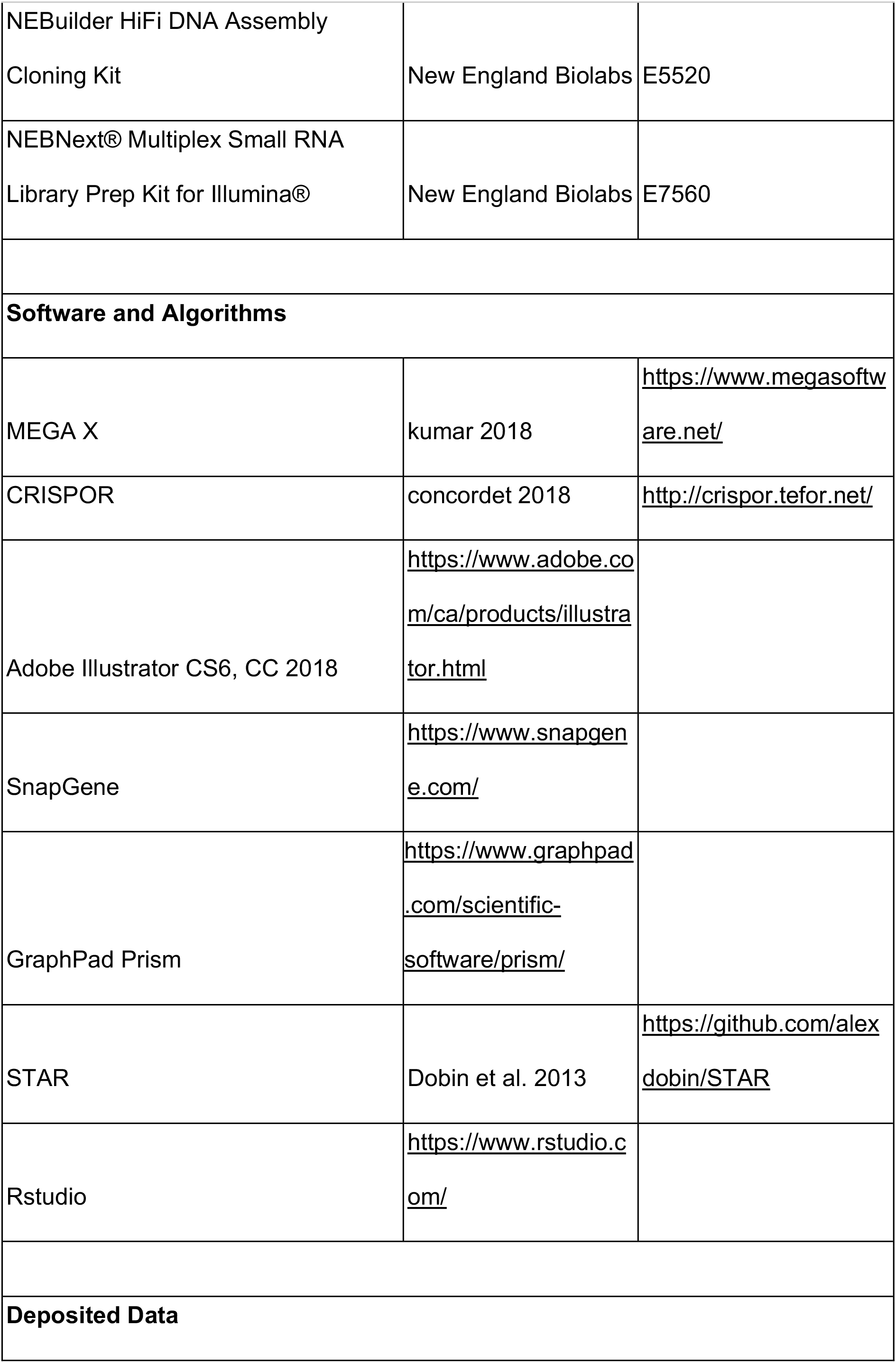
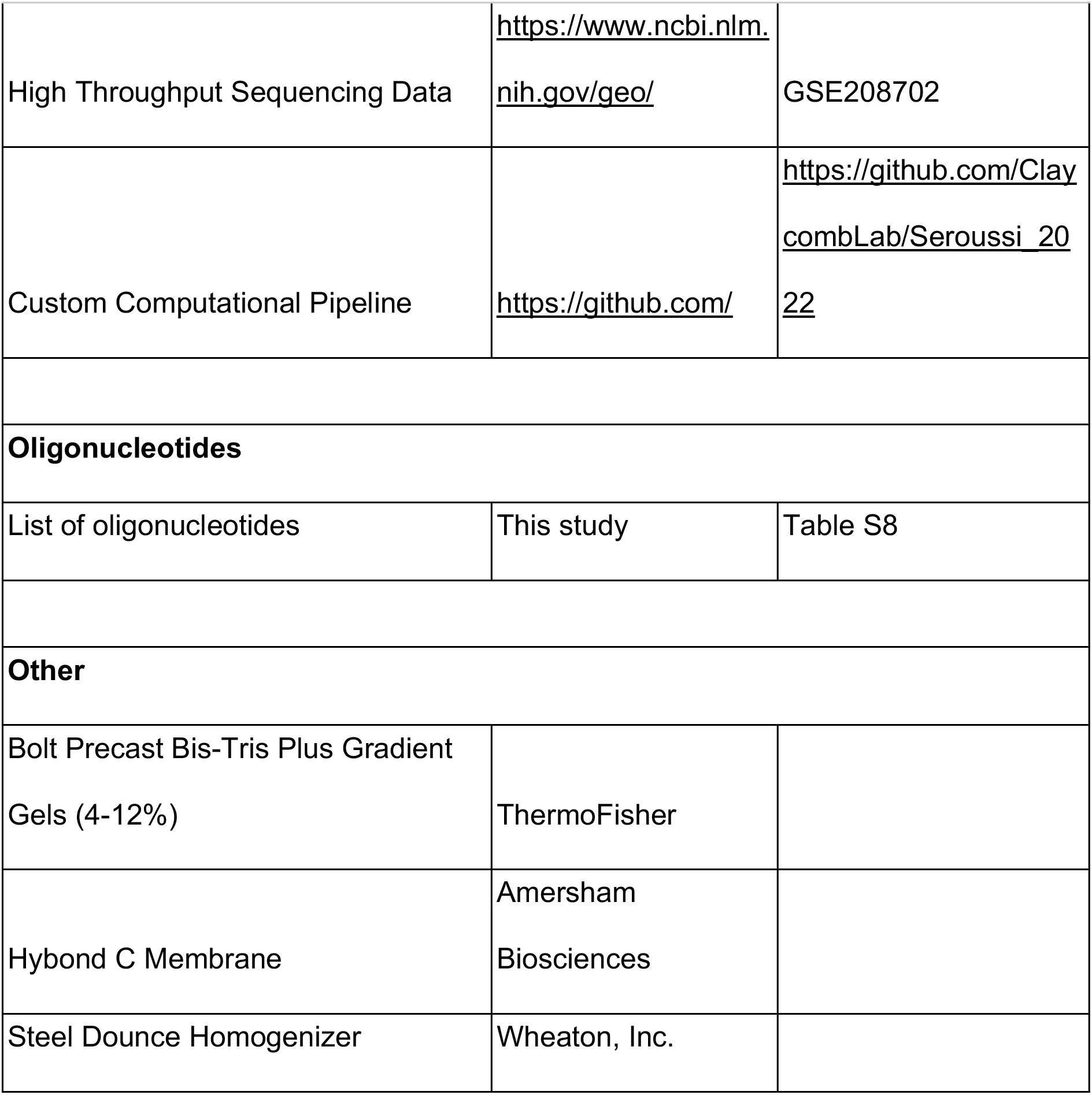

| REAGENT or RESOURCE | SOURCE | IDENTIFIER |
| --- | --- | --- |
| **Antibodies** |  |  |
| GFP-Trap®_MA | ChromoTek | gtma |
| RFP-Trap®_MA | ChromoTek | rtma |
| Monoclonal ANTI-FLAG® M2 antibody produced in mouse | Sigma-Aldrich | F1804 |
| Anti-mouse IgG, HRP-linked Antibody | Cell Signaling Technology | 7076S |
| **Bacterial and Virus Strains** |  |  |
| *E. coli: Strain OP50* | Caenorhabditis Genetics Center | WormBase: OP50 |
| *E. coli: Strain HT115* | Caenorhabditis Genetics Center | WormBase: HT115 |
| **C. elegans Strains** |  |  |
| List of *C. elegans* Strains | This study | Table S8 |
| **Chemicals, Peptides, and Recombinant Proteins** |  |  |
| Q5® High-Fidelity DNA Polymerase | New England Biolabs | M0491 |
| T4 DNA Ligase | New England Biolabs | M0202 |
| Tri Reagent | Molecular Research Centre | TR118 |
| Phenol Chloroform Isoamyl Alcohol | Sigma-Aldrich | P2069 |
| cOmplete™, Mini, EDTA-free Protease Inhibitor Cocktail | Roche | 11836170001 |
| RNA 5' Polyphosphatase | Epicentre | RP8092H |
| Phosphatase Inhibitor Cocktail 2 | Sigma-Aldrich | P5726 |
| Phosphatase Inhibitor Cocktail 3 | Sigma-Aldrich | P0044 |
| DTT | BioShop Canada | DTT001 |
| NP-40 |  |  |
| Tetramisole hydrochloride |  |  |
| Bovine Serum Albumin | BioBasic | 9048-46-8 |
| SUPERase• In™ RNase Inhibitor | ThermoFisher<br>Scientific/Invitrogen | AM2696 |
| T4 Polynucleotide Kinase | New England Biolabs | M0201 |
| Glycogen | Ambion | AM9510 |
| RNase and DNase Away | BioBasic | DB0339 |
| Dynabeads™ Protein G | ThermoFisher<br>Scientific/Invitrogen | 10003D |
| GB-Magic Protein A/G<br>Immunoprecipitation Magnetic Beads | GeneBiosystems | 22202B |
| Protein Assay Reagent A | Bio-Rad | 5000113 |
| Protein Assay Reagent B | Bio-Rad | 5000114 |
| Protein Assay Reagent S | Bio-Rad | 5000115 |
| Nitrocellulose blotting membrane | GE Healthcare | 10600016 |
| Luminata Classico Western HRP<br>substrate | Millipore | WBLUC0500 |
| **Critical Commercial Assays** |  |  |
| NEBuilder HiFi DNA Assembly Cloning Kit | New England Biolabs | E5520 |
| NEBNext® Multiplex Small RNA Library Prep Kit for Illumina® | New England Biolabs | E7560 |
| **Software and Algorithms** |  |  |
| MEGA X | kumar 2018 | https://www.megasoftware.net/ |
| CRISPOR | concordet 2018 | http://crispor.tefor.net/ |
| Adobe Illustrator CS6, CC 2018 | https://www.adobe.com/ca/products/illustrator.html |  |
| SnapGene | https://www.snapgene.com/ |  |
| GraphPad Prism | https://www.graphpad.com/scientific-software/prism/ |  |
| STAR | Dobin et al. 2013 | https://github.com/alexdobin/STAR |
| Rstudio | https://www.rstudio.com/ |  |
| **Deposited Data** |  |  |
| High Throughput Sequencing Data | https://www.ncbi.nlm.nih.gov/geo/ | GSE208702 |
| Custom Computational Pipeline | https://github.com/ | https://github.com/ClaycombLab/Seroussi_2022 |
| **Oligonucleotides** |  |  |
| List of oligonucleotides | This study | Table S8 |
| **Other** |  |  |
| Bolt Precast Bis-Tris Plus Gradient Gels (4-12%) | ThermoFisher |  |
| Hybond C Membrane | Amersham Biosciences |  |
| Steel Dounce Homogenizer | Wheaton, Inc. |  |

## Supplemental Information

### Supplemental Figures

**Figure S1. Epitope Tag locations and western blot validations**

(A) Gene diagrams of each Argonaute. The green arrow points to the GFP::3xFlag tag insertion site. The number corresponds to the distance in amino acids from the first methionine. The letter corresponds to the isoform if there is more than one. Genotypes of mutants that were used in this study are indicated in red.

(B) Western blots of GFP::3xFLAG::AGO IPs. Inputs are 100µg of total protein lysate. Mock conditions are IPs that were conducted using RFP antibodies.

**Figure S2. Additional analyses of tagged Argonautes**

(A) Gene diagrams of WAGO-5 and WAGO-11 as in Fig S1 (left). GFP driven under the promoter of WAGO-11 (right). GFP is observed in the distal gonadal sheath of L4 hermaphrodites.

(B) Genome browser tracks of mRNA tiling array sequencing at the *wago-5* (left) and *wago-11* (right) loci from modENCODE.

(C) RT-PCR and gel electrophoresis analysis of *gfp::3xflag::wago-5* and *gfp::3xflag::wago-11*. Only the *gfp::3xflag::wago-11* cDNA is detected by PCR.

(D) Genome browser tracks of small RNAs targeting the *wago-5* locus in WAGO-1 IP.

(E) Western blots of 3xFLAG tagged AGO IPs. Inputs are 100ug of total protein lysate. Mock conditions are IPs conducted using non-antibody conjugated beads. 3xFLAG tag insertion sites are at the same position as GFP::3xFLAG tags shown in Figure S1.

**Figure S3. Metagene analysis of sRNAs**

Metagene profiles for the protein coding gene targeting sRNAs enriched in association with each AGO. The number of targets is indicated in parenthesis. To account for differences in the size of targets, each gene was partitioned into 100 bins and the mean coverage in RPM was determined for each bin. Green line = AGO IP, green dashed line = input, red line = *ago* mutant, red dashed line = wild-type.

**Figure S4. Additional analysis of the endo-siRNA binding AGO IPs**

(A) A Venn diagram showing the overlaps of pseudogene targets of the AGO clusters as highlighted by the color scheme in Figure 5A.

(B) A Venn diagram showing the overlaps of transposon targets of the AGO clusters as highlighted by the color scheme in Figure 5A.

(C) Box plots showing the enrichment levels (IP/input) of sRNAs targeting each transposon broken down by transposon family in each AGO IP.

(D) A Venn diagram showing the overlaps of lincRNA targets of the AGO clusters as highlighted by the color scheme in Figure 5A.

(E) Network analysis showing which AGO is enriched for sRNAs targeting another AGO. Arrow direction indicates the direction of regulation; i.e. the AGO at the blunt end regulates the AGO at the pointy arrow end.

**Figure S5. Loss of *ago* genes results in differential effects of sRNA levels associated with each AGO**

Box plots showing the expression levels (y-axis) of the sRNAs that target the protein coding targets for which sRNAs were enriched in each AGO IP (x-axis) in the indicated argonaute mutant (panels). Red dashed lines indicate 2-fold change cutoff.

**Figure S6. AGO expression in embryos**

Images of embryos from left to right: 2 cell stage, 4 cell stage, ∼40 cell stage, ∼200 cell stage (end of gastrulation), comma/bean elongation stage, 3 fold stage/pretzel stage. Scale bar = 50µm.

**Figure S7. AGO expression in L1 larvae**

Images of L1 larvae. Scale bar = 50µm.

**Figure S8. AGO expression in L2 larvae**

Images of L2 larvae. Scale bar = 50µm.

**Figure S9. AGO expression in L3 larvae**

Images of L3 larvae. Scale bar = 50µm.

**Figure S10. AGO expression in L4 larvae**

Images of L4 larvae. Scale bar = 50µm.

**Figure S11. AGO expression in adult worms**

Images of adult worms. Scale bar = 50µm.

**Figure S12. AGO expression in males**

Images of adult males. Scale bar = 50µm.

**Figure S13. Representative images of developmental stages of the RDE-1 transcriptional reporter**

Top row of each panel is DIC + GFP images. Bottom row GFP images. (A) embryos, (B) L1, (C) L2, (D) L3, (E) L4, (F) adult. Scale bar = 50µm.

**Figure S14. Additional analysis of AGO mRNA expression**

(A) mRNA levels of AGOs in different developmental stages. Data is an aggregate of public datasets available on WormBase version WS273.

(B) Illustration of hermaphrodite germline with sections that were sequenced in (Tzur *et al*., 2018).

(C) Table showing in heatmap form the relative expression of each argonaute mRNA in each germline section. Note that *alg-3*, *alg-4*, and *wago-10* were not detected. Note *alg-2*, *ergo-1*, *C04F12.1/vsra-1*, and *nrde-3* show increasing expression in oocyte sections.

**Figure S15. The mortal germline of *wago* mutants is reversible and associated with reduced germline proliferation**

(A) A temperature shift experiment where *wago* mutants were placed at 20°C for one generation, then transferred to 25°C for three generations and then back to 20°C. Brood size was measured every generation (N = 10 per genotype).

(B) Count of nuclei at the mitotic zone (N = 5). *** = *p*-value < 0.001, Two-way ANOVA with Dunnet’s Post Hoc multiple comparison test.

(C) Count of oocytes in germline arms (N >= 22). * = *p-*value < 0.05, *** = *p*-value < 0.001,Two-way ANOVA with Dunnet’s Post Hoc multiple comparison test.

(D) Count of apoptotic nuclei in germlines of acridine orange stained *wago* mutants (N >= 19).

### Supplemental Tables

**Table S1. A table summarizing previously published data on *C. elegans* AGOs.**

**Table S2. A table summarizing small RNA library information.**

**Table S3. A table summarizing the number and biotype of sRNAs enriched in AGO IPs.**

**Table S4. A table summarizing the mirDeep2 analysis results in predicting novel high confidence miRNAs.**

**Table S5. A table summarizing the 466 unannotated 21U sRNA sequences enriched in PRG-1 IPs and depleted in *prg-1* mutants.**

**Table S6. Tables summarizing the enrichment of reads in all libraries used in this study.**

**Table S7. Gene Ontology analysis results of AGO targets.**

**Table S8. Strains and primers used in this study.**

## References

Ahmed, S. and Hodgkin, J. (2000) ‘MRT-2 checkpoint protein is required for germline immortality and telomere replication in C. elegans.’, Nature, 403(6766), pp. 159–64. Available at: https://doi.org/10.1038/35003120.

Ahringer, J. (2006) ‘Reverse genetics’, WormBook [Preprint]. Available at: https://doi.org/10.1895/wormbook.1.47.1.

Ambros, V. et al. (2003) ‘MicroRNAs and other tiny endogenous RNAs in C. elegans.’, Curr Biol, 13(10), pp. 807–18.

Angeles-Albores, D. et al. (2016) ‘Tissue enrichment analysis for C. elegans genomics’, BMC Bioinformatics, 17(1), p. 366. Available at: https://doi.org/10.1186/s12859-016-1229-9.

Angeles-Albores, D. et al. (2018) ‘Two new functions in the WormBase Enrichment Suite’. Available at: https://doi.org/10.17912/W25Q2N.

Bakowski, M.A. et al. (2014) ‘Ubiquitin-Mediated Response to Microsporidia and Virus Infection in C. elegans’, PLoS Pathogens. Edited by D.S. Schneider, 10(6), p. e1004200. Available at: https://doi.org/10.1371/journal.ppat.1004200.

Batista, P.J. et al. (2008) ‘PRG-1 and 21U-RNAs interact to form the piRNA complex required for fertility in C. elegans.’, Mol Cell, 31(1), pp. 67–78. Available at: https://doi.org/10.1016/j.molcel.2008.06.002.

Bolte, S. and Cordelières, F.P. (2006) ‘A guided tour into subcellular colocalization analysis in light microscopy’, Journal of Microscopy, 224(3), pp. 213–232. Available at: https://doi.org/10.1111/j.1365-2818.2006.01706.x.

Braukmann, F., Jordan, D. and Miska, E. (2017) ‘Artificial and natural RNA interactions between bacteria and *C. elegans*’, RNA Biology, 14(4), pp. 415–420. Available at: https://doi.org/10.1080/15476286.2017.1297912.

Brenner, S. (1974) ‘The genetics of Caenorhabditis elegans.’, Genetics, 77(1), pp. 71–94.

Brown, K.C. et al. (2017) ‘ALG-5 is a miRNA-associated Argonaute required for proper developmental timing in the Caenorhabditis elegans germline.’, Nucleic Acids Res, 45(15), pp. 9093–9107. Available at: https://doi.org/10.1093/nar/gkx536.

Buckley, B.A. et al. (2012) ‘A nuclear Argonaute promotes multigenerational epigenetic inheritance and germline immortality.’, Nature, 489(7416), pp. 447–51. Available at: https://doi.org/10.1038/nature11352.

Bukhari, S.I.A. et al. (2012) ‘The microRNA pathway controls germ cell proliferation and differentiation in C. elegans’, Cell Research, 22(6), pp. 1034–1045. Available at: https://doi.org/10.1038/cr.2012.31.

Cecere, G. (2021) ‘Small RNAs in epigenetic inheritance: from mechanisms to trait transmission’, FEBS letters, 595(24), pp. 2953–2977. Available at: https://doi.org/10.1002/1873-3468.14210.

Charlesworth, A.G. et al. (2021) ‘Two isoforms of the essential *C. elegans* Argonaute CSR-1 differentially regulate sperm and oocyte fertility’, *Nucleic Acids Research*, p. gkab619. Available at: https://doi.org/10.1093/nar/gkab619.

Chaves, D.A. et al. (2021) ‘The RNA phosphatase PIR-1 regulates endogenous small RNA pathways in C. elegans’, Molecular Cell, 81(3), pp. 546–557.e5. Available at: https://doi.org/10.1016/j.molcel.2020.12.004.

Claycomb, J.M. et al. (2009) ‘The Argonaute CSR-1 and its 22G-RNA cofactors are required for holocentric chromosome segregation.’, Cell, 139(1), pp. 123–34. Available at: https://doi.org/10.1016/j.cell.2009.09.014.

Conine, C.C. et al. (2010) ‘Argonautes ALG-3 and ALG-4 are required for spermatogenesis-specific 26G-RNAs and thermotolerant sperm in Caenorhabditis elegans.’, Proc Natl Acad Sci U S A, 107(8), pp. 3588–93. Available at: https://doi.org/10.1073/pnas.0911685107.

Cornes, E. et al. (2022) ‘piRNAs initiate transcriptional silencing of spermatogenic genes during C. elegans germline development’, Developmental Cell, 57(2), pp. 180–196.e7. Available at: https://doi.org/10.1016/j.devcel.2021.11.025.

Correa, R.L. et al. (2010) ‘MicroRNA-directed siRNA biogenesis in Caenorhabditis elegans.’, PLoS Genet, 6(4), p. e1000903. Available at: https://doi.org/10.1371/journal.pgen.1000903.

Dickinson, D.J. et al. (2015) ‘Streamlined Genome Engineering with a Self-Excising Drug Selection Cassette.’, Genetics, 200(4), pp. 1035–49. Available at: https://doi.org/10.1534/genetics.115.178335.

Dueck, A. and Meister, G. (2014) ‘Assembly and function of small RNA – Argonaute protein complexes.’, Biol Chem [Preprint]. Available at: https://doi.org/10.1515/hsz-2014-0116.

Ewe, C.K. et al. (2020) ‘Natural cryptic variation in epigenetic modulation of an embryonic gene regulatory network’, Proceedings of the National Academy of Sciences, 117(24), pp. 13637–13646. Available at: https://doi.org/10.1073/pnas.1920343117.

Felix, M.A. et al. (2011) ‘Natural and experimental infection of Caenorhabditis nematodes by novel viruses related to nodaviruses.’, PLoS Biol, 9(1), p. e1000586. Available at: https://doi.org/10.1371/journal.pbio.1000586.

Frank, F., Sonenberg, N. and Nagar, B. (2010) ‘Structural basis for 5’-nucleotide base-specific recognition of guide RNA by human AGO2.’, Nature, 465(7299), pp. 818–22. Available at: https://doi.org/10.1038/nature09039.

Franz, C.J. et al. (2014) ‘Orsay, Santeuil and Le Blanc viruses primarily infect intestinal cells in Caenorhabditis nematodes’, Virology, 448, pp. 255–264. Available at: https://doi.org/10.1016/j.virol.2013.09.024.

Friedländer, M.R. et al. (2012) ‘miRDeep2 accurately identifies known and hundreds of novel microRNA genes in seven animal clades’, Nucleic Acids Research, 40(1), pp. 37–52. Available at: https://doi.org/10.1093/nar/gkr688.

Grishok, A. et al. (2001) ‘Genes and mechanisms related to RNA interference regulate expression of the small temporal RNAs that control C. elegans developmental timing.’, Cell, 106(1), pp. 23–34.

Gu, W. et al. (2009) ‘Distinct argonaute-mediated 22G-RNA pathways direct genome surveillance in the C. elegans germline.’, Mol Cell, 36(2), pp. 231–44. Available at: https://doi.org/10.1016/j.molcel.2009.09.020.

Guang, S. et al. (2008) ‘An Argonaute transports siRNAs from the cytoplasm to the nucleus.’, Science, 321(5888), pp. 537–41. Available at: https://doi.org/10.1126/science.1157647.

Han, T. et al. (2009) ‘26G endo-siRNAs regulate spermatogenic and zygotic gene expression in Caenorhabditis elegans.’, Proc Natl Acad Sci U S A, 106(44), pp. 18674–9. Available at: https://doi.org/10.1073/pnas.0906378106.

Houri-Zeevi, L. et al. (2021) ‘Stress resets ancestral heritable small RNA responses’, eLife, 10, p. e65797. Available at: https://doi.org/10.7554/eLife.65797.

Kaletsky, R. et al. (2020) ‘C. elegans interprets bacterial non-coding RNAs to learn pathogenic avoidance’, Nature, 586(7829), pp. 445–451. Available at: https://doi.org/10.1038/s41586-020-2699-5.

Kim, D.H. (2002) ‘A Conserved p38 MAP Kinase Pathway in Caenorhabditis elegans Innate Immunity’, Science, 297(5581), pp. 623–626. Available at: https://doi.org/10.1126/science.1073759.

Kumar, S. et al. (2018) ‘MEGA X: Molecular Evolutionary Genetics Analysis across Computing Platforms’, Molecular Biology and Evolution. Edited by F.U. Battistuzzi, 35(6), pp. 1547–1549. Available at: https://doi.org/10.1093/molbev/msy096.

Lee, H.C. et al. (2012) ‘C. elegans piRNAs Mediate the Genome-wide Surveillance of Germline Transcripts.’, Cell, 150(1), pp. 78–87. Available at: https://doi.org/10.1016/j.cell.2012.06.016.

Love, M.I., Huber, W. and Anders, S. (2014) ‘Moderated estimation of fold change and dispersion for RNA-seq data with DESeq2.’, Genome Biol, 15(12), p. 550. Available at: https://doi.org/10.1186/s13059-014-0550-8.

Ma, J.B. et al. (2005) ‘Structural basis for 5’-end-specific recognition of guide RNA by the A. fulgidus Piwi protein.’, Nature, 434(7033), pp. 666–70. Available at: https://doi.org/10.1038/nature03514.

Madeira, F. et al. (2019) ‘The EMBL-EBI search and sequence analysis tools APIs in 2019’, Nucleic Acids Research, 47(W1), pp. W636–W641. Available at: https://doi.org/10.1093/nar/gkz268.

Meister, G. (2013) ‘Argonaute proteins: functional insights and emerging roles.’, Nat Rev Genet, 14(7), pp. 447–59. Available at: https://doi.org/10.1038/nrg3462.

modENCODE Consortium et al. (2009) ‘Unlocking the secrets of the genome’, Nature, 459(7249), pp. 927–930. Available at: https://doi.org/10.1038/459927a.

Montgomery, B.E. et al. (2021) ‘Dual roles for piRNAs in promoting and preventing gene silencing in C. elegans’, Cell Reports, 37(10), p. 110101. Available at: https://doi.org/10.1016/j.celrep.2021.110101.

Moore, R.S., Kaletsky, R. and Murphy, C.T. (2019) ‘Piwi/PRG-1 Argonaute and TGF-β Mediate Transgenerational Learned Pathogenic Avoidance’, Cell, 177(7), pp. 1827–1841.e12. Available at: https://doi.org/10.1016/j.cell.2019.05.024.

Nakanishi, K. et al. (2012) ‘Structure of yeast Argonaute with guide RNA.’, Nature, 486(7403), pp. 368–74. Available at: https://doi.org/10.1038/nature11211.

Nam, J.-W. and Bartel, D.P. (2012) ‘Long noncoding RNAs in *C. elegans*’, Genome Research, 22(12), pp. 2529–2540. Available at: https://doi.org/10.1101/gr.140475.112.

Nguyen, D.A.H. and Phillips, C.M. (2021) ‘Arginine methylation promotes siRNA-binding specificity for a spermatogenesis-specific isoform of the Argonaute protein CSR-1’, Nature Communications, 12(1), p. 4212. Available at: https://doi.org/10.1038/s41467-021-24526-6.

Ni, J.Z. et al. (2016) ‘A transgenerational role of the germline nuclear RNAi pathway in repressing heat stress-induced transcriptional activation in C. elegans.’, Epigenetics Chromatin, 9, p. 3. Available at: https://doi.org/10.1186/s13072-016-0052-x.

Ortiz, M.A. et al. (2014) ‘A new dataset of spermatogenic vs. oogenic transcriptomes in the nematode Caenorhabditis elegans.’, G3 (Bethesda), 4(9), pp. 1765–72. Available at: https://doi.org/10.1534/g3.114.012351.

Ozata, D.M. et al. (2019) ‘PIWI-interacting RNAs: small RNAs with big functions.’, Nat Rev Genet, 20(2), pp. 89–108. Available at: https://doi.org/10.1038/s41576-018-0073-3.

Phillips, C.M. et al. (2012) ‘MUT-16 promotes formation of perinuclear mutator foci required for RNA silencing in the C. elegans germline.’, Genes Dev, 26(13), pp. 1433–44. Available at: https://doi.org/10.1101/gad.193904.112.

Pink, R.C. et al. (2011) ‘Pseudogenes: Pseudo-functional or key regulators in health and disease?’, RNA, 17(5), pp. 792–798. Available at: https://doi.org/10.1261/rna.2658311.

Rechavi, O. et al. (2014) ‘Starvation-induced transgenerational inheritance of small RNAs in C. elegans.’, Cell, 158(2), pp. 277–87. Available at: https://doi.org/10.1016/j.cell.2014.06.020.

Rechavi, O., Minevich, G. and Hobert, O. (2011) ‘Transgenerational Inheritance of an Acquired Small RNA-Based Antiviral Response in C. elegans.’, Cell, 147(6), pp. 1248–56. Available at: https://doi.org/10.1016/j.cell.2011.10.042.

Reddy, K.C. et al. (2019) ‘Antagonistic paralogs control a switch between growth and pathogen resistance in C. elegans’, PLOS Pathogens. Edited by J.J. Collins, 15(1), p. e1007528. Available at: https://doi.org/10.1371/journal.ppat.1007528.

Ruby, J.G. et al. (2006) ‘Large-scale sequencing reveals 21U-RNAs and additional microRNAs and endogenous siRNAs in C. elegans.’, Cell, 127(6), pp. 1193–207. Available at: https://doi.org/10.1016/j.cell.2006.10.040.

Sapetschnig, A. et al. (2015) ‘Tertiary siRNAs Mediate Paramutation in C. elegans’, PLOS Genetics. Edited by N.C. Lau, 11(3), p. e1005078. Available at: https://doi.org/10.1371/journal.pgen.1005078.

Schneider, C.A., Rasband, W.S. and Eliceiri, K.W. (2012) ‘NIH Image to ImageJ: 25 years of image analysis.’, Nat Methods, 9(7), pp. 671–5.

Schott, D., Yanai, I. and Hunter, C.P. (2014) ‘Natural RNA interference directs a heritable response to the environment.’, Sci Rep, 4, p. 7387. Available at: https://doi.org/10.1038/srep07387.

Schreier, J. et al. (2022) ‘Membrane-associated cytoplasmic granules carrying the Argonaute protein WAGO-3 enable paternal epigenetic inheritance in Caenorhabditis elegans’, Nature Cell Biology, 24(2), pp. 217–229. Available at: https://doi.org/10.1038/s41556-021-00827-2.

Sheu-Gruttadauria, J. and MacRae, I.J. (2017) ‘Structural Foundations of RNA Silencing by Argonaute’, Journal of Molecular Biology, 429(17), pp. 2619–2639. Available at: https://doi.org/10.1016/j.jmb.2017.07.018.

Shirayama, M. et al. (2012) ‘piRNAs Initiate an Epigenetic Memory of Nonself RNA in the C. elegans Germline.’, Cell, 150(1), pp. 65–77. Available at: https://doi.org/10.1016/j.cell.2012.06.015.

Simon, M. et al. (2014) ‘Reduced insulin/IGF-1 signaling restores germ cell immortality to Caenorhabditis elegans Piwi mutants.’, Cell Rep, 7(3), pp. 762–73. Available at: https://doi.org/10.1016/j.celrep.2014.03.056.

Spracklin, G. et al. (2017) ‘The RNAi Inheritance Machinery of *Caenorhabditis elegans*’, Genetics, 206(3), pp. 1403–1416. Available at: https://doi.org/10.1534/genetics.116.198812.

Statello, L. et al. (2021) ‘Gene regulation by long non-coding RNAs and its biological functions’, Nature Reviews Molecular Cell Biology, 22(2), pp. 96–118. Available at: https://doi.org/10.1038/s41580-020-00315-9.

Steiner, F.A. et al. (2007) ‘Structural features of small RNA precursors determine Argonaute loading in Caenorhabditis elegans.’, Nat Struct Mol Biol, 14(10), pp. 927–33. Available at: https://doi.org/10.1038/nsmb1308.

Sundby, A.E., Molnar, R.I. and Claycomb, J.M. (2021) ‘Connecting the Dots: Linking Caenorhabditis elegans Small RNA Pathways and Germ Granules’, Trends in Cell Biology, p. S0962892421000040. Available at: https://doi.org/10.1016/j.tcb.2020.12.012.

Svendsen, J.M. et al. (2019) ‘henn-1/HEN1 Promotes Germline Immortality in Caenorhabditis elegans’, Cell Reports, 29(10), pp. 3187–3199.e4. Available at: https://doi.org/10.1016/j.celrep.2019.10.114.

Swarts, D.C. et al. (2014) ‘The evolutionary journey of Argonaute proteins.’

Tabara, H. et al. (1999) ‘The rde-1 gene, RNA interference, and transposon silencing in C. elegans.’, Cell, 99(2), pp. 123–32.

Tzur, Y.B. et al. (2018) ‘Spatiotemporal Gene Expression Analysis of the *Caenorhabditis elegans* Germline Uncovers a Syncytial Expression Switch’, Genetics, 210(2), pp. 587–605. Available at: https://doi.org/10.1534/genetics.118.301315.

Vasale, J.J. et al. (2010) ‘Sequential rounds of RNA-dependent RNA transcription drive endogenous small-RNA biogenesis in the ERGO-1/Argonaute pathway.’, Proc Natl Acad Sci U S A, 107(8), pp. 3582–7. Available at: https://doi.org/10.1073/pnas.0911908107.

Wan, G. et al. (2018) ‘Spatiotemporal regulation of liquid-like condensates in epigenetic inheritance.’, Nature, 557(7707), pp. 679–683. Available at: https://doi.org/10.1038/s41586-018-0132-0.

Wang, G. and Reinke, V. (2008) ‘A C. elegans Piwi, PRG-1, regulates 21U-RNAs during spermatogenesis.’, Curr Biol, 18(12), pp. 861–7. Available at: https://doi.org/10.1016/j.cub.2008.05.009.

Wedeles, C.J., Wu, M.Z. and Claycomb, J.M. (2013) ‘Protection of germline gene expression by the C. elegans Argonaute CSR-1.’, Dev Cell, 27(6), pp. 664–71. Available at: https://doi.org/10.1016/j.devcel.2013.11.016.

Wu, J. et al. (2020) ‘Argonaute proteins: Structural features, functions and emerging roles’, Journal of Advanced Research, 24, pp. 317–324. Available at: https://doi.org/10.1016/j.jare.2020.04.017.

Xu, F. et al. (2018) ‘A Cytoplasmic Argonaute Protein Promotes the Inheritance of RNAi.’, Cell Rep, 23(8), pp. 2482–2494. Available at: https://doi.org/10.1016/j.celrep.2018.04.072.

Yigit, E. et al. (2006) ‘Analysis of the C. elegans Argonaute family reveals that distinct Argonautes act sequentially during RNAi.’, Cell, 127(4), pp. 747–57. Available at: https://doi.org/10.1016/j.cell.2006.09.033.

Zhang, L. et al. (2015) ‘The auxin-inducible degradation (AID) system enables versatile conditional protein depletion in C. elegans.’, Development, 142(24), pp. 4374–84. Available at: https://doi.org/10.1242/dev.129635.

