## Supplemental Figures for "A Comprehensive Survey of *C. elegans* Argonaute Proteins Reveals Organism-wide Gene Regulatory Networks and Functions"

### Figure S1

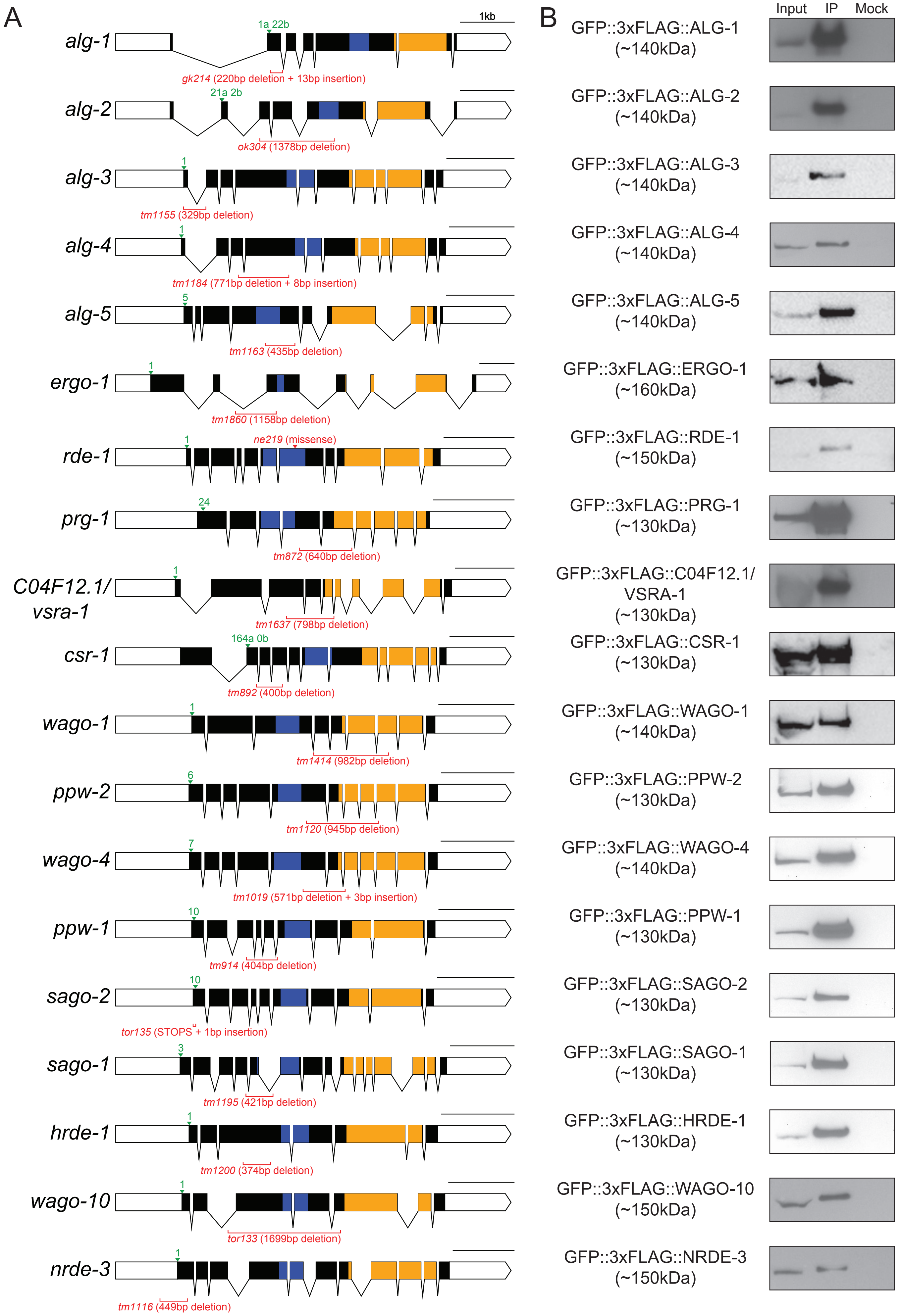
A
B
1kb
alg-1
1a 22b
gk214 (220bp deletion + 13bp insertion)
alg-2
21a 2b
ok304 (1378bp deletion)
alg-3
1
tm1155 (329bp deletion)
alg-4
1
tm1184 (771bp deletion + 8bp insertion)
alg-5
5
tm1163 (435bp deletion)
ergo-1
1
tm1860 (1158bp deletion)
rde-1
1
ne219 (missense)
prg-1
24
tm872 (640bp deletion)
C04F12.1/ vsra-1
1
tm1637 (798bp deletion)
csr-1
164a 0b
tm892 (400bp deletion)
wago-1
1
tm1414 (982bp deletion)
ppw-2
6
tm1120 (945bp deletion)
wago-4
7
tm1019 (571bp deletion + 3bp insertion)
ppw-1
10
tm914 (404bp deletion)
sago-2
10
tor135 (STOPS + 1bp insertion)
sago-1
3
tm1195 (421bp deletion)
hrde-1
1
tm1200 (374bp deletion)
wago-10
1
tor133 (1699bp deletion)
nrde-3
1
tm1116 (449bp deletion)
Input
IP
Mock
GFP::3xFLAG::ALG-1 (~140kDa)
GFP::3xFLAG::ALG-2 (~140kDa)
GFP::3xFLAG::ALG-3 (~140kDa)
GFP::3xFLAG::ALG-4 (~140kDa)
GFP::3xFLAG::ALG-5 (~140kDa)
GFP::3xFLAG::ERGO-1 (~160kDa)
GFP::3xFLAG::RDE-1 (~150kDa)
GFP::3xFLAG::PRG-1 (~130kDa)
GFP::3xFLAG::C04F12.1/ VSRA-1 (~130kDa)
GFP::3xFLAG::CSR-1 (~130kDa)
GFP::3xFLAG::WAGO-1 (~140kDa)
GFP::3xFLAG::PPW-2 (~130kDa)
GFP::3xFLAG::WAGO-4 (~140kDa)
GFP::3xFLAG::PPW-1 (~130kDa)
GFP::3xFLAG::SAGO-2 (~130kDa)
GFP::3xFLAG::SAGO-1 (~130kDa)
GFP::3xFLAG::HRDE-1 (~130kDa)
GFP::3xFLAG::WAGO-10 (~150kDa)
GFP::3xFLAG::NRDE-3 (~150kDa)

Figure S2

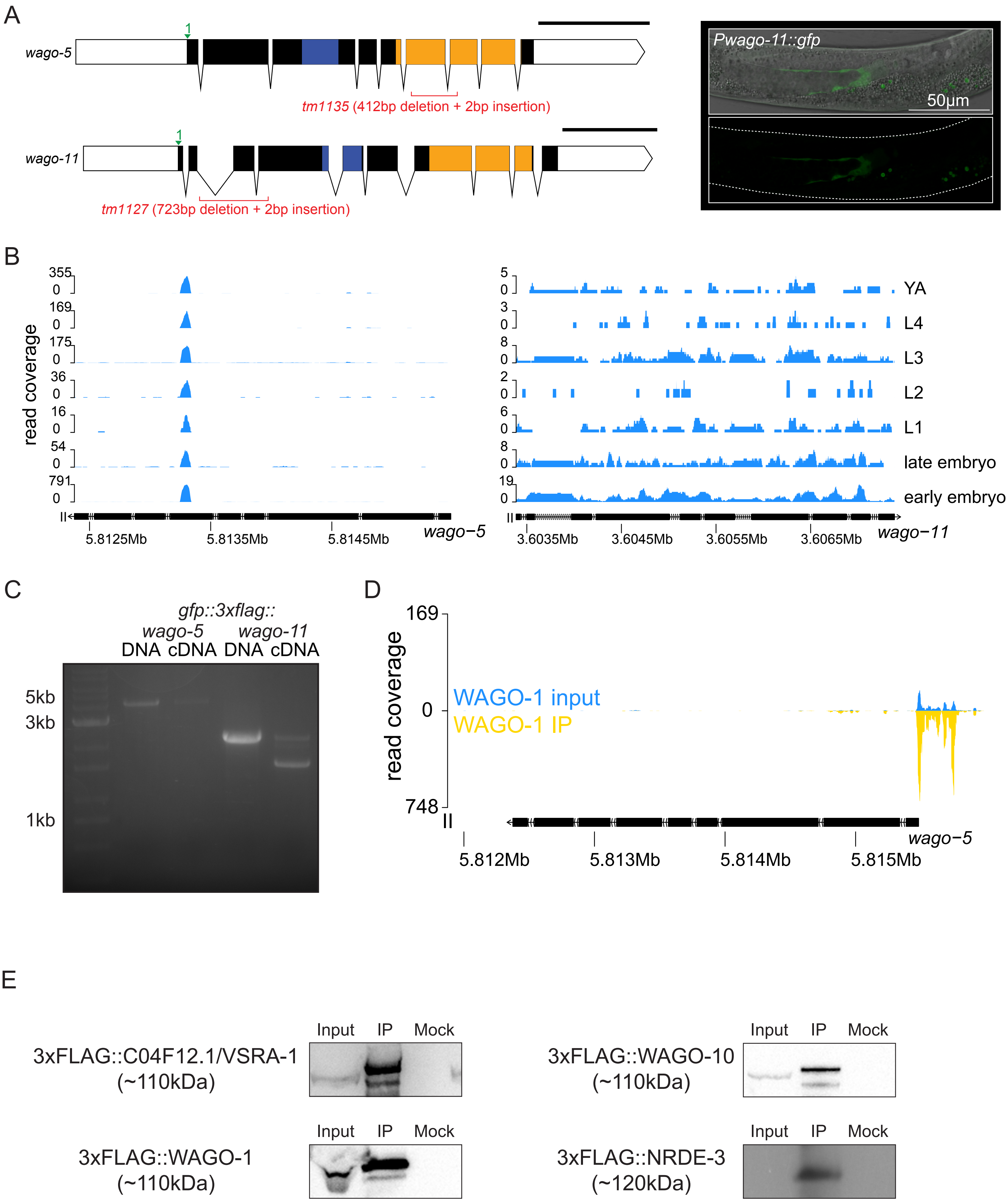
A
wago-5
1
tm1135 (412bp deletion + 2bp insertion)
wago-11
1
tm1127 (723bp deletion + 2bp insertion)
Pwago-11::gfp
50μm
B
read coverage
YA
L4
L3
L2
L1
late embryo
early embryo
5.8125Mb
5.8135Mb
5.8145Mb
wago−5
3.6035Mb
3.6045Mb
3.6055Mb
3.6065Mb
wago−11
C
gfp::3xflag::
wago-5
wago-11
DNA cDNA DNA cDNA
5kb
3kb
1kb
D
169
0
748
read coverage
WAGO-1 input
WAGO-1 IP
5.812Mb
5.813Mb
5.814Mb
5.815Mb
wago−5
E
Input IP Mock
3xFLAG::C04F12.1/VSRA-1
(~110kDa)
3xFLAG::WAGO-10
(~110kDa)
3xFLAG::WAGO-1
(~110kDa)
3xFLAG::NRDE-3
(~120kDa)

Figure S3

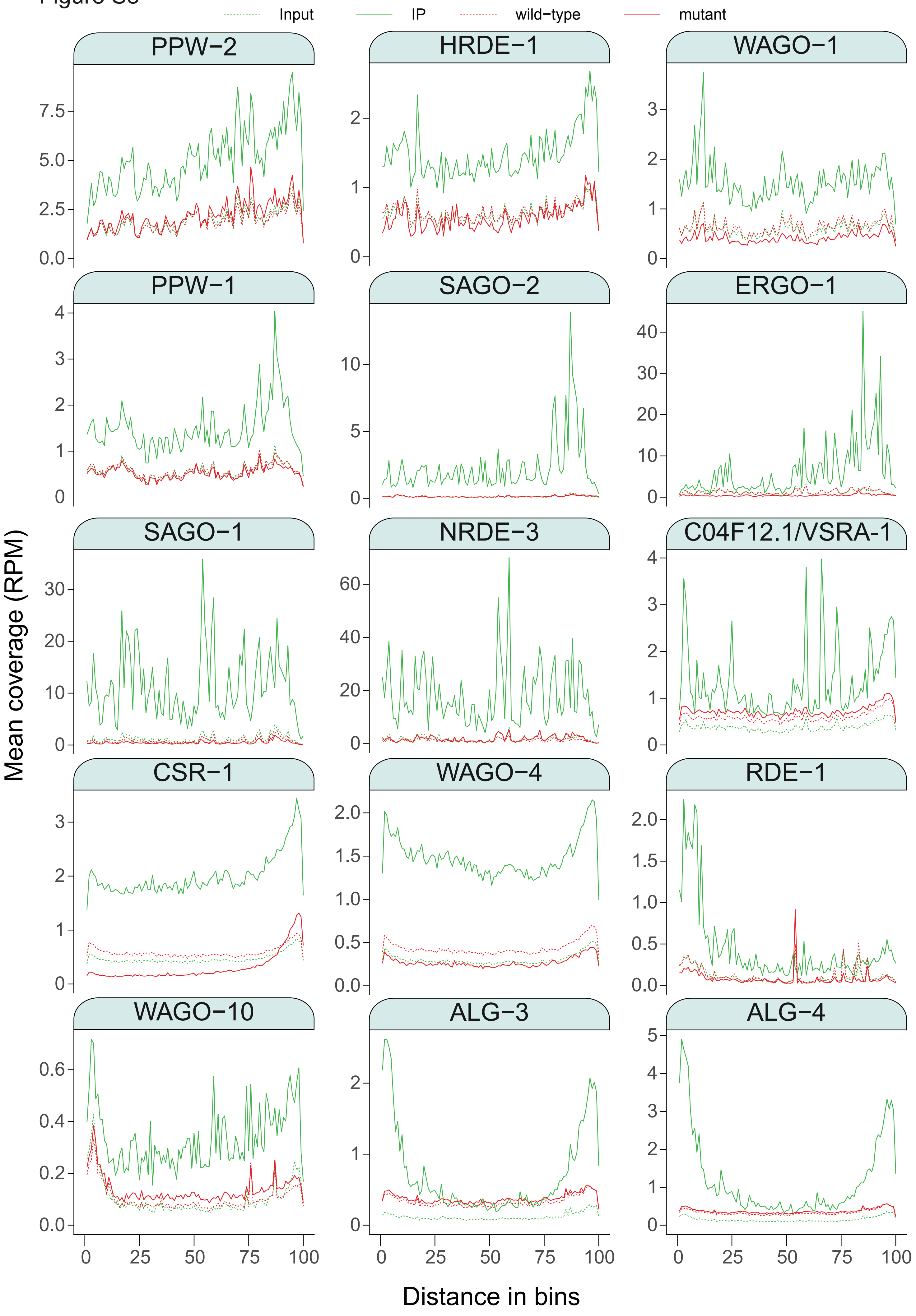

Input
IP
wild−type
mutant
PPW−2
HRDE−1
WAGO−1
PPW−1
SAGO−2
ERGO−1
SAGO−1
NRDE−3
C04F12.1/VSRA−1
CSR−1
WAGO−4
RDE−1
WAGO−10
ALG−3
ALG−4
Mean coverage (RPM)
Distance in bins

Figure S4

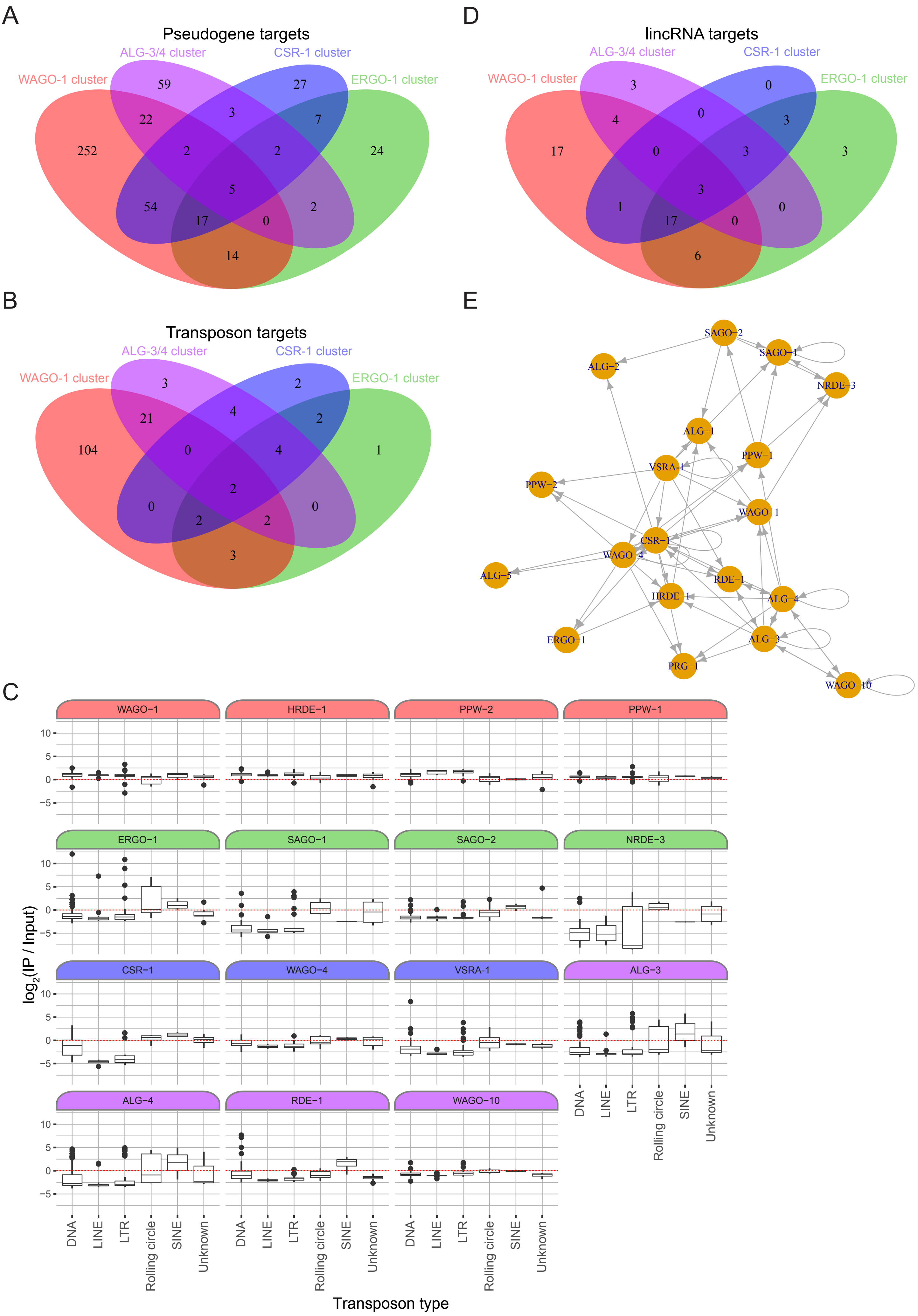
A
Pseudogene targets
WAGO-1 cluster
ALG-3/4 cluster
CSR-1 cluster
ERGO-1 cluster
D
lincRNA targets
WAGO-1 cluster
ALG-3/4 cluster
CSR-1 cluster
ERGO-1 cluster
B
Transposon targets
WAGO-1 cluster
ALG-3/4 cluster
CSR-1 cluster
ERGO-1 cluster
E
SAGO-2
SAGO-1
ALG-2
NRDE-3
ALG-1
PPW-1
VSRA-1
PPW-2
WAGO-1
CSR-1
WAGO-4
ALG-5
RDE-1
HRDE-1
ALG-4
ERGO-1
ALG-3
PRG-1
WAGO-10
C
WAGO-1
HRDE-1
PPW-2
PPW-1
ERGO-1
SAGO-1
SAGO-2
NRDE-3
CSR-1
WAGO-4
VSRA-1
ALG-3
ALG-4
RDE-1
WAGO-10
log2(IP / Input)
DNA
LINE
LTR
Rolling circle
SINE
Unknown
Transposon type

### Figure S5

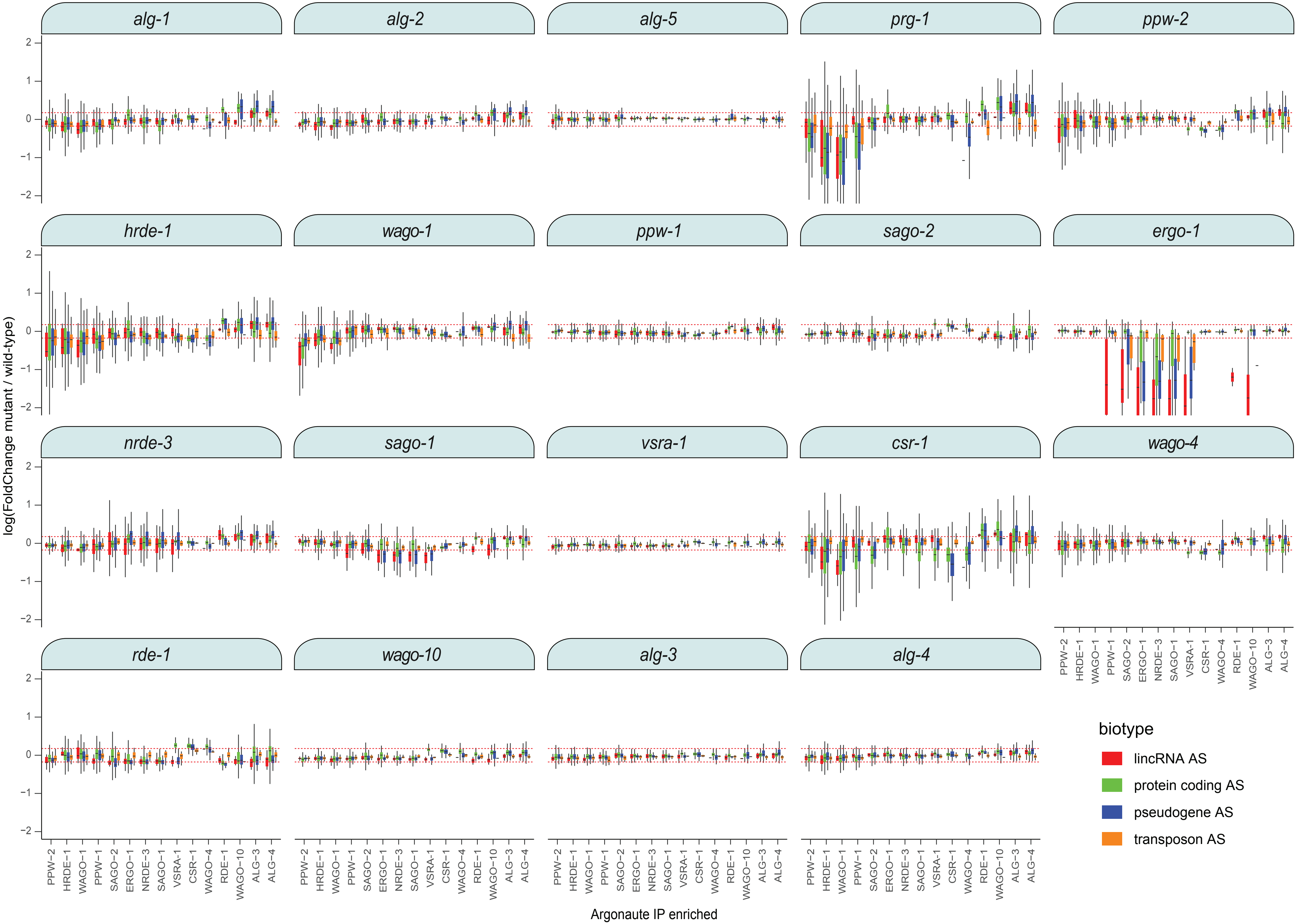
alg-1
alg-2
alg-5
prg-1
ppw-2
hrde-1
wago-1
ppw-1
sago-2
ergo-1
nrde-3
sago-1
vsra-1
csr-1
wago-4
rde-1
wago-10
alg-3
alg-4
log(FoldChange mutant / wild-type)
2
1
0
−1
−2
PPW−2
HRDE−1
WAGO−1
PPW−1
SAGO−2
ERGO−1
NRDE−3
SAGO−1
VSRA−1
CSR−1
WAGO−4
RDE−1
WAGO−10
ALG−3
ALG−4
Argonaute IP enriched
biotype
lincRNA AS
protein coding AS
pseudogene AS
transposon AS

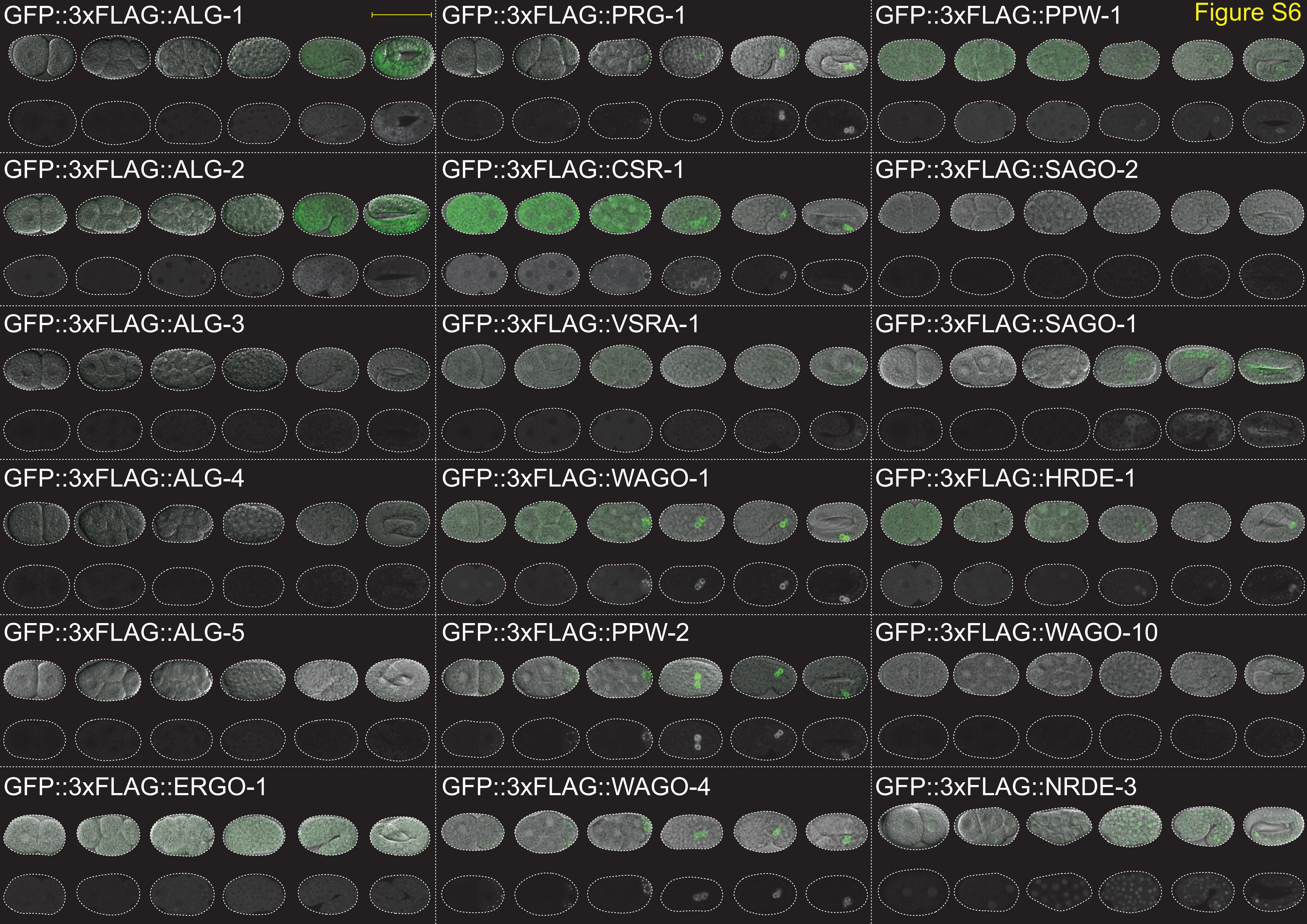
Figure S6
GFP::3xFLAG::ALG-1
GFP::3xFLAG::PRG-1
GFP::3xFLAG::PPW-1
GFP::3xFLAG::ALG-2
GFP::3xFLAG::CSR-1
GFP::3xFLAG::SAGO-2
GFP::3xFLAG::ALG-3
GFP::3xFLAG::VSRA-1
GFP::3xFLAG::SAGO-1
GFP::3xFLAG::ALG-4
GFP::3xFLAG::WAGO-1
GFP::3xFLAG::HRDE-1
GFP::3xFLAG::ALG-5
GFP::3xFLAG::PPW-2
GFP::3xFLAG::WAGO-10
GFP::3xFLAG::ERGO-1
GFP::3xFLAG::WAGO-4
GFP::3xFLAG::NRDE-3

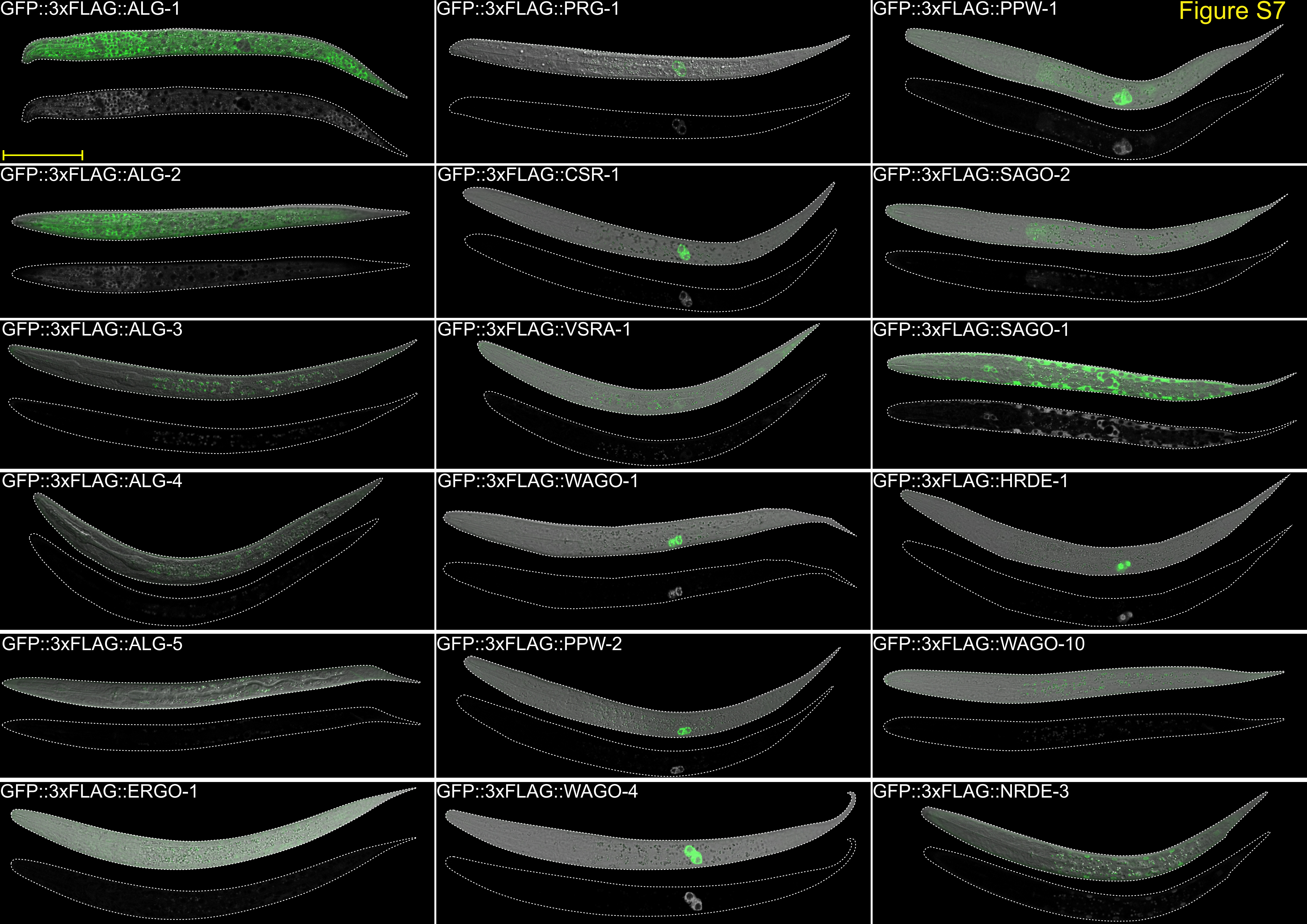
Figure S7
GFP::3xFLAG::ALG-1
GFP::3xFLAG::PRG-1
GFP::3xFLAG::PPW-1
GFP::3xFLAG::ALG-2
GFP::3xFLAG::CSR-1
GFP::3xFLAG::SAGO-2
GFP::3xFLAG::ALG-3
GFP::3xFLAG::VSRA-1
GFP::3xFLAG::SAGO-1
GFP::3xFLAG::ALG-4
GFP::3xFLAG::WAGO-1
GFP::3xFLAG::HRDE-1
GFP::3xFLAG::ALG-5
GFP::3xFLAG::PPW-2
GFP::3xFLAG::WAGO-10
GFP::3xFLAG::ERGO-1
GFP::3xFLAG::WAGO-4
GFP::3xFLAG::NRDE-3

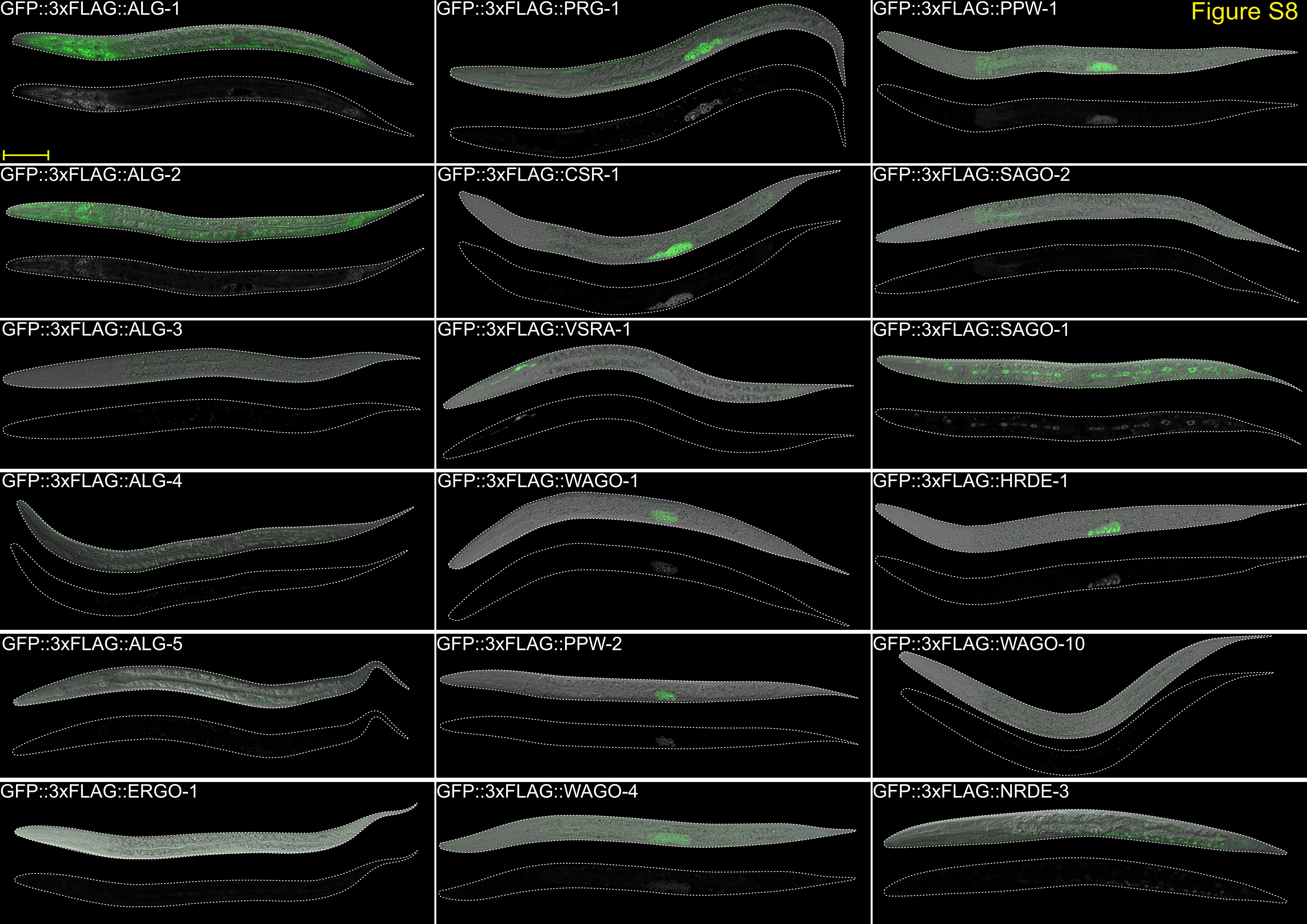
Figure S8
GFP::3xFLAG::ALG-1
GFP::3xFLAG::PRG-1
GFP::3xFLAG::PPW-1
GFP::3xFLAG::ALG-2
GFP::3xFLAG::CSR-1
GFP::3xFLAG::SAGO-2
GFP::3xFLAG::ALG-3
GFP::3xFLAG::VSRA-1
GFP::3xFLAG::SAGO-1
GFP::3xFLAG::ALG-4
GFP::3xFLAG::WAGO-1
GFP::3xFLAG::HRDE-1
GFP::3xFLAG::ALG-5
GFP::3xFLAG::PPW-2
GFP::3xFLAG::WAGO-10
GFP::3xFLAG::ERGO-1
GFP::3xFLAG::WAGO-4
GFP::3xFLAG::NRDE-3

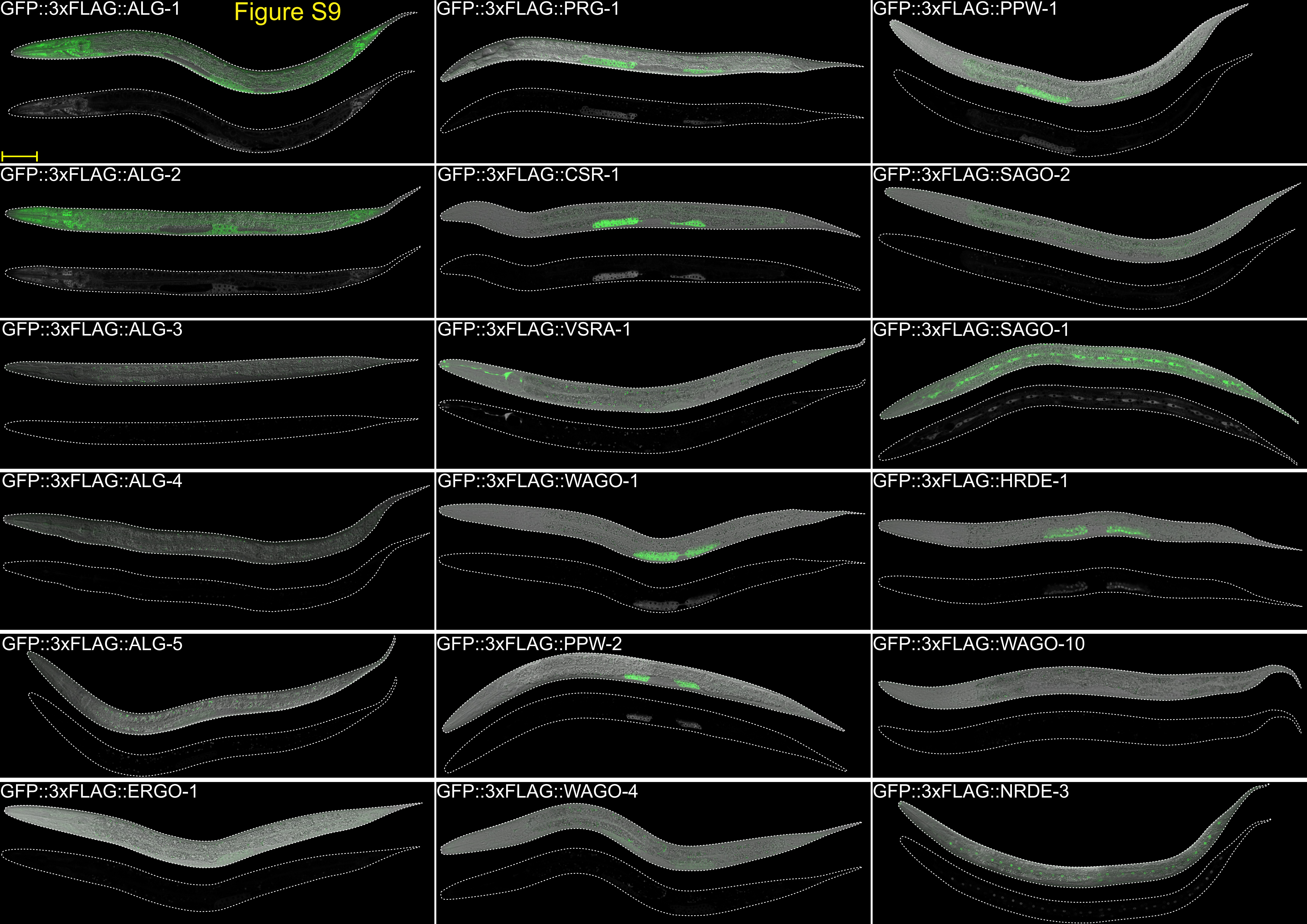
Figure S9
GFP::3xFLAG::ALG-1
GFP::3xFLAG::PRG-1
GFP::3xFLAG::PPW-1
GFP::3xFLAG::ALG-2
GFP::3xFLAG::CSR-1
GFP::3xFLAG::SAGO-2
GFP::3xFLAG::ALG-3
GFP::3xFLAG::VSRA-1
GFP::3xFLAG::SAGO-1
GFP::3xFLAG::ALG-4
GFP::3xFLAG::WAGO-1
GFP::3xFLAG::HRDE-1
GFP::3xFLAG::ALG-5
GFP::3xFLAG::PPW-2
GFP::3xFLAG::WAGO-10
GFP::3xFLAG::ERGO-1
GFP::3xFLAG::WAGO-4
GFP::3xFLAG::NRDE-3

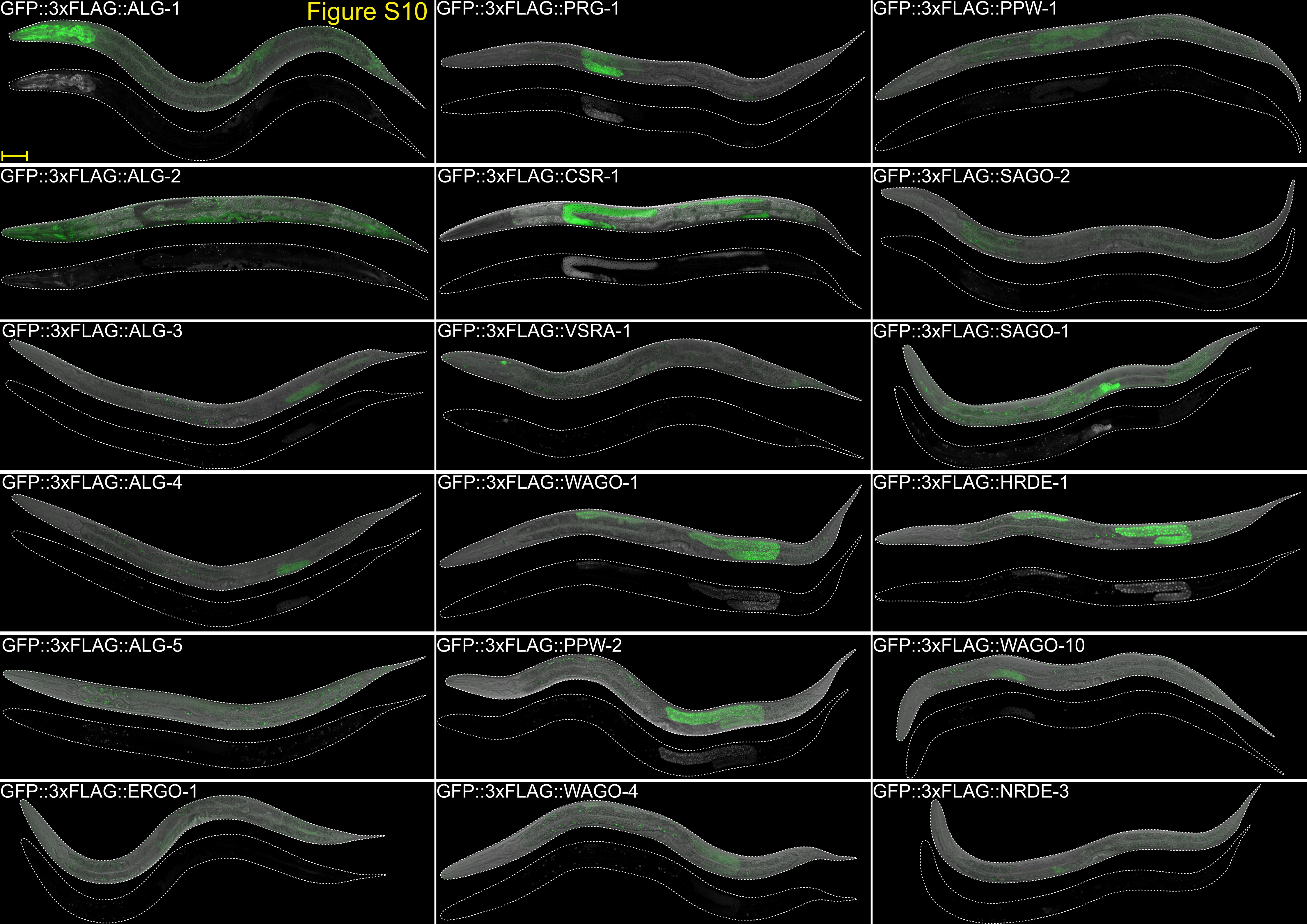
Figure S10
GFP::3xFLAG::ALG-1
GFP::3xFLAG::PRG-1
GFP::3xFLAG::PPW-1
GFP::3xFLAG::ALG-2
GFP::3xFLAG::CSR-1
GFP::3xFLAG::SAGO-2
GFP::3xFLAG::ALG-3
GFP::3xFLAG::VSRA-1
GFP::3xFLAG::SAGO-1
GFP::3xFLAG::ALG-4
GFP::3xFLAG::WAGO-1
GFP::3xFLAG::HRDE-1
GFP::3xFLAG::ALG-5
GFP::3xFLAG::PPW-2
GFP::3xFLAG::WAGO-10
GFP::3xFLAG::ERGO-1
GFP::3xFLAG::WAGO-4
GFP::3xFLAG::NRDE-3

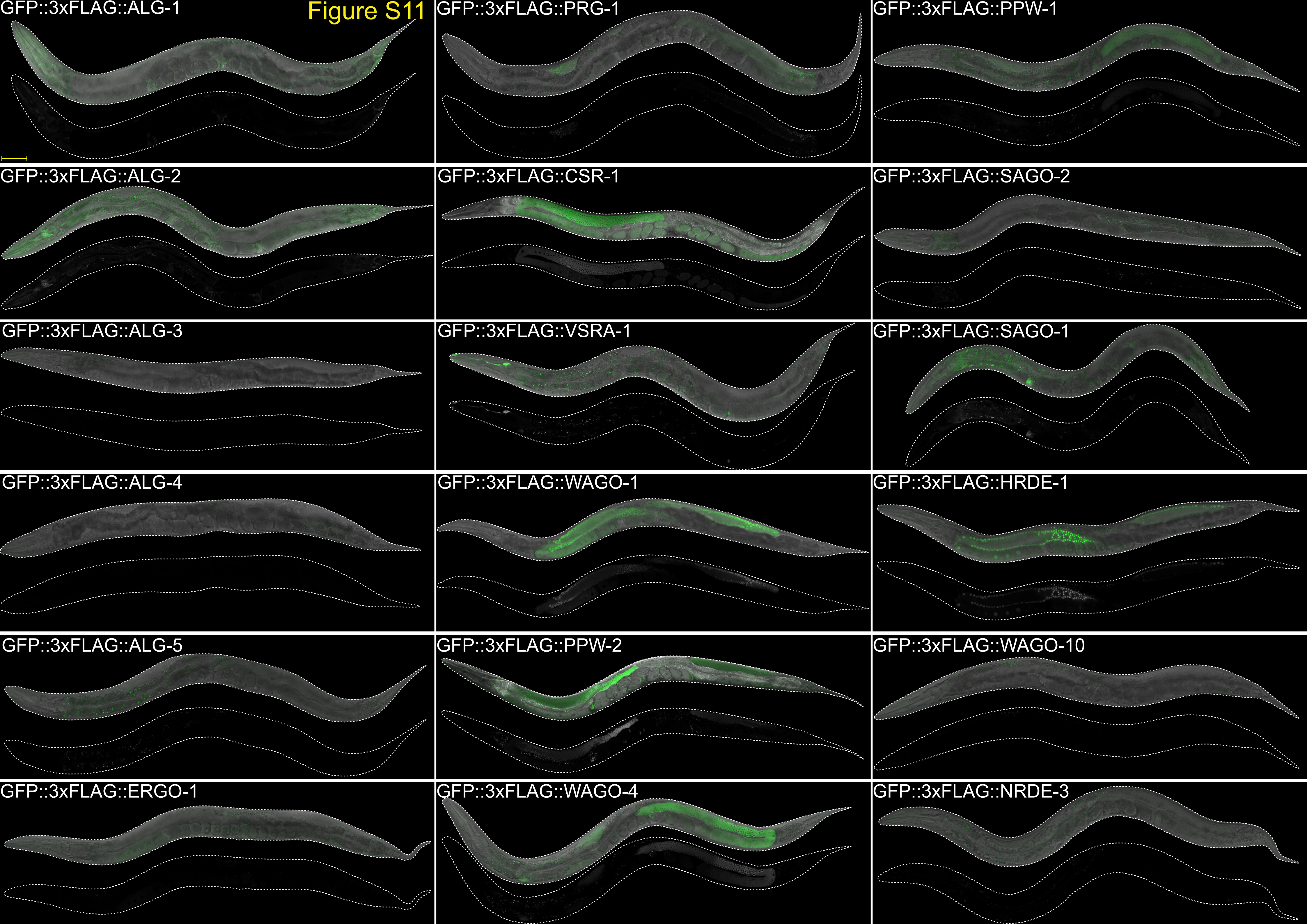
Figure S11
GFP::3xFLAG::ALG-1
GFP::3xFLAG::PRG-1
GFP::3xFLAG::PPW-1
GFP::3xFLAG::ALG-2
GFP::3xFLAG::CSR-1
GFP::3xFLAG::SAGO-2
GFP::3xFLAG::ALG-3
GFP::3xFLAG::VSRA-1
GFP::3xFLAG::SAGO-1
GFP::3xFLAG::ALG-4
GFP::3xFLAG::WAGO-1
GFP::3xFLAG::HRDE-1
GFP::3xFLAG::ALG-5
GFP::3xFLAG::PPW-2
GFP::3xFLAG::WAGO-10
GFP::3xFLAG::ERGO-1
GFP::3xFLAG::WAGO-4
GFP::3xFLAG::NRDE-3

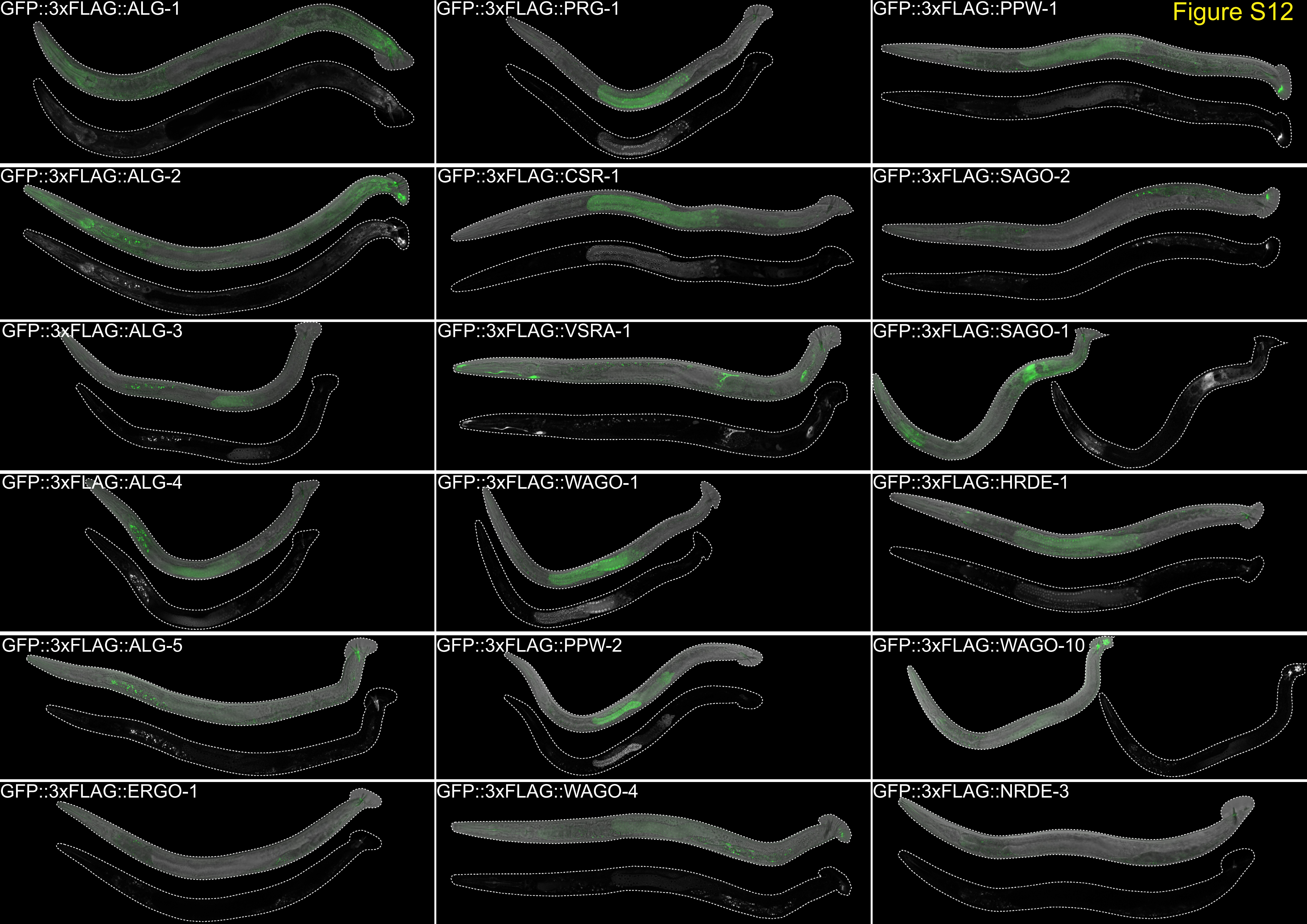
Figure S12
GFP::3xFLAG::ALG-1
GFP::3xFLAG::PRG-1
GFP::3xFLAG::PPW-1
GFP::3xFLAG::ALG-2
GFP::3xFLAG::CSR-1
GFP::3xFLAG::SAGO-2
GFP::3xFLAG::ALG-3
GFP::3xFLAG::VSRA-1
GFP::3xFLAG::SAGO-1
GFP::3xFLAG::ALG-4
GFP::3xFLAG::WAGO-1
GFP::3xFLAG::HRDE-1
GFP::3xFLAG::ALG-5
GFP::3xFLAG::PPW-2
GFP::3xFLAG::WAGO-10
GFP::3xFLAG::ERGO-1
GFP::3xFLAG::WAGO-4
GFP::3xFLAG::NRDE-3

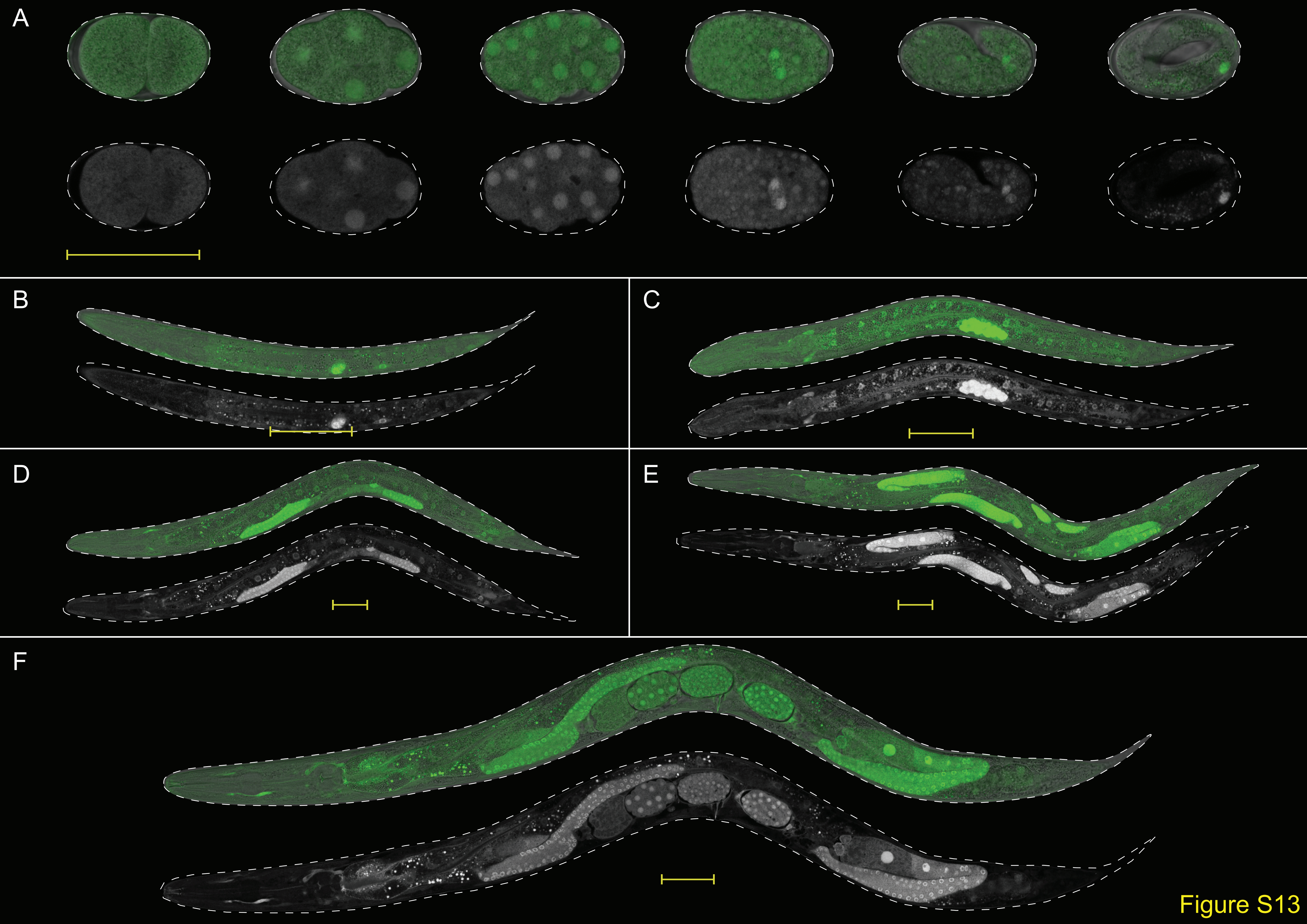
A
B
C
D
E
F
Figure S13

Figure S14

A

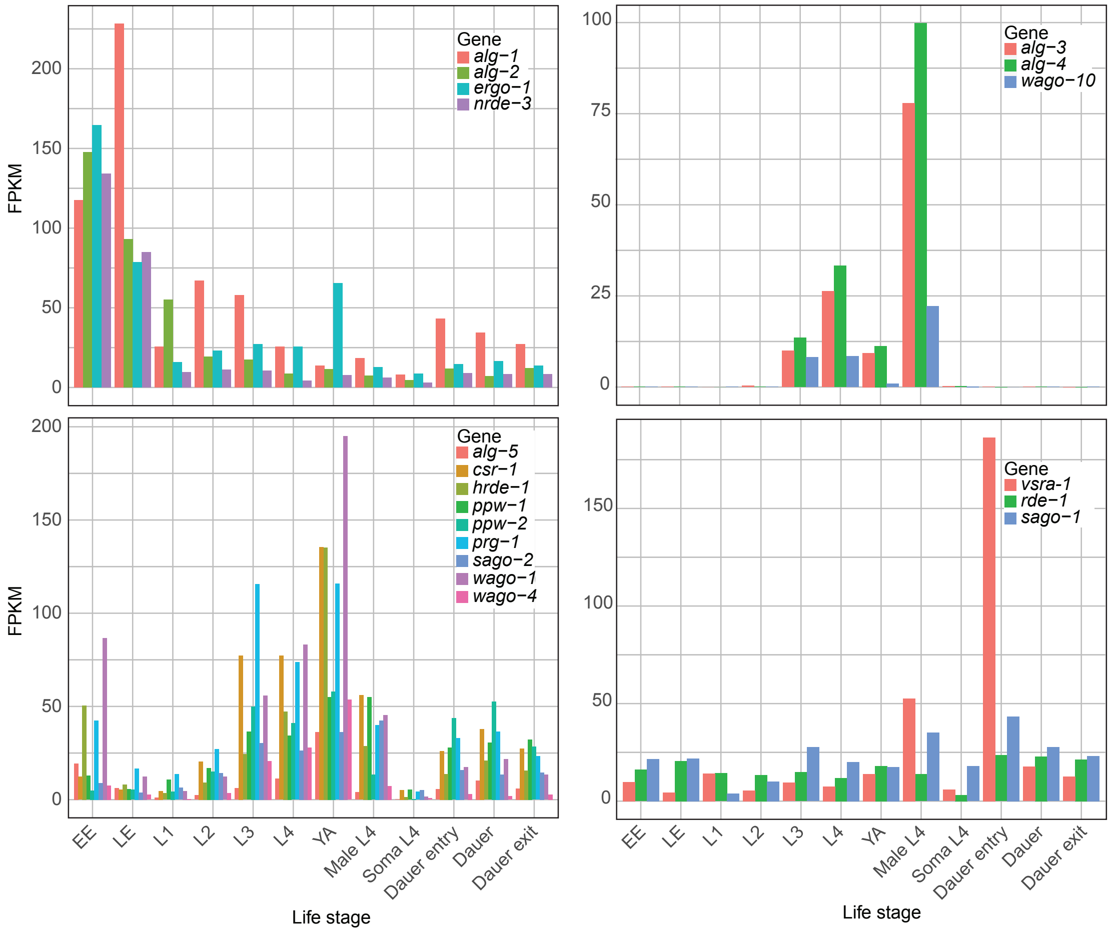


B

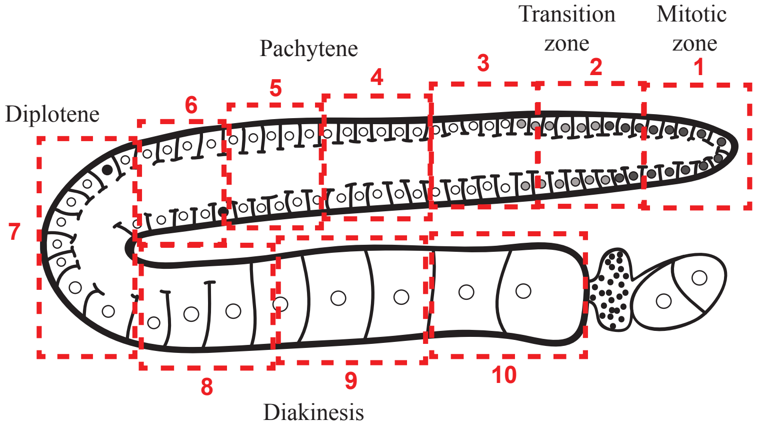


C

Relative expression by row: −2 −1 0 1 2

| | section 1 | section 2 | section 3 | section 4 | section 5 | section 6 | section 7 | section 8 | section 9 | section 10 |
|---|---|---|---|---|---|---|---|---|---|---|
| *alg−5* | −1.63 | −1.27 | −0.85 | −0.34 | 0.71 | 0.71 | 0.68 | 1.39 | −0.08 | 0.69 |
| *prg−1* | −0.42 | 1.26 | 1.52 | 1.09 | −0.24 | −0.94 | −0.76 | 0.22 | −0.29 | −1.45 |
| *csr−1* | 0.30 | 0.92 | 1.17 | 1.49 | 0.55 | −0.51 | −0.93 | −0.88 | −0.96 | −1.15 |
| *wago−1* | −1.56 | −0.74 | −0.69 | −0.41 | 0.32 | −0.28 | −0.46 | 1.57 | 1.23 | 1.02 |
| *ppw−2* | 1.75 | 1.42 | 0.69 | 0.37 | −0.19 | −0.87 | −0.67 | −0.78 | −0.88 | −0.84 |
| *wago−4* | 0.18 | 2.27 | 1.15 | 0.09 | −0.40 | −0.61 | −0.92 | −0.44 | −0.55 | −0.78 |
| *ppw−1* | 1.25 | 1.67 | 0.16 | −0.22 | 0.96 | −0.54 | −0.80 | −1.45 | −0.73 | −0.29 |
| *hrde−1* | −1.75 | −1.42 | 0.20 | 0.27 | 0.59 | −0.17 | −0.67 | 1.11 | 0.77 | 1.08 |
| *alg−3* | | | | | | | | | | |
| *alg−4* | | | | | | | | | | |
| *wago−10* | | | | | | | | | | |
| *alg−2* | −1.05 | −0.86 | −0.93 | −0.95 | −0.75 | 0.32 | 1.35 | 1.04 | 0.62 | 1.20 |
| *ergo−1* | −1.49 | −0.94 | −1.02 | −0.72 | −0.02 | 0.40 | 0.54 | 1.56 | 0.95 | 0.74 |
| *vsra−1* | −1.02 | −0.98 | −0.84 | −1.02 | −0.42 | 0.37 | 1.50 | 0.23 | 0.73 | 1.45 |
| *nrde−3* | −0.94 | −0.94 | −0.81 | −0.91 | −0.87 | 0.43 | 0.81 | 1.28 | 1.51 | 0.44 |

Figure S15

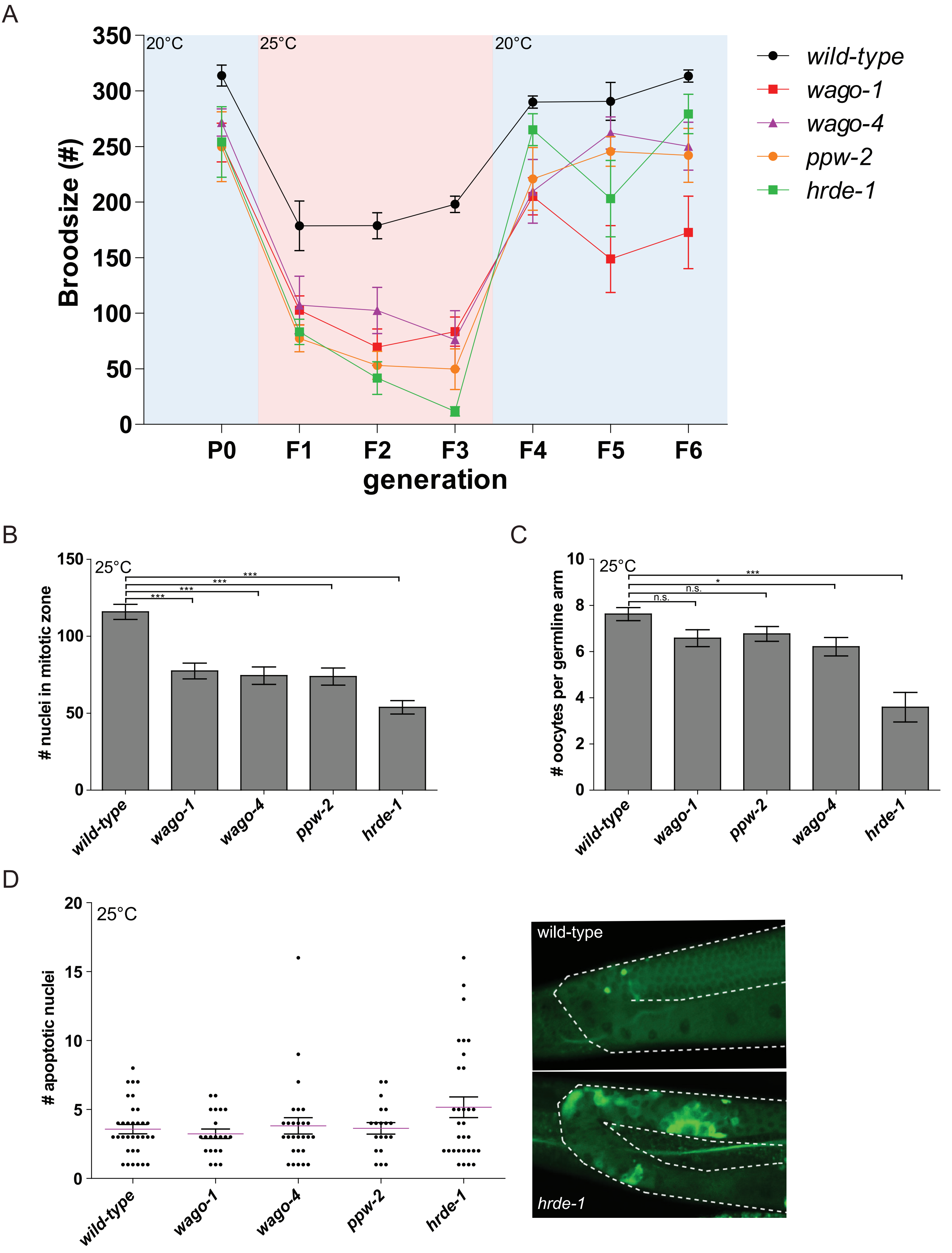
A
20°C
25°C
20°C
Broodsize (#)
350
300
250
200
150
100
50
0
P0
F1
F2
F3
F4
F5
F6
generation
wild-type
wago-1
wago-4
ppw-2
hrde-1
B
25°C
### nuclei in mitotic zone
150
100
50
0
***
***
***
***
wild-type
wago-1
wago-4
ppw-2
hrde-1
C
25°C
### oocytes per germline arm
10
8
6
4
2
0
***
*
n.s.
n.s.
wild-type
wago-1
ppw-2
wago-4
hrde-1
D
25°C
### apoptotic nuclei
20
15
10
5
0
wild-type
wago-1
wago-4
ppw-2
hrde-1
wild-type
hrde-1
